# Interdependence between histone marks and steps in Pol II transcription

**DOI:** 10.1101/2020.04.08.032730

**Authors:** Zhong Wang, Alexandra G. Chivu, Lauren A. Choate, Edward J. Rice, Donald C. Miller, Tinyi Chu, Shao-Pei Chou, Nicole B. Kingsley, Jessica L. Petersen, Carrie J. Finno, Rebecca R. Bellone, Douglas F. Antczak, John T. Lis, Charles G. Danko

## Abstract

The role of histone modifications in transcription remains incompletely understood. Here we used experimental perturbations combined with sensitive machine learning tools that infer the distribution of histone marks using maps of nascent transcription. Transcription predicted the variation in active histone marks and complex chromatin states, like bivalent promoters, down to single-nucleosome resolution and at an accuracy that rivaled the correspondence between independent ChIP-seq experiments. Blocking transcription rapidly removed two punctate marks, H3K4me3 and H3K27ac, from chromatin indicating that transcription is required for active histone modifications. Transcription was also required for maintenance of H3K27me3 consistent with a role for RNA in recruiting PRC2. A subset of DNase-I hypersensitive sites were refractory to prediction, precluding models where transcription initiates pervasively at any open chromatin. Our results, in combination with past literature, support a model in which active histone modifications serve a supportive, rather than a regulatory, role in transcription.

## Introduction

The discovery that core histones are post-transcriptionally modified fueled nearly six decades of speculation about the role that histone modifications play in transcriptional regulation by RNA polymerase II (Pol II)^1^. Many of the best-studied histone modifications are deeply conserved within eukaryotes, indicating important functional roles^2–4^. Indeed numerous examples illustrate how the disruption of histone modifications, or their associated writer and eraser enzymes, lead to defects in transcription and cellular phenotypes^5–8^. Histone modifications are found in highly stereotyped patterns across functional elements, including promoters, enhancers, and over the body of transcribed genes and non-coding RNAs. Promoters and enhancers are associated with a pattern of chromatin organization consisting of a nucleosome depleted core flanked by +1 and -1 nucleosomes marked with specific histone modifications, including histone 3 lysine 4 trimethylation (H3K4me3), lysine 27 acetylation (H3K27ac), and lysine 4 monomethylation (H3K4me1)^9–15^. Actively transcribed gene bodies are marked by histone 3 lysine 36 trimethylation (H3K36me3) and histone 4 lysine 20 monomethylation (H4K20me1)^9, 16^. Finally, two modifications are enriched in transcriptionally depleted regions, including histone 3 lysine 27 trimethylation (H3K27me3) and lysine 9 trimethylation (H3K9me3)^16^.

The stereotyped pattern of histone modifications makes them useful in the annotation of functional elements in eukaryotic genomes. A collection of 11 histone modifications, used to broadly analyze different cell types by the ENCODE project, has been applied to identify functional elements in metazoans^2, 16–18^. Numerous studies have used histone modifications to reveal the location of enhancers, lincRNAs, and other types of functional elements^10, 19–22^. Histone modifications aid in interpreting phenotype-associated genetic variation^23, 24^ and discovering molecular changes in disease^25–30^. Likewise, histone modifications have been proposed for applications in selecting individualized therapeutic strategies^31^. Applications such as these have led to genome annotation efforts in a myriad of mammals^32^, plants^33–35^, and other eukaryotic organisms^36^. New annotation efforts will be launched alongside ‘moonshot goals’ to sequence and annotate genomes across the tree of life^37^. However, despite extensive efforts to decrease cost and improve the throughput of experimental methods^38–42^, and to “impute” (or guess) the abundance of marks that were not directly observed^43, 44^, genome annotation still requires concerted efforts of large, well-funded, interdisciplinary consortia.

Despite the widespread use of histone modifications in genome annotation, the precise nature of the relationship between histone modifications and transcription remains unknown. Specific histone modifications are enriched in either transcriptionally active or quiescent regions^9, 45–48^. However, the extent to which histone modifications have a direct role in transcriptional regulation or an indirect role as “cogs” in the transcription machinery, remains debated^49^. Certain combinations of histone modifications, most notably the bivalent chromatin signature consisting of H3K4me3 and H3K27me3, are speculated to mark specific genes for transcriptional activation in later developmental stages^50^. In another example, the balance between H3K4me1 and H3K4me3, which has long been known to correlate with enhancer and promoter activity^10^, has been proposed to establish these two regulatory roles^51^. Another question that remains heavily debated is the extent to which distinct histone modifications mark DNA sequence elements that otherwise have similar functional activities. H3K27ac, H3K64ac, and H3K122ac are all reported to denote distinct sets of enhancers^52^. Finally, to what extent do histone modifications cause transcription? The nature of the quantitative relationship between transcription and histone modification lies at the crux of this open question. Under one model, marks serve as a placeholder which might contribute to transcriptional regulation in a distinct cellular state, in which case we expect large amounts of histone modifications that are not explained by current transcription events. Alternatively, if histone modifications serve as “cogs” that have a critical role in transcription, but do not themselves have a regulatory role independent of transcription factors, we might expect that they are completely correlated with on-going transcription.

Here we trained sensitive machine learning models that decompose maps of primary transcription into ChIP-seq profiles representing 9 distinct histone modifications. We show that transcription measured using precision run-on and sequencing (PRO-seq)^45, 53, 54^ can recover the pattern of active histone modifications at nucleosome resolution and with an accuracy that rivals the correlation between independent ChIP-seq experiments in holdout cell types. To define the nature of the causal relationship between transcription and histone modifications, we perturbed transcription and examined the genomic distribution of four active and one repressive histone modification: H3K4me1, H3K4me3, H3K27ac, H3K36me3, H3K27me3. Surprisingly, transcription was critical in the deposition of promoter associated histone modifications, H3K4me3 and H3K27ac. Although transcription accurately predicted nearly all histone modifications and also open chromatin structure, we found a subset of DNase-I hypersensitive sites that were refractory to prediction. Collectively, our results (1) support models in which histone modifications are “cogs” with a supportive role, rather than a direct regulatory role, in transcription, (2) preclude models in which transcription initiates pervasively as a consequence of open chromatin, and (3) provide a new strategy for genome annotation using a single functional assay that is tractable for an individual lab to perform.

## Results

### Accurate imputation of histone modifications at nucleosome resolution using nascent transcription

To better understand the nature of the relationship between transcription and histone modifications, we trained <u>d</u>iscriminative <u>h</u>istone imputation using transcription (dHIT). dHIT uses the distribution of RNA polymerase, measured using any of the related run-on and sequencing methodologies PRO-seq, GRO-seq, or ChRO-seq (henceforth referred to simply as PRO-seq), to impute the level of histone modifications genome-wide. dHIT passes transformed PRO-seq read counts in windows of various sizes to a support vector regression (SVR) (see Methods). Run-on assays provide a readout of the position and density of RNA polymerase with single nucleotide resolution, which the SVR uses to impute information about chromatin structure and marks. During a training phase, the SVR optimized a function that mapped PRO-seq signal to the quantity of ChIP-seq signal at each position of the genome (**Fig. 1a**; see Methods). Once a dHIT model was trained using existing ChIP-seq data, it can impute steady state histone modifications in any cell type, provided that the relationship between histone modification and transcription is preserved. We trained dHIT to impute the levels of 10 different histone modifications that are widely deployed to analyze chromatin state (**Fig. 1a**)^16, 17, 55, 56^. To avoid overfitting to batch-specific features in a single run-on and sequencing dataset^56^, training was performed using seven datasets in K562 cells that exemplify the range of variation commonly observed between data in library quality, sequencing depth, run-on strategy (PRO-seq or GRO-seq), and pausing index (**Supplementary Table 1; Supplementary Table 3**).

**Figure 1.**
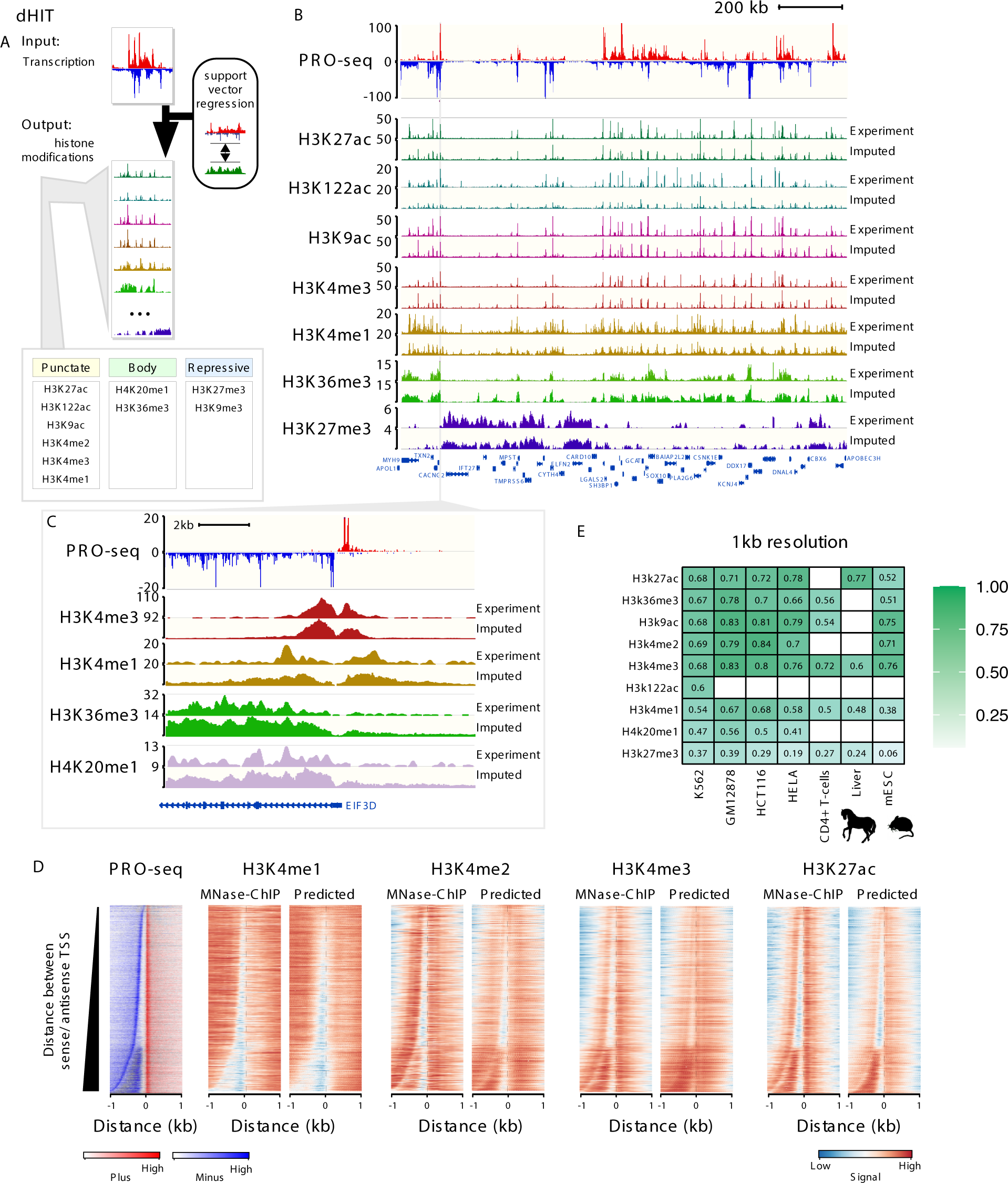
dHIT imputes his tone modifications us ing nas cent transcription. (A) Schematic of the dHIT algorithm. PRO-seq and ChIP-seq data in K 562 cells was used to train a support vector regression (SVR) classifier to impute 10 different his tone modifications. (B) Genome-browser compares experimental and predicted his tone modifications on a holdout chromosome (chr22). PRO-seq data used to generate each imputation is shown on top. (C) Genome-browser compares experimental and predicted his tone marks near the promoter of EIF 3D. PRO-seq data used to generate each imputation is shown on top. (D) Heatmaps show the distribution of transcription (left) and his tone modifications (right) measured using MNase ChIP-seq or predicted using transcription. Rows represent transcription initiation domains in K 562 cells. Heatmaps were ordered by the distance between the most frequently used TSS in each transcription initiation domain on the plus and minus strand. (E) Pears on’s correlation between predicted and expected values for nine his tone modifications. Values are computed on the holdout chromosome (chr22) in humans, chr1 in horse, and chr1 in mice. Empty cells indicate that no experimental data is available for comparis on in the cell type shown.

We evaluated the accuracy of each dHIT imputation model on a holdout chromosome in one of the training datasets (chr22; **Fig. 1b-c; Supplementary Fig. 1; Supplementary Fig. 2**). Histone modification signal intensity imputed using dHIT was highly correlated with experimental data for a variety of marks with different genomic distributions, including marks with focused signal on promoter or enhancer regions (e.g., H3K4me1/2/3, H3K27ac, H3K9ac), marks spread across active gene bodies (H3K36me3, H4K20me1), and over large domains of PRC2-dependent repressive heterochromatin (H3K27me3). The most notable differences between imputed and experimental signals tended to be small differences in background regions with low intensity in both experimental and imputed signal, but which added up over large windows, reflecting technical sources of variation in ChIP-seq background signal that were not reflected in PRO-seq signal (**Supplementary Fig. 1; Supplementary Note 1**). We did not observe major differences in accuracy at different types of functional elements, including regions of high signal intensity in either experimental or imputed data, near gene promoters^44, 57^, at distal enhancer elements, and at stable and unstable transcription start sites^11^ (**Supplementary Fig. 2**). In addition to well-studied and commonly used histone marks, we also obtained a high degree of correspondence for less widely studied histone modifications. For instance, acetylation of lysine 122 (H3K122ac), a residue on the lateral surface of the H3 globular domain^58^, was reported to mark a distinct set of enhancers compared with H3K27ac^52^. Nevertheless, dHIT models trained to impute H3K122ac had a high correlation on the holdout chromosome (**Fig. 1b**). Of the marks for which we attempted to train models, only the repressive mark H3K9me3 did not perform well against either ENCODE data, or against higher-quality CUT&RUN data^59^ (**Supplementary Fig. 3**).

In many cases, imputation captured the fine-scale distribution of histone mark signals near the transcription start site (TSS) of annotated genes or enhancers (**Fig. 1c; Supplementary Fig. 4; Supplementary** **Fig 5**). To explore the limit of the resolution for histone mark imputation using transcription, we obtained new ChIP-seq data for four active marks whose distribution correlates with enhancers and promoters (H3K4me1, H3K4me2, H3K4me3, and H3K27ac) at nucleosome resolution by using MNase to fragment DNA. We also analyzed the gene body mark H3K36me3 that is depleted near the TSS^9^. We trained new SVR models in K562 cells that take advantage of the higher-resolution MNase ChIP-seq data, excluding chromosome 22 as a holdout to confirm a high correlation (**Supplementary Fig. 6**). Examination of genome-browser traces near the TSS of genes on the holdout chromosome confirmed that dHIT could impute active marks with high resolution (**Supplementary Fig. 7**).

Genome-wide, several aspects of chromatin organization were correlated with the precise location of TSSs and Pol II pause sites. These features are readily apparent when sorting by the distance between the strongest TSS on the plus and minus strand^13–15^ (**Fig. 1d**). First, when the distance between the maximal sense and divergent TSS was larger than ∼300 bp, we observed a nucleosome between the divergent start sites that was marked predominantly with H3K4me3 and H3K27ac but depleted for H3K4me1. Second, H3K4me3 and H3K27ac signals were highest on the +1 nucleosome, as well as the nucleosome found inside of the initiation domain. Third, H3K4me2 was highest on the -1 nucleosome. Fourth, the gene body mark, H3K36me3, was depleted at the promoter, and enriched in the body of transcribed genes (**Supplementary Fig. 8**). Each of these correlations between TSSs and chromatin marks were also observed to varying degrees in genome-wide imputation in K562 cells (**Fig. 1d**), and imputation data in a complete holdout cell type, GM12878 (**Supplementary Fig. 9**). Thus, dHIT recovered the placement of nucleosomes constrained to ordered arrays whose position correlated with transcription initiation.

### Active histone modifications have a similar relationship to transcription across mammalian cells

We asked whether the relationship between transcription and histone modifications is a general feature that is shared across mammalian cell types. We computed the correlation between imputed and experimental histone marks in five holdout datasets without retraining the model. Holdout datasets were selected to represent a range of cultured cells, primary cells, and tissues from multiple mammalian species (**Supplementary Table 1**). Holdout data also explored a range of technical variation in both run-on assays and ChIP-seq validation experiments, including data collected by different labs, different fragmentation methods, and, for the run-on experiment, using different variants of a run-on assay (**Supplementary Table 1-2**).

Despite a variety of technical differences between ChIP-seq in holdout cell types and the ENCODE training dataset (**Supplementary Table 2**), active marks were recovered with a similar fidelity in holdout cell types as observed for K562 (**Fig 1e, Supplementary Fig. 10a-c**). At 1kb resolution, dHIT recovered active marks indicative of promoters, enhancers, and gene-bodies at a median Pearson correlation of 0.73 (Pearson’s R = 0.38-0.84), substantially higher than copying values from the training dataset (**Supplementary Fig. 10d**). Lower correlations were generally observed when the experimental ChIP-seq data (certain CD4+ T-cell datasets) or the PRO-seq data (e.g., HeLa) had fewer sequenced reads or lower values in other data quality metrics (**Supplementary Table 3**). For marks that were distributed across broad genomic regions (H3K36me3 and H3K27me3), dHIT imputation identified broad regions of high signal with reasonably high accuracy but smoothed over fine-scale variation (**Fig. 1e; Supplementary Fig. 1**). Finally, cell-type specific signal differences were predicted with reasonably high accuracy (Pearson’s R = 0.44-0.70 for active marks; **Supplementary Fig. 10f**), providing additional confidence that dHIT was not simply learning the average signal intensity of histone modification^60^. Thus, dHIT accurately recovered the distribution of active histone marks in a way that generalized to all new cell types examined here.

To place correlations observed between imputed and experimental data into context, we compared correlations between imputed and ChIP-seq data to those observed between different experimental datasets in K562 and GM12878, two cell lines for which multiple data exists for each mark. For active marks, and for H3K27me3, correlations between dHIT imputation and experimental data were often within the range observed between experimental datasets (**Supplementary Fig. 11**). In addition to signal intensity, imputation could also recover the location of ENCODE peak calls in GM12878 with an accuracy rivaling ChIP-seq experiments (**Supplementary Fig. 2a-b**). These data indicate that imputation achieved performance similar to ChIP-seq experimental replication for most marks, with the notable exception of H3K9me3.

We examined specific loci in which imputed histone marks differed substantially from experimental data. Differences could reflect either cases in which histone modifications deviate from transcription for a specific mechanistic reason, or they may reflect biological differences between cell stocks reflecting genetic factors, growth conditions, handling, or other confounding factors. To distinguish between these possibilities, we repeated ChIP-seq for H3K27ac in K562 cells that were closely matched with those used to prepare PRO-seq libraries. In nearly all cases, our own ChIP-seq data resolved major discrepancies between imputed and ENCODE datasets (**Supplementary Fig. 12**). We therefore concluded that major discrepancies between imputed and experimental marks predominantly reflect intrinsic biological or experimental/technical differences, rather than divergence between transcription and histone modifications.

**Figure 2.**
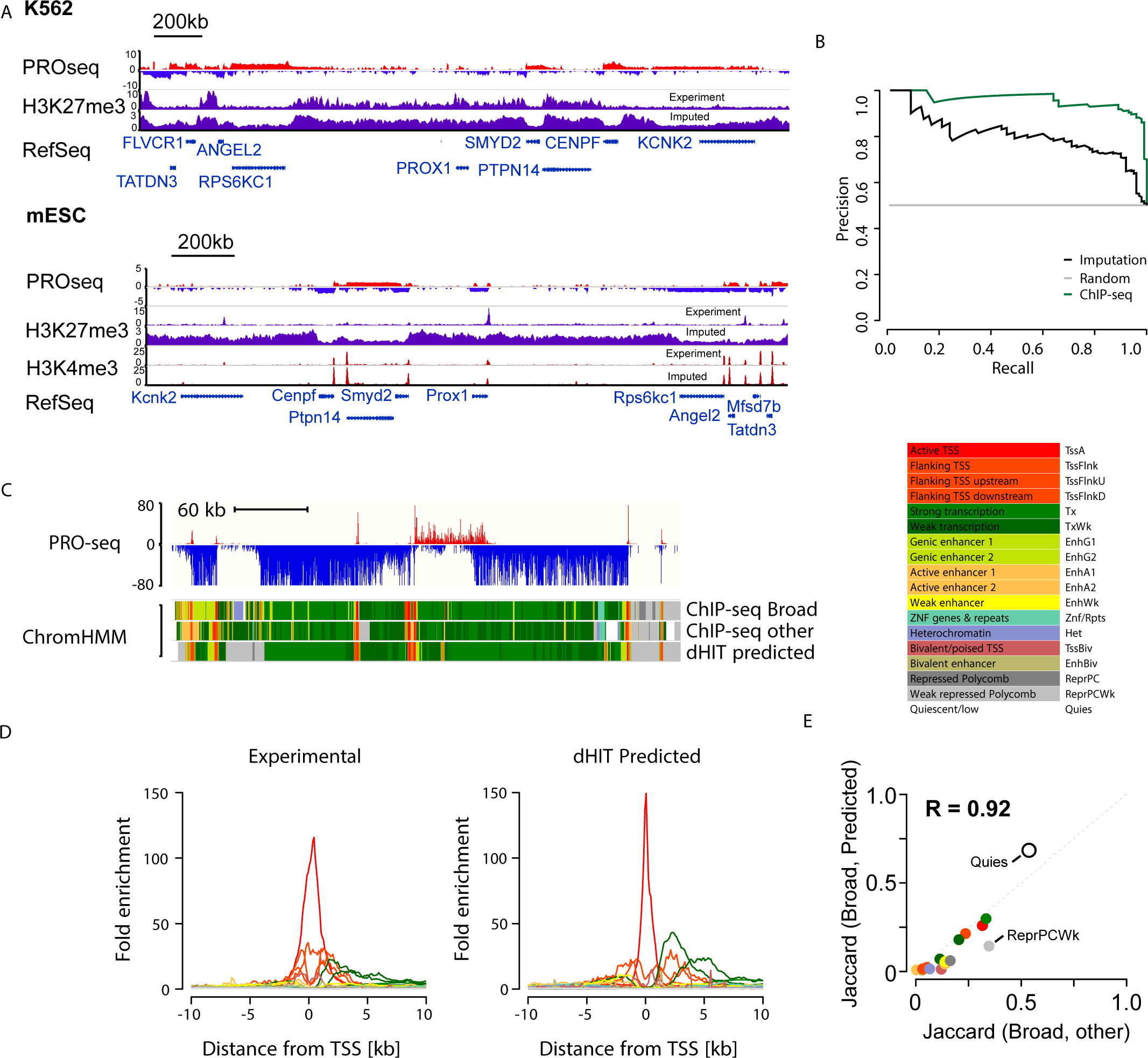
dHIT identifies bivalent H3K4me3, H3K27me3 marked genes. **(A)** Genome-browser shows PRO-seq data and histone modification data measured by ChIP-seq or predicted using PRO-seq in the Prox1 locus. Prox1 is marked by bivalent H3K4me3 and H3K27me3 histone modifications in mESCs. **(B)** Precision recall curve illustrates the accuracy of bivalent gene classification by a random forest classifier using ChIP-seq data (green) or dHIT imputation (black). The gray line denotes random classification. Classification was performed on a matched set of TSSs (50% bivalent, 50% not bivalent) that was held out during random forest training. **(C)** Genome-browser in K562 cells shows 18 state chromHMM model using either ChIP-seq data used to train the model (Broad), alternative ChIP-seq data in K562 (other), or based on imputation (dHIT predicted). PRO-seq data used during dHIT imputation is shown on top. **(D)** Enrichment in each of 18 chromatin states as a function of distance from RefSeq annotated transcription start sites. **(E)** Jaccard distance between chromHMM states inferred using ChIP-seq from Broad and predicted data (y-axis) and states inferred using ChIP-seq from Broad and an alternative compilation of high-quality ChIP-seq data (x-axis).

### Two separate patterns of H3K27me3 reflect stem- and differentiated-cell states

We identified one important exception on the extent to which histone imputation generalized between cell types. The repressive mark, H3K27me3, had a reasonable correlation with experimental data in K562, GM12878, and horse liver (median Pearson’s R = 0.31), consistent with the correlation expected from biological replication in K562. In these cell types, H3K27me3 was distributed across broad genomic intervals, which were identified with reasonable fidelity by dHIT imputation (**Fig. 2A, top**). However, we observed a much weaker correlation in mouse embryonic stem cells (mESCs, Pearson’s R = 0.06). Examination of signal tracks showed that the distribution of H3K27me3 differed dramatically from the K562 cell dataset. In mESCs, H3K27me3 was predominantly positioned in punctate peaks near weakly transcribed promoters (**Fig. 2A, bottom**). Although a handful of loci with critical developmental importance, notably all four HOX clusters, had a broad distribution in the mESC data, these did not show the pattern expected in the mark based on transcription (**Supplemental Fig. 13**). Analysis of H3K27me3 in 86 high-quality samples showed that stem, germ, and certain progenitor cells usually had a punctate pattern, whereas most somatic cell types had the broadly distributed pattern (**Supplemental Fig. 14B-C**). Thus, although we cannot completely discount the possibility that technical factors contribute to this difference in H3K27me3 distribution^61–63^, both punctate and broad H3K27me3 distributions appear even when libraries were prepared by the same lab^64^ or consortium^16, 17^. These observations suggest that H3K27me3 can occur in at least two distinct profiles, and that transcription is able to predict the broadly distributed profile found in somatic cell types with reasonable accuracy.

### Imputation of bivalent promoters and other chromatin states

We next asked whether dHIT could impute complex chromatin states consisting of multiple histone marks. The bivalent chromatin state in mESC is a perfect example where nucleosomes near gene promoters are marked with both H3K4me3 and H3K27me3, which are associated with gene activation and repression, respectively^50^. The bivalent chromatin state is best described in ESCs and germ cells, and tends to mark the promoter of genes with developmental importance^50, 65–67^. We used dHIT models trained on ENCODE ChIP-seq data in K562 cells to impute H3K4me3 and H3K27me3 based on a GRO-seq dataset in mESCs^68^. Despite cell-type-specific differences in the relationship between transcription and H3K27me3 between K562 and mESCs (noted above), we observed a strong tendency for bivalent promoters in mESCs to fall inside broad domains that dHIT predicted to have high H3K27me3. For example, the K562 model predicts that *Prox1* resides inside of a broad H3K27me3 domain, based on low transcription levels from *Prox1* and surrounding regions (**Fig. 2A**). Despite being far from highly transcribed genes, the *Prox1* promoter is weakly transcribed, and the imputation correctly places a peak of H3K4me3. The general pattern where bivalent genes were transcribed within H3K27me3-high domains was consistent enough that nearly 80% of bivalent gene promoters could be separated from promoters associated with either mark alone, or neither mark, with a precision of 80%, using a random forest on holdout data (**Fig. 2B**). Notably, promoters that carry the H3K27me3 mark in mESCs were distinguished accurately from those carrying no mark, indicating that promoters carrying the H3K27me3 mark are generally not transcriptionally silent. Taken together, these results demonstrate that bivalent genes can be identified based on the distribution of active transcription alone.

To generalize our observations on bivalent genes to other chromatin states, we asked whether chromatin marks imputed using transcription can infer chromatin states defined by chromHMM^69^. We used a previously reported chromHMM model that defined 18 distinct chromatin states using ChIP-seq data from six marks for which we trained imputation models (H3K4me3, H3K27ac, H3K4me1, H3K36me3, H3K9me3, and H3K27me3)^17, 70^. Examination on the WashU epigenome browser revealed that chromatin states were highly similar, regardless of whether they were defined using experimental data from ENCODE or dHIT imputation (**Fig. 2C**). Patterns of chromHMM state enriched near the TSS of annotated genes were similar between experimental or dHIT imputed data as input (**Fig 2D, Supplementary Fig. 15**). To determine the concordance expected between chromatin states defined using independent collections of experimental data, we applied chromHMM to a distinct collection of ChIP-seq data in the same cell type (**Supplementary Table 1**). The Jaccard similarity index between imputed and experimental data were highly correlated with those observed between other ChIP-seq datasets (Pearson’s R = 0.92; **Fig. 2E, Supplementary Fig. 16**). Taken together, these results suggest that transcription alone is sufficient to infer complex chromatin states, especially active chromatin states.

### Genome annotation using a single functional assay

Histone modifications are widely used to annotate mammalian genomes. We hypothesized that since dHIT can accurately predict chromatin marks, it provides a strategy for genome annotation in limited samples or new mammalian species using a single molecular tool. We analyzed chromatin states in 20 primary glioblastomas (GBMs) for which we recently published ChRO-seq data^54^. ChromHMM analysis revealed both broad similarities and putative differences in chromatin states between different GBMs. For instance, a subset of samples was characterized by active transcription in *ADM2* and *MIOX* (**Fig. 3A**). Analysis of histone modifications of this same set of samples would have taken 120 separate ChIP-seq experiments (**Fig. 3B**) and require sequencing of each experiment to a depth ∼4 times greater than ChRO-seq using ENCODE guidelines. Thus, ChRO-seq and dHIT can resolve intricate patterns of chromatin organization using a single molecular assay.

**Figure 3.**
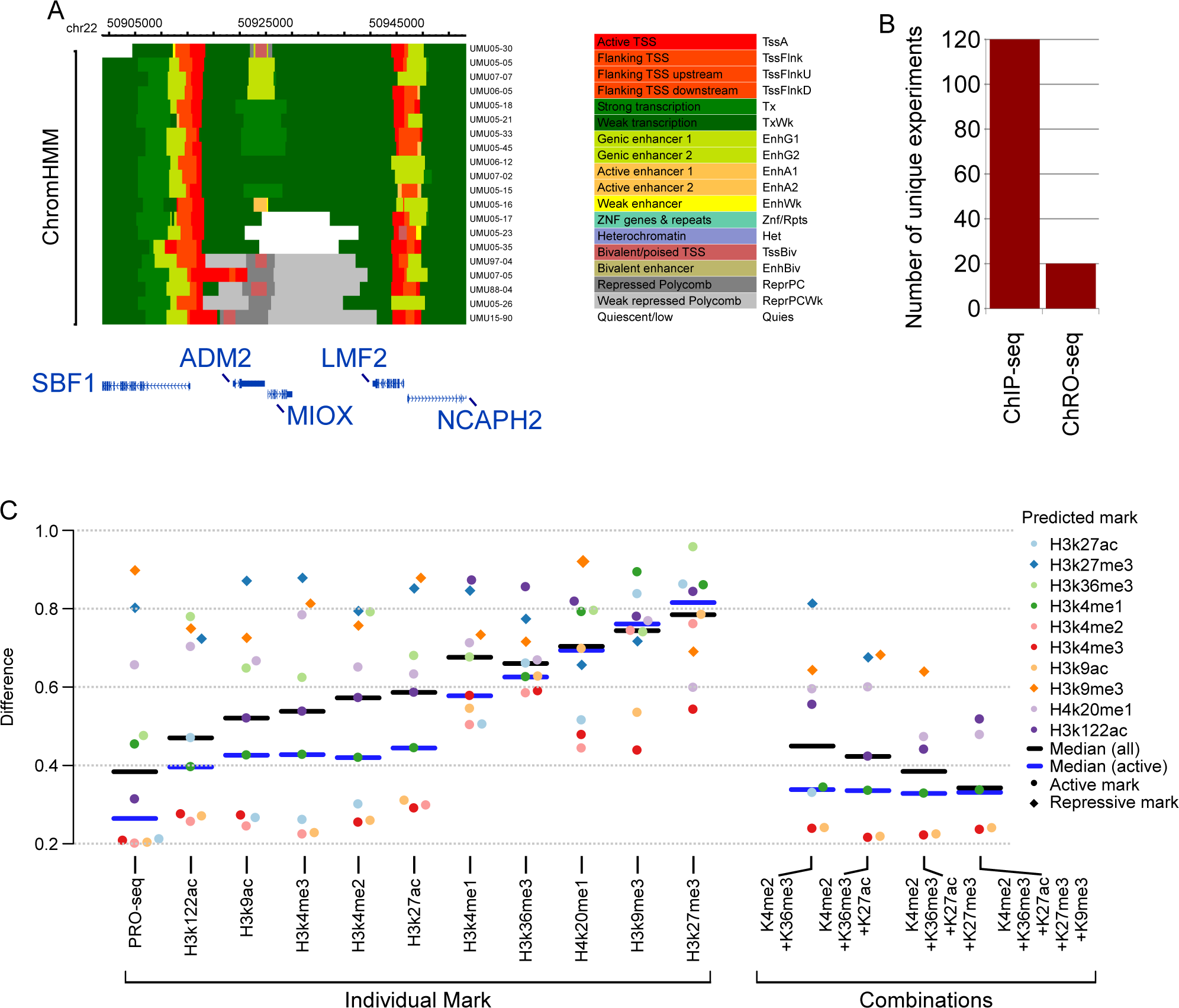
Inference of chromatin states defined by chromHMM using transcription. (A) ChromHMM states inferred using ChRO-seq data from 20 primary glioblastomas. (B) The number of unique ChRO-seq or ChIP-seq libraries required to analyze chromatin states in 20 primary glioblastomas. (C) The mean difference between predicted and experimental ChIP-seq data on a holdout chromosome (chr22) (Y-axis). SVR models were trained using the indicated experimental mark (left) or the indicated combination of histone marks (right).

Another critical application is to efficiently annotate functional elements in diverse tissues from understudied species. We obtained ChRO-seq data from the liver of two horses that serve as the focus of the Functional Annotation of Animal Genomes (FAANG) project^32, 71, 72^. Using dHIT and models trained in K562 cells to impute histone modifications, we obtained patterns of H3K27ac, H3K4me3, H3K4me1, and H3K27me3 that were highly correlated with experimental data from the same tissues (**Fig. 1E; Supplementary Fig. 17A**). In addition to those histone marks measured by FAANG, dHIT also imputed patterns for five additional histone marks, providing new information about chromatin state that was not obtained by the FAANG consortium. Next, we prepared ChRO-seq libraries in eight murine tissues (**Supplementary Fig. 17B; Supplementary Fig. 18**). After accounting for biological replication in this experiment^73^ (7 replicates x 8 tissues x 9 histone marks), it would have taken 504 ChIP-seq assays to prepare this same dataset. Thus, using dHIT to interpret ChRO-seq data provides individual labs access to consortium scale annotation of functional elements in mammalian genomes, and this information has potential applications in precision diagnostic medicine and genome annotation.

To further assess the relative power of PRO-seq and dHIT for predicting unobserved histone modifications, we asked whether PRO-seq more accurately predicted unobserved histone modifications than SVR models trained using a small number of observed histone modifications. To identify the best assay for this task, we trained SVR imputation models that use either PRO-seq or ChIP-seq data for each of the 10 different histone marks to predict each of the other experimental ChIP-seq datasets. We evaluated performance using the L1 norm, defined as the average of the median centered distance between imputed and experimental marks in 10bp windows on a holdout chromosome (**see Methods**). PRO-seq achieved a lower median L1 norm than any other individual assay by a fairly wide margin (**Fig. 3C, black**). Examining imputation tracks led us to attribute the relative success of PRO-seq to two features. First, PRO-seq captured the boundaries and direction of gene bodies in a manner that could not be achieved by other marks (**Supplementary Fig. 19A**). Second, PRO-seq was the most accurate at recovering the relative distribution of signal intensities in focal marks near the TSS (**Supplementary Fig. 19B**). Thus, we conclude that PRO-seq improved the accuracy of histone mark imputation by encoding signals from multiple functional regions and by improving spatial resolution compared with ChIP-seq data.

We next trained SVRs using combinations of multiple histone marks to determine whether training on multiple experimental datasets improved imputation performance. Because the space of potential histone mark combinations was extremely large and training was time consuming, we manually selected combinations of histone marks that provide orthogonal information to each other. We first selected H3K4me2 and H3K36me3, which combined a mark denoting promoter/ enhancer regions with one denoting gene bodies^9, 74^. The pair of experimental datasets together slightly improved the imputation of most ChIP-seq marks relative to the best performing individual mark, for instance H3K4me1 and H3K9me3 (**Fig. 3**). However, the median L1 norm was still worse than PRO-seq. We tested combinations where larger numbers of marks were observed by adding H3K27ac, H3K27me3, and H3K9me3, and evaluating the accuracy with which imputation could recover experimental marks. In most cases using additional marks made only a minor difference in performance (**Fig. 3**). Although we observed a decrease in the median accuracy using multiple marks (**Fig. 3** **black**), this was explained largely by replacing the worst performing marks with experimental data. Our results therefore suggest that capturing information about the relative position of TSSs and gene bodies was enough to saturate performance using our current framework. Thus, PRO-seq data predicted ChIP-seq signals of unobserved active histone marks at least as well as ChIP-seq data for five different histone marks.

### Transcription is required for promoter-associated histone modification

The strong correlation observed between Pol II and histone modifications fit a model in which there is a casual relationship between the histone marks and transcriptional activity. However, correlations do not provide insight into which direction causality might run. To assess whether Pol II activity is necessary, and thus causal, for histone modifications, we rapidly blocked transcription initiation using the small molecule Trp and observed the immediate effects on both transcription (using PRO-seq) and histone modifications (using MNase-ChIP-seq) (**Fig. 4A**)^75, 76^. After spike-in normalization, PRO-seq revealed the expected pattern of changes in Pol II throughout the time-course^68^: a large loss in Pol II near active transcription start sites by 1h of treatment, followed by an almost complete loss in signal across the entire genome by 4h (**Fig. 4C, right; Supplementary Fig. 20**). As Trp does not affect engaged RNA polymerase, we observed a clearing wave of Pol II ∼100kb from the TSSs on long genes at 1h (**Fig. 4C, left**), consistent with reported elongation rates of ∼1-3kb per minute^68, 77^.

**Figure 4.**
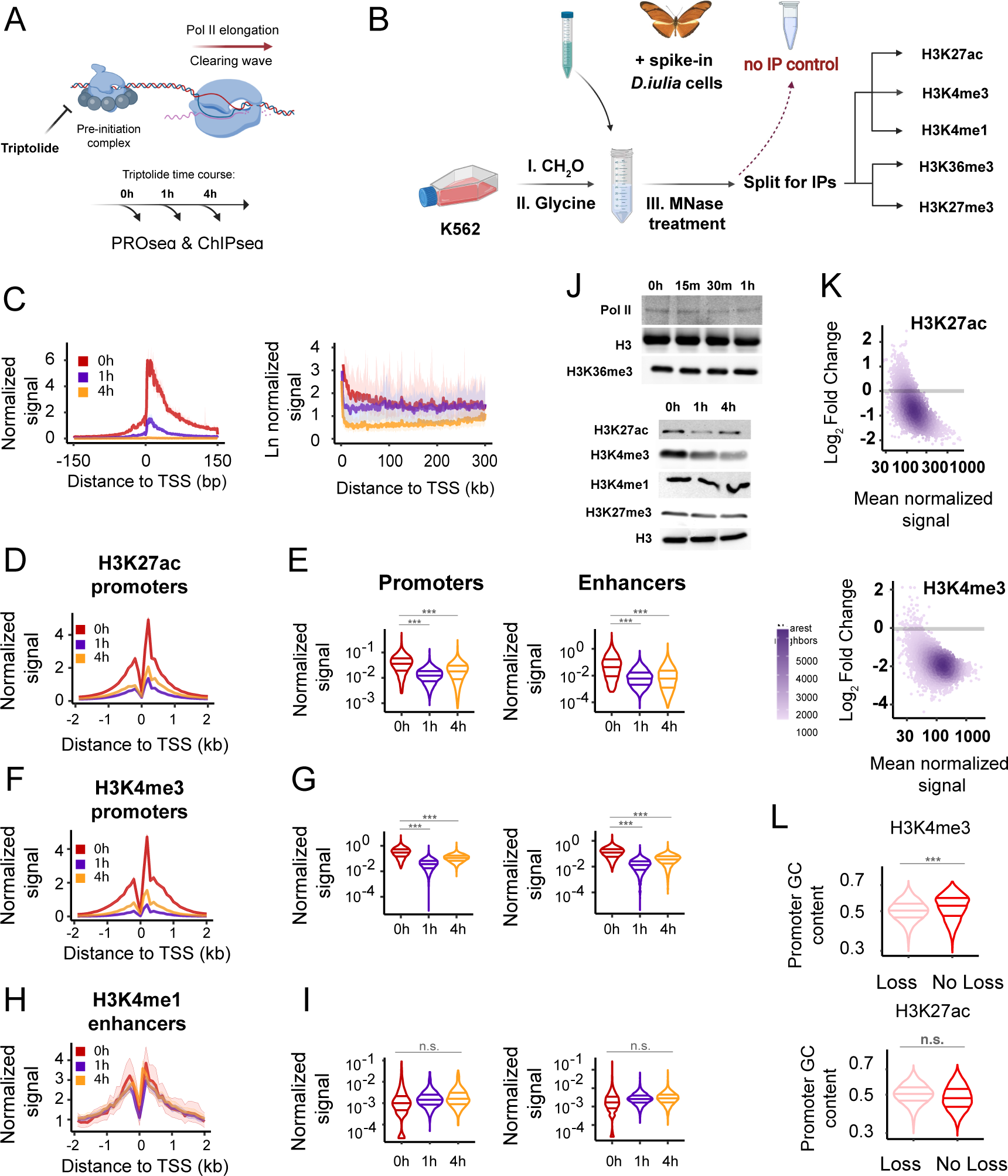
ChIP-seq measures changes in histone modifications following transcription inhibition by Trp. (A) Model of Trp action on transcription pre-initiation complex. (B) Metaplots of PRO-seq signal after Trp treatment. Pol II density is depicted on a linear scale in a 300bp window centered on maximum TSS (left), or on a natural log scale (right). (C) Depiction of ChIP-seq experimental design where D. iulia chromatin was used as spike-in normalization control. See also STAR Methods. (D-I) Meta plots and quantification of H3K27ac (D-E), H3K4me3 (F-G), and H3K4me1 (H-I) signals at enhancers and gene promoters. A paired, two-sided, Wilcoxon test was performed to estimate statistical significance in signal changes, where (***) denote p-value < 2.2e-16 and (n.s.) p-value =1. (J) Western blots show global changes in histone marks after Trp treatment. Each blot depicts chromatin associated histone marks and Pol II after the indicated Trp incubation time. See also Supplementary Fig. 20. (K) MA plots display the loss in H3K4me3 and H3K27ac between 0h and 1h of Trp treatment. Log2 fold changes and mean normalized signals between time points were computed with DEseq2. A gray bar marks log2 fold change at 0. (L) Violin plots quantify the levels of H3K4me3 and H3K27ac as a function of GC-richness of promoter sequences. Statistical significance was

**Figure 5.**
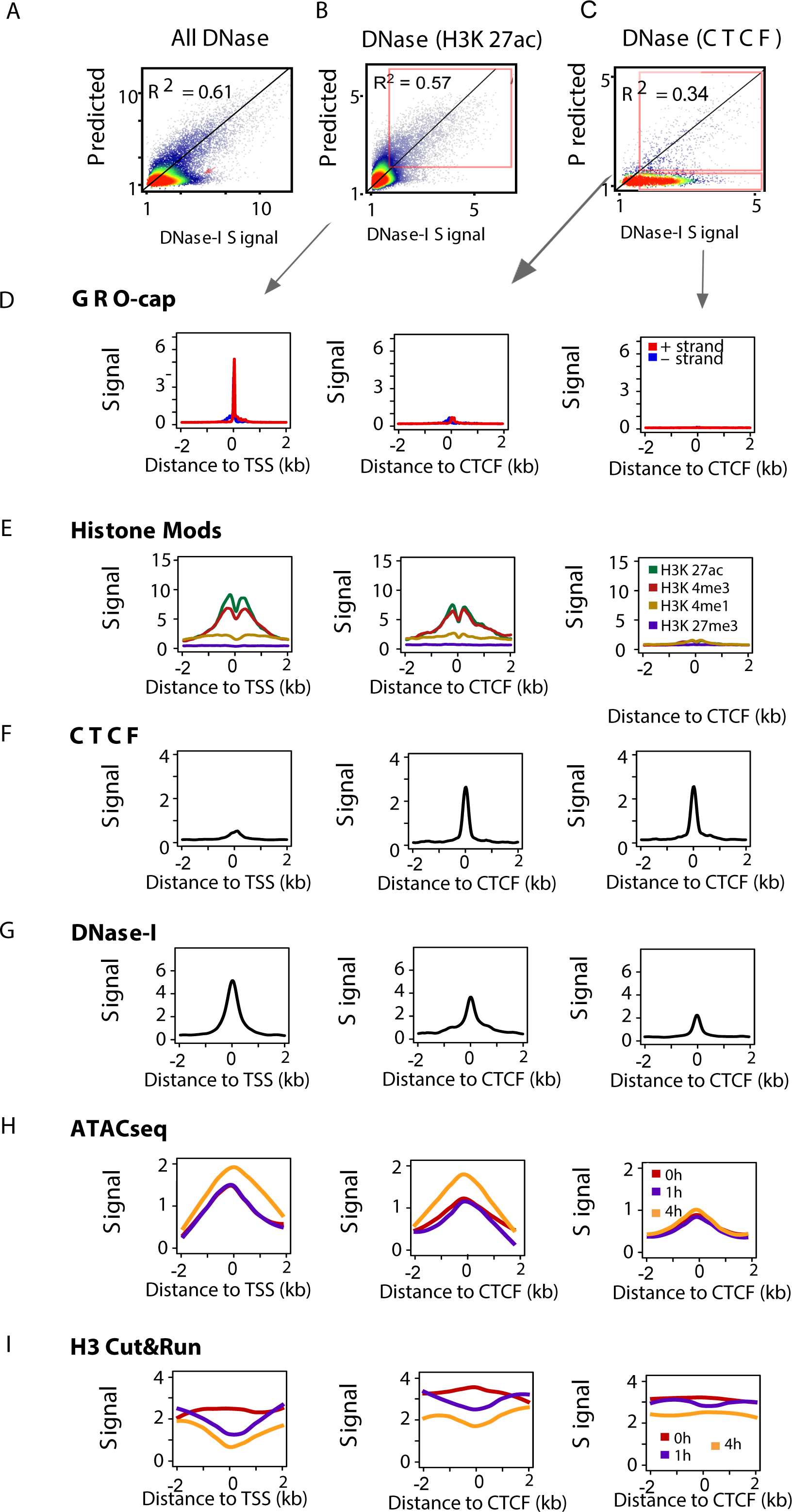
Chromatin accessibility is not sufficient for transcription initiation. (A-C) Scatterplots show experimental DNase-I hypersensitivity (x-axis) as a function of predicted DNase-I hypersensitivity (y-axis) in 100bp windows intersected with DNase-I hypersensitive sites (A), H3K27ac (B), or CTCF peaks (C) on a holdout chromosome (chr22). (D-G) Meta plots show GROcap, histone modifications, CTCF binding, and DNase-I hypersensitivity signal near H3K27ac peaks in which DNase-I hypersensitivity signal was accurately predicted by transcription (left), near CTCF peaks in which DNase-I hypersensitivity signal was accurately predicted by transcription (middle), and near CTCF peaks in which DNase-I hypersensitivity signal was not accurately predicted by transcription (right). (H-I) Meta plots show ATACseq (H) and CUT&RUN H3 signal (I) at regions in D-G.

**Figure 6.**
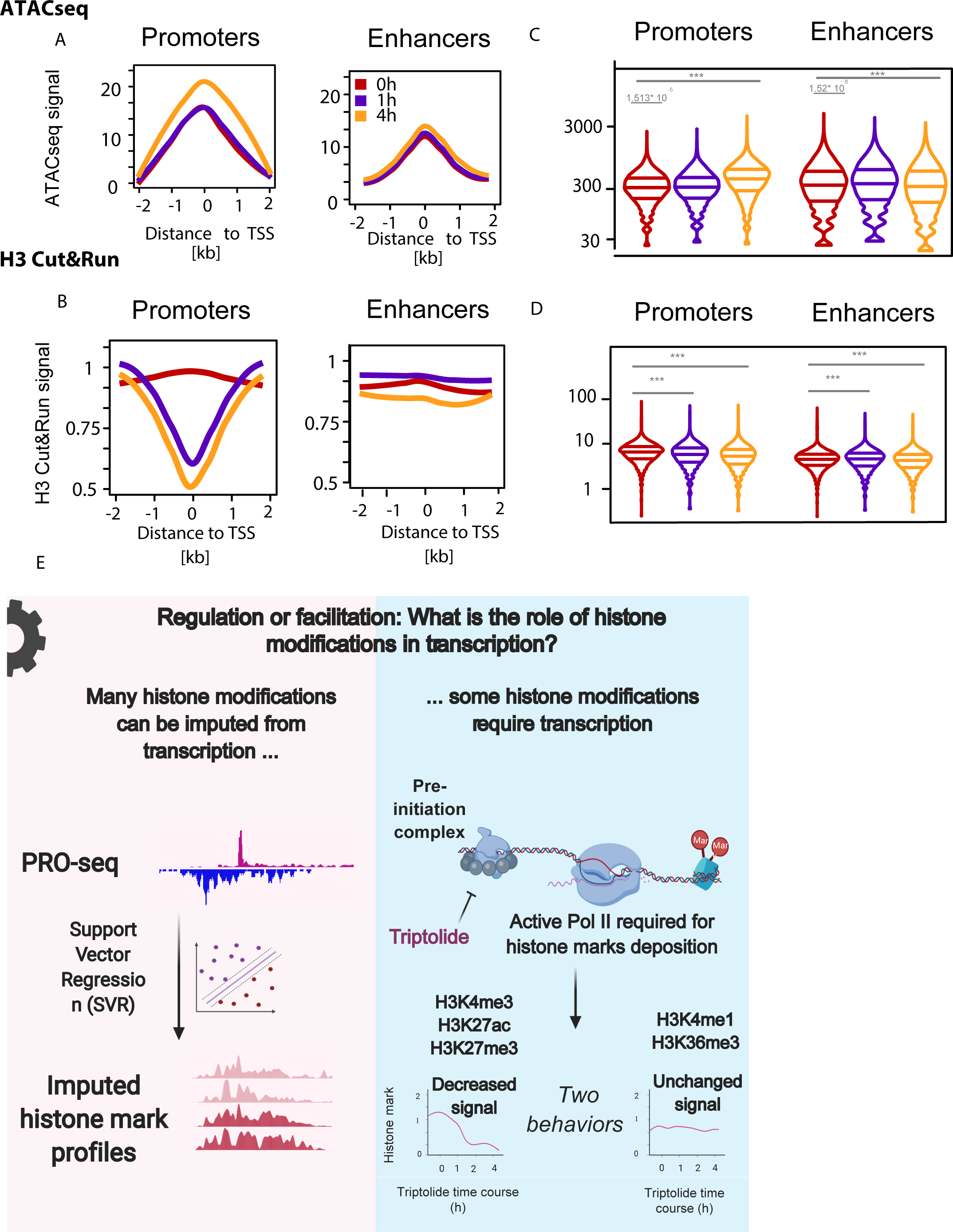
Transcription is required for chromatin landscaping. (A-B) Meta plots display ATAC-seq (A) and H3 CUT&TAG (B) signal measured at gene promoters and enhancers. (C-D) Violin plots quantify the change in ATAC-seq (C) and H3 CUT&TAG (D) signals at gene promoters and enhancers. Significance was calculated by Wilcoxon test, where (***) denotes p-value < 2.2e-16. (E) Summary figure

In parallel with PRO-seq we also performed MNase-ChIP-seq (ChIP-seq) for four active histone marks: H3K4me1, H3K4me3, H3K27ac, H3K36me3, and one repressive mark: H3K27me3. To normalize libraries for systematic variation in MNase cutting and immunoprecipitation efficiency, we added *D.iulia* butterfly cells to each immunoprecipitation as a spike-in control and normalized data using a modified version of the Spike Adjustment Procedure (SAP) normalization strategy^78^ (see **Methods**). Collectively, our data collection and analysis strategy resulted in ChIP-seq experiments that were highly correlated with each other and with public ENCODE datasets (**Supplementary Fig. 20D-E, G**).

Analysis of MNase-ChIP-seq data revealed that histone modifications have a broad range of dependence on Pol II. Trp had no effect on either H3K36me3 or H3K4me1 (**Fig. 4D-E; Supplementary Fig. 21A-B**). Although H3K36me3 is deposited co-transcriptionally^79–81^, it has a long half-life on chromatin^82^, which the 4 hour time point used in our study is not likely to have captured. Surprisingly, two punctate marks, H3K27ac and H3K4me3, were rapidly lost by 1h of Trp and remained low after 4h of Trp (**Fig. 4F-I, top**). Western blotting for chromatin bound modified histones confirmed the global loss in H3K27ac and H3K4me3 signal observed by ChIP-seq, as well as the muted effects on H3K36me3 and H3K4me1 (**Fig. 4J, bottom; Supplementary Fig. 22**). Taken together, these results indicate a surprising and rapid dependence of punctate histone marks on on-going transcriptional activity.

The loss of active histone modifications from chromatin may be caused by either rapid enzymatic deacetylation or demethylation of histone tails or increased nucleosome turnover at TSSs. To differentiate between these hypotheses, we focused on H3K27ac. We performed additional Western blots in cells treated with a combination of triptolide and the pan-deacetylase inhibitor, Trichostatin A (TSA). In the presence of Trichostatin A, H3K27ac was retained on chromatin (**Supplementary Fig. 23**). Moreover, triptolide and TSA did not have a major impact on cell viability at the time points used in our present study, indicating that effects on chromatin were unlikely to be explained by an impact on cell viability (**Supplementary Fig. 24**). Collectively our results suggest that rapid deacylation of H3K27 is responsible for the loss observed after blocking transcription.

Next we examined a small number of sites (<5%) at which H3K4me3 or H3K27ac did not appear to change dramatically following Trp treatment. Retention of signals on this small set of sites could be explained in part by Pol II independent histone mark deposition acting at loci with low levels of Pol II prior to Trp treatment. Indeed, all of the sites with little or no loss in signal had low levels of Pol II in untreated cells (**Fig. 4K; Supplementary Fig. 25A-B**). We explored one possible mechanism for depositing histone modifications in the absence of Pol II: the CxxC zinc finger protein 1 (CFP1) binds unmethylated CpG dinucleotides and recruits SET1, the main H3K4me3 methyltransferase^83^. Sites which retained H3K4me3 had significantly higher density of CpG dinucleotides (**Fig. 4L**). Moreover, this enrichment was not found for sites that retained H3K27ac (**Fig. 4L**), illustrating that retention near CpG dinucleotides was specific to H3K4me3. Thus, at least for H3K4me3, a reasonable model is that the bulk of histone modification is deposited in a manner that is dependent on Pol II. Small amounts of H3K4me3 can be deposited in a manner that depends on other factors, but Pol II is critical to achieve high levels at most loci. These findings highlight a critical role for Pol II in maintaining H3K4me3 and H3K27ac on chromatin.

### Transcription required for H3K27me3 near PRC2 binding sites, but not for H3K27me3 spread

We also analyzed the PRC2 dependent repressive mark, H3K27me3, after Trp treatment. H3K27me3 is deposited in large domains by the Polycomb repressive complexes 1 and 2 (PRC1/2)^84, 85^. H3K27me3 intensity was decreased following Trp treatment near focal binding sites of Enhancer of zeste homolog 2 (EZH2), a component of the PRC2 complex (**Supplementary Fig. 21C**), consistent with a requirement for RNA in recruiting PRC2 and depositing H3K27me3^86^. A small number of EZH2 binding sites contained divergent transcription, detectable even at low PRO-seq sequencing depth, that was lost coincidently with H3K27me3 (**Supplementary Fig. 21D**). However, 1-4h of Trp did not cause large changes in H3K27me3 accumulation over broad regions away from PRC2 (**Supplementary Fig. 21C**), and we found no evidence for global changes in H3K27me3 by Western blotting (**Fig. 4J**). Likewise, although the deletion of the *Ephx1* TSSs led to an accumulation of H3K27me3 over gene bodies over long time-scales^87^, acute and genome-wide inhibition of transcription by Trp did not increase H3K27me3 signal on gene bodies in general (**Supplementary Fig. 21E**). Thus, while transcription may prevent H3K27me3 spread over extended timescales, inhibiting transcription acutely (1-4h) did not change the level of H3K27me3 that is broadly distributed away from PRC2 regions. These findings suggest that H3K27me3 is consistently renewed near PRC2 binding sites in a transcription-dependent manner, but that H3K27me3 levels across heterochromatin are reasonably stable.

### Chromatin accessibility is not sufficient for transcription initiation

In classical models, gene regulation in eukaryotes primarily involves removing nucleosomes from the promoter of active genes, at which point Pol II initiates in an indiscriminate manner^88^. More recent studies support such accessibility models by observations that Pol II initiates at nearly all DNase-I hypersensitive chromatin^89, 90^. However, these recent studies are controversial, and at odds with other literature showing only a subset of DNase-I hypersensitivity sites have evidence of active transcription^11, 38, 91, 92^. Central to this controversy are a number of various arbitrary choices used in each study to identify DNase-I hypersensitive sites that putatively are (or are not) transcribed. To more rigorously detect transcription at DNase-I accessible regions, we trained an SVR to impute smoothed DNase-I-seq data using PRO-seq in the same manner as we used for histone modifications. The best model predicted a holdout chromosome (chr22) with an accuracy of 0.61 or 0.77 (R^2^) at resolutions of 100 and 1,000 bp (**Fig. 5A-C**), consistent with a strong correlation between chromatin accessibility and transcription initiation^13, 89^. Nevertheless, a substantial number of DNase-I hypersensitive sites had predicted values near zero, indicating a subset of sites that were refractory to prediction based on PRO-seq transcription data (**Fig. 5A, red arrow**). Intersecting experimental and imputed DNase-I-seq intensities (100 bp windows) with ChIP-seq data revealed that poorly performing windows were enriched for binding of CTCF (**Fig. 5C**), or to a lesser extent for transcriptional repressors and co-repressors such as REST, RFX5, or HDAC2 (**Supplementary Fig. 26**). In contrast H3K27ac peaks were depleted for 100 bp windows with poor matches between experimental and imputed DNase-I-seq data (**Fig. 5B**).

To confirm the absence of transcription and further investigate the chromatin environment at each of these sites, we divided 100 bp windows into those in which DNase-I-seq was predicted well by PRO-seq, and those for which it was predicted poorly (**Fig. 5B-C, red boxes**). Windows in which DNase-I-seq was predicted well by dHIT for both CTCF and H3K27ac had a high signal for transcription initiation in GRO-cap data, which measures transcription initiation, and active histone modifications (H3K27ac, H3K4me3, and H3K4me1) (**Fig. 5D-E**). Windows in which DNase-I-seq was predicted poorly had a high CTCF signal, but virtually no evidence of transcription initiation based on GRO-cap, and weak signal for active histone modifications (**Fig. 5F**). Yet, despite substantial differences in histone marks, the quantity of DNase-I-seq signal was similar in these regions (**Fig. 5D-G, l**). Thus, a substantial portion of DNase-I accessible regions show no evidence of transcription initiation. Our analysis supports a model in which both chromatin accessibility and other aspects of the local chromatin environment, including transcription factors, pre-initiation complex machinery, chromatin remodelers, and other transcription related proteins, are all necessary to facilitate transcription initiation by Pol II.

### Chromatin accessibility at transcription start sites does not depend on transcription

Paused Pol II is necessary for proper nucleosome positioning^93^, although it may not be required to establish sufficient levels of chromatin accessibility for other biological functions to take place, such as transcription factor recognition. To test the hypothesis that chromatin accessibility requires paused Pol II, we treated K562 cells with Trp to prevent transcription initiation. Unexpectedly, a time-course of Trp treatment resulted in a small but significant increase in Tn5 accessibility, measured using ATAC-seq (**Fig. 5H, 6A**). To more precisely examine the position of nucleosomes, we performed CUT&RUN for histone H3^40^. We observed a loss in H3 signal inside of the nucleosome depleted region and adjacent +1/ -1 nucleosomes (**Fig. 6B**). Changes in ATAC-seq and CUT&RUN were specific to DNase-I hypersensitive sites that had robust evidence of transcription initiation (**Fig. 5H-I**). Notably, changes in both H3 and ATAC-seq signals were observed exclusively in transcription initiation regions, and did not appear in CTCF-bound and untranscribed control regions (**Fig. 5H-I, 6K-L**). Thus, we conclude that events prior to transcription initiation are primarily responsible for nuclease accessibility.

Two mechanistic models could explain the loss of nucleosomes near transcription initiation regions upon Trp inhibition of transcription. First, Trp inhibition of TFIIH activity could increase the time that Pol II spends in the PIC or initiation mode, thereby keeping nucleosomes at bay. Second, pioneer factors, which bind and cooperate with SWI/SNF chromatin remodelers to open chromatin^94, 95^, could be locked in a histone evicting mode by the failure of Pol II to properly pass through initiation. To distinguish between these models, we performed CUT&RUN for TATA-binding protein (TBP), which showed increased TBP occupancy following 30 min of Trp (**Supplementary Fig. 26G-H**). This result supports the idea that the residence time of the pre-initiation complex plays some role in establishing chromatin accessibility following Trp treatment. Notably, our results mirror observations of chromatin accessibility during mitosis^96^, a cellular context during which Pol II is depleted from chromatin, but accessibility to Tn5 and the signal for TBP are increased. Thus, although paused Pol II may help to establish the position of +1 and -1 nucleosomes near open chromatin regions^93^, chromatin accessibility can be established by multiple factors and does not necessarily require Pol II initiation or pausing.

## Discussion

We have conducted an extensive analysis of the relationship between transcription, histone modifications and open chromatin. Our work leverages novel machine learning methods trained to infer the genomic distribution of 9 widely studied histone modifications using nascent transcription as input. We complemented new analytical tools with experiments perturbing both transcription and histone modifications. Our work provides new insight into the role of histone modifications in regulating transcription by Pol II. Collectively, our results support models in which active histone modifications are “cogs” with a supportive role, rather than a direct regulatory role, in transcription. Finally, we provide a new strategy for genome annotation using a single functional assay that is tractable for a single lab to perform.

### Active histone modifications as essential cogs, rather than causes, of transcription

Our incomplete knowledge about the role that histone modifications play in transcription results in part from a lack of information about the precise strength of correspondence between histone modifications and transcription. We demonstrate that the correlation between histone modifications and transcription is nearly as strong as the correlation between biological replicates of experimental histone modification ChIP-seq data. Moreover, we likely underestimate the actual correlation between transcription and histone modifications, due to technical factors including imperfections in the model fit, low resolution experimental procedures, and biological differences between cells cultured in different labs. The strong correspondence between histone modification and transcription that we have observed here addresses several open questions about the biological role of histone modifications. For instance, do histone modifications serve in part to “bookmark” critical functional elements for later activation by developmental or environmental processes? The strong correspondence that we observe is not compatible with models where histone modifications routinely “bookmark” future transcription events. Rather, our work indicates that histone modifications reflect the transcription patterns active in the current cell state. Do different “chromatin states” comprised of distinct histone modifications (e.g., H3K122ac and H3K27ac) interchangeably produce similar transcriptional outcomes in distinct parts of the genome? Our results indicate that histone modifications serve as critical pieces of a uniform molecular machinery (i.e., cogs or gears), which are highly interconnected with Pol II and play a supportive role in transcription.

A cog model implies that the genomic distribution of histone modifications and Pol II are highly interdependent on one another. In support of this, blocking Pol II transcription initiation for short durations (1-4 h) had rapid and large-scale effects on the genomic distribution of three histone modifications: H3K4me3, H3K27ac, and H3K27me3. Surprisingly, punctate focal marks, H3K4me3 and H3K27ac, were almost entirely dependent on active transcription to remain on chromatin. For lysine acetylation, this result mirrors elegant complementary experiments focused on histone acetylation in yeast^97^. Loss in punctate histone modifications coincides with a rapid loss in histone H3 associated with regulatory regions, indicating that nucleosome turnover may play some role in punctate mark depletion from promoters after Trp. However, several lines of evidence indicate that, at least for H3K27ac, mark removal by deacetylases plays a role as well. First, the loss in H3 does not appear large enough near the +1 or -1 nucleosome to explain the substantial depletion in punctate histone mark (although we note that CUT&RUN and ChIP-seq are substantially different assays, precluding a quantitative comparison of changes). Second, we also found that depletion of H3K27ac after blocking transcription was prevented by HDAC inhibitors, indicating that active transcription affects the intricate balance between the addition and removal of histone acetylation. Although two additional marks, H3K4me1 and H3K36me3, do not undergo major changes during the 4h after blocking transcription initiation, this is likely explained by the known longer half-life of these particular methylation marks on chromatin^82^.

Despite an extremely large loss in H3K4me3 at over 90% of loci as a result of transcription inhibition, a small number of focal modifications remain on chromatin. Loci retaining H3K4me3 tend to be highly CpG-rich, potentially consistent with CFP1 binding unmethylated CpG dinucleotides and recruiting SET1 to deposit H3K4me3^83, 98^. These unaffected promoters have both low transcription and low H3K4me3 prior to the time-course. Taken together, these results suggest two modes of H3K4me3 deposition: a transcription-dependent mode, which appears to be required for the vast majority of H3K4me3, and a minor component which is dependent on recruitment of transcription activators in response to DNA sequence features. Our model of CFP1 is compatible with the broad notion that transcription factors drive patterns of both histone modifications and Pol II transcription. Our work is also compatible with many histone modifications being unstable due to a constitutive interplay between enzymatic erasers of the marks in dynamic competition with writers^99, 100^. Additionally, recent studies depleting histone modifications widely believed to be critical for transcription have found surprisingly limited effects on gene expression^101–103^. These studies support a model in which most active histone modifications work to fine-tune aspects of transcriptional regulation, rather than regulating gene expression on their own.

### Facultative heterochromatin responds to perturbations in transcription

A great example of the interconnected loop between transcription and histone modification is H3K27me3. H3K27me3 is a maker of facultative heterochromatin and has long been hypothesized to serve a role in transcriptional repression. Blocking transcription initiation led to a decrease in H3K27me3 near PRC2 binding sites, consistent with recent work^86^. Our study supports the model in which transcription initiation produces short RNAs that recruit PRC2 to deposit H3K27me3, as observed at the Xist locus^104^. Nevertheless, less differentiated cell types appear to just have punctate patterns of H3K27me3 near PRC2 binding sites, without the broad H3K27me3 spread across a domain. As an example, when comparing more differentiated cell types such as primary or adult tissue with multipotent or pluripotent cell stages, H3K27me3 shifts its pattern from large domains to narrow, punctate regions. These less differentiated cell types may have a tightly regulated balance between mark addition and removal that prevents H3K27me3 from spreading.

### Chromatin accessibility: Necessary, but not sufficient, for transcription

Our results also have implications for the debate about whether transcription initiates pervasively at open chromatin regions. Classical models of gene regulation developed during the 1970s held that histones were general suppressor proteins that passively silenced gene expression^88^. Genome-wide analyses uncovered distinct classes of DNase-I accessible regions that either do or do not have evidence of transcription initiation^11, 91, 92^. Nevertheless, these recent results remain actively debated by papers arguing that all nucleosome depleted regions initiate transcription with some frequency, regardless of whether they show histone modifications or regulatory activity^89^. We show that there were a substantial number of DNase-I accessible open chromatin regions that were not identified by dHIT imputation. Unlike previous work, which relied on arbitrary heuristics to select sites with or without evidence for transcription, dHIT allowed us to directly identify candidate DNase-I accessible regions with atypically large imbalance between experimental and predicted transcription. These DNase-I accessible regions had no evidence of either transcription or histone modifications but were enriched for other chromatin binding proteins like CTCF. Likewise, blocking transcription had a small impact on chromatin accessibility uniquely at transcribed DNase-I hypersensitive sites that were accurately predicted by dHIT. Our findings indicate chromatin accessibility is not sufficient for transcription initiation, ruling out the hypothesis that Pol II is so promiscuous that it simply uses any accessible DNA to initiate^89, 90^. Rather, Pol II requires specific features of core promoters^105–107^ or local protein-DNA environment, such as transcriptional activators, to initiate transcription.

### dHIT: A powerful tool for genome annotation

Currently genome annotation requires conducting assays for multiple independent histone modifications to identify functional elements. Since functional elements are known to be highly tissue specific and their activity is dependent on environmental conditions, assays must be performed in numerous tissues and conditions to exhaustively identify functional elements. As a result, genome annotation efforts still benefit greatly from the coordinated efforts of large consortia. However, consortium efforts are not tractable to apply in all species and tissues, especially as major efforts to sequence eukaryotic organisms begin to produce large numbers of high-quality reference genomes^37^. This creates a great need for individual communities to annotate functional elements using the most efficient molecular and computational tools. We show here that nascent transcription measured using PRO-seq provides at least as much information about chromatin state as the combination of multiple ChIP-seq datasets.

Why is dHIT so effective at predicting most active histone modifications? Our success in training dHIT was possible because PRO-seq data is an uncharacteristically rich source of information about genome function. PRO-seq provides the density of RNA polymerase across the genome at single nucleotide resolution and in a manner that is not confounded or restricted by RNA processing or RNA stability. PRO-seq directly measures polymerase densities on all transcription units (which is highly correlated with mRNA^108^) and it also provides signatures of active regulatory activity at both promoters and enhancers. It picks out promoter and enhancer pause peak positions where Pol II accumulates to high density as part of a checkpoint or regulatory mechanism. And, for many histone marks that are intimately connected to transcription and/or transcription states, like H3K36me3, H3K4me3, H3K27ac, the predictions made by dHIT are especially strong.

dHIT complements existing algorithms that predict transcription units and transcriptional regulatory elements. Tools such as NRSA^109^ and Tfit^110, 111^ leverage similar information, such as the shape or density distribution of nascent transcription, to annotate functional elements in eukaryotic genomes. dHIT builds on these approaches by training accurate machine learning models to transfer transcription “shape” features into chromatin states. Chromatin state annotations made using PRO-seq data could provide an efficient path to genome annotation, especially when complemented by experimental data for which dHIT provides incomplete or inaccurate information (e.g., H3K9me3, H3K27me3, and open chromatin). Our view is that, depending on the problem at hand, dHIT/ PRO-seq would be complemented best by the addition of experimental H3K9me3 (which we are not able to predict at all), followed by ATAC-seq (which adds the position of candidate insulators) and H3K27me3.

Even using just PRO-seq data alone, the development of dHIT maximizes information obtained about genome function from a single experiment. In addition to chromatin state, nascent transcription is also known to provide direct information about gene expression^108^, transcription factor binding^110^, the location of transcription start sites, and the grammar of transcription initiation domains^11, 15, 56, 91^. Moreover, the introduction of new biochemical tools that allow the application of PRO-seq techniques with greater ease in solid tissues and other samples that have proven challenging to measure using conventional genomic techniques has the potential to further democratize these technologies^54, 112^. Thus, using dHIT to decompose PRO-seq data into separate information about active chromatin modifications is a supremely efficient strategy to gain information about functional elements using a single molecular assay for each tissue and condition.

## Methods

### Experimental methods

#### Cell culture

K562 cells (ATCC, CCL-243) were cultured at 37°C, 5% CO_2_ at a density between 0.3-1 x 10^6^cells/mL in RPMI medium (VWR 45000-396) topped up with 10% Fetal Bovine Serum (Genesee Scientific, cat: #25-514). Cells were split at a consistent interval of 3 days, when the cells reached 10^6^ cells/mL.

#### Cell culture for Triptolide and Trichostatin A time course

24h prior to drug treatments, K562 cells were resuspended in fresh (RPMI) medium at a density of 0.6 x 10^6^ cells/mL. On the day of the experiment, cells were recounted, aliquoted in equal cell numbers to T-25 or T-100 ThermoFisher Tissue Culture Flasks (each flask corresponding to one time point) and treated with Triptolide (Sigma-Aldrich, T3652-1MG) or Trichostatin A (Sigma-Aldrich, T8552-1MG). Final concentrations used in our experiments were: 500 nM Triptolide, and 250nM Trichostatin A. All drug treatments were performed for 0 min, 1h, and respectively 4h.

#### Cells cross-linking for ChIP

After Triptolide treatment, K562 cells were cross-linked in 1% CH_2_O freshly prepared in 1x PBS on the day of the experiment to reach the final concentration of 0.1% CH_2_O in the media. Following a 5 min incubation at room temperature on a rocking platform, the cross-linker was quenched with 1M Glycine to reach a final concentration of 0.135 M Glycine. Lastly, cells were washed twice in 1x PBS, then harvested and snap frozen on dry ice.

#### MNase ChIP-seq - chromatin extraction

We prepared MNase ChIP-seq data for six histone marks in K562 cells, including H3K4me1 (ab8895, lot: GR3206285-1), H3K4me2 (ab7766, lot: GR102810-4), H3K4me3 (ab8580, lot: GR3197347-1), H3K27ac (ab4729, lot: GR3231937-1), H3K36me3 (ab9050, lot: GR3257952-2), and H3K27me3 (ab6002, lot: GR3228496-2). All buffers and solutions used were provided by Cell Signaling Technology (91820S Simple ChIP kit). Cross-linked K562 cells were thawed on ice and resuspended in 1 mL cold Buffer A, mixed well, and centrifuged at 2000x g for 5 min at 4°C. The pellet was then mixed in 0.5 mL cold Buffer B, centrifuged at 2000x g for 5 min at 4°C andresspended again in Buffer B. While still in Buffer B, chromatin was digested with 0.5 uL MNase for 13 min at 37°C. Tubes were inverted every 2 min during the incubation time. Finally, the reaction was stopped by the addition of 40 uL 0.5 M EDTA, and the tubes were moved to 4°C. The cell suspension got topped up with 1.5 mL cold ChIP Buffer, transferred to a 7 mL glass dounce homogenizer, and dounced ∼30 times with a tight pestle to release the chromatin. The chromatin was further diluted with 1 mL cold ChIP Buffer and aliquoted to 1.5 mL Eppendorf tubes to be centrifuged at 12000x g for 10 min at 4°C. The supernatant was collected and total chromatin quantified before each immunoprecipitation.

#### MNase ChIP-seq - Immunoprecipitation

Total digested chromatin was diluted to a total volume of 1 mL in cold ChIP Buffer. ChIP samples were incubated with 3ug anti-histone antibody at 4°C overnight rotating, then incubated for an extra 2h at 4°C with 20 ug magnetic beads (50% protein A, 50% protein G). After incubation, samples were placed on a magnetic rack and washed three times with 1 mL Low Salt Wash Buffer for 5 min at 4°C, and three times with High Salt Wash Buffer for 5 min at 4°C. Lastly, the beads were resuspended in 150 uL Elution Buffer and incubated on a shaking Thermomixer for 1.5 h at 65°C. The eluted fractions were saved, treated with 2 uL 5M NaCl and 10 uL Proteinase K, and incubated overnight at 65°C to reverse the cross-linker. Samples were cleaned up, the DNA quantified with Qubit, and library prep was performed using the NEBNext Ultra II DNA Library Prep Kit for Illumina (E7645S). The barcodes used were purchased from NEB:NEBNext Multiplex Oligos for Illumina (E6440S). Before Bioanalyzer and Illumina sequencing, all libraries were size-selected by being run on a 6% Native PAGE. The fragments corresponding to 200-700 bp were cut out of the gel and the DNA extracted from the polyacrylamide using 3 volumes of a DNA extraction buffer (10mM Trip pH=8, 300mM NaAc, 20mM MgCl2, 1mM EDTA, 0.1% SDS) per gram gel slice. The tubes were closed, covered with parafilm, and incubated overnight at 50°C shaking, on a Thermomixer. The following day, Spin-X columns (CLS8160, Millipore Sigma) were used to remove gel bits from the eluate which got Phenol/Chloroform precipitated. The precipitated DNA was resuspended in a 15 uL nuclease-free H_2_O and the library quantified using Qubit.

#### Measuring chromatin-associated proteins by Western blotting

For the Triptolide experiment, we used matched cells with the ones in the ChIP-Seq experiments. The Trichostatin A Western blots were performed on cells of a different passage number. For each reaction, 500,000 K562 Triptolide treated-cells were thawed on ice, spun down at room temperature in a swing bucket centrifuge for 5 min, then washed twice in 5mL Permeabilization buffer (10mM Tris-HCl pH 7.5, 10mM KCl, 250mM Sucrose, 5mM MgCl_2_, 1mM EGTA, 0.05% Tween-20, 0.5 mM DTT, 40 units/10mM AM2694 SUPERaseIn Thermo Scientific, 0.2% NP-40, A32963 EDTA-free PIERCE Protease Inhibitors). Cells were incubated on ice with Permeabilization Buffer 5min each eashing time. Isolated nuclei were verified by Trypan Blue staining. Chromatin-bound proteins were isolated by centrifugation at 12,500xg for 30min, at 4C. After centrifugation each cell pellet was dissolved in 2xSDS loading dye and syndicated on high setting for 5 min (30s ON: 30s OFF). Samples were boiled at 95C for 5min and loaded on a 15% SDS-PAGE gel. The same antibodies used for ChIPseq were used for western blotting. Cell Signaling Technology 9715 anti-H3 antibody and abcam 8WG16 anti-Pol II were also used.

#### Measuring cytotoxicity of Triptolide and Trichostatin A

Cells were grown in a 96-well plate following the “*Cell culture for Triptolide and Trichostatin A time course”* protocol. On the day of the experiment, cells were treated with either Triptolide alone or a Triptolide + Trichostatin A dual treatment, then incubated with almarBlue (BIORAD, BUF012A) following the BIORAD protocol. Absorbance of cells incubated with almarBlue was measured at 590 nm. The experiment was performed in two biological replicates and compared with a DMSO kill curve as positive control, and cells untreated with any drugs as positive controls.

#### CUT&RUN

We measured histone H3 (Cell Signaling Technology 9715) and TBP (ab818) levels on chromatin following a time-course of Trp treatment using the High Ca2+ / Low Salt CUT&RUN protocol (https://dx.doi.org/10.17504/protocols.io.zcpf2vn). Each experiment was performed with 250,000 K562 cells.

#### ATACseq

K562 cells were treated with Triptolide and a total of 500,000 cells per condition were used for ATACseq. After Triptolide treatment, cells were washed in 1xPBS, then lysed in 1mL cold Lysis Buffer (10mM Tris-HCl pH 7.4, 10mM NaCl, 5 mM MgCl_2_, 0.2% NP-40) while incubating for 3min on ice. The lysis buffer was removed by 10min centrifugation at 600xg, 4C, and cell pellets were resuspended in 48.5uL Transposition Buffer (10mM Tris-HCl pH 7.4, 10% DMF, 5 mM MgCl_2_). 1.5uL in-house purified Tn5 (stock concentration 3.5 ug/uL) was used per reaction. The transposition took place for 30min at 37C while shaking. Phenol/Chloroform was used to extract transposed DNA which was further PCR amplified to add NExtera sequencing adapters.

#### PRO-seq library prep

New PRO-seq or ChRO-seq libraries were prepared from cultured K562 cells, and from equine liver tissue samples. We prepared PRO-seq libraries in K562 cells, matched to the MNase ChIP-seq. *D.melanogaster*, S2 cells, were used as heterogeneous spikeins and added to each sample before the run-on in a ratio of 1:10,000 = S2:K562 chromatin.

### Data processing for newly collected MNase ChIP-seq, CUT&RUN, ATAC-seq, and PRO-seq

We used hg19 as the primary genome assembly in our data analyses to facilitate comparisons with ENCODE data (which primarily used hte hg19 assembly at the time these analyses were conducted). Data from each experiment was aligned to genome assemblies as following:

-MNase ChIP-seq reads were aligned to hg19 merged to *D.iulia* assembly^113, 114^. All positions with sequence similarity between the two genomes were masked using bedtools maskfasta;

-ChRO-seq data was aligned to hg19 merged to the *D.melanogaster* dm3 genome assembly. All positions with sequence similarity between the two genomes were masked using bedtools maskfasta;

-CUT&RUN was aligned to hg19 merged to *S.cerevisiae* SacCer1;

- ATAC-seq was only aligned to hg19.

Masking hg19 was performed with BedTools maskfasta ^115^. All sequencing data was aligned using bowtie2 version 2.3.5.1 ^116^ with parameters: --no-discordant --no-dovetail --no-unal --no-mixed. Reads mapping multiple times were removed with samtools view ^117^, parameter: -F 256. The remaining reads were converted to paired-end BigWig files using BedTools and visualized in the WashU genome browser version 46.2 ^118, 119^.

#### ChIP-seq normalization strategy (for MNase-seq triptolide time course)

In our experiments, both the human and spike-in samples were mixed and treated with MNase together, before the antibody incubation. To correct IP signals for biases in MNase cutting efficiency, handling, and other errors, we used the spike adjusted procedure (SAP) method ^78^. We assume that ChIP-seq data reflects a linear combination of three factors: signal from the mark of interest, background which may be partially correlated with the mark, and random noise. SAP assumes that the background signals should be the same in treated and untreated samples and enforces this assumption by subtracting the expected background read count observed in the input. Because the data is noisy and we cannot assume input samples are sequenced deeply enough to estimate the background directly, SAP subtracts the expected background estimated using a linear regression fit in background regions. To define background regions, we selected a set of coordinates in the human genome with the following properties:

- They are untranscribed. The number of reads aligning within these sites should not change during the Trp time course. We first masked human coordinates for all annotated gencode transcripts, then removed regions near PRO-seq reads aligned to hg19;

- They are found outside MACS2-called ChIP-seq peaks ^120^. The number of reads aligning within these sites should not be affected by differences in IP efficiencies;

- They are located in accessible chromatin and have a broad range of MNase sensitivities that cover the range observed in loci of interest.

To satisfy all these requirements we found a set of ENCODE CTCF peaks in K562 that were located at least 40kb away from any annotated transcription initiation region (TIRs). All TIRs were identified using dREG ^91, 121^. In the case of punctate histone modifications (H3K27ac, H3K4me1, H3K4me3), we counted the input and IP reads in each 500bp bin over a 5.5kb region centered on the CTCF peak. To capture the variation in MNase accessibility for marks deposited within broader domains (H3K36me3 and H3K27me3) we enlarged the bin size to 1kb spanning 30kb adjacent to the CTCF peaks. To account for differences in IP efficiency and MNase accessibility due to biases in chromatin accessibility, we then ranked CTCF peaks by their DNase-I hypersensitivity (DHS) and summed up the counts in each bin into 10 deciles.

##### Box 1: Notations

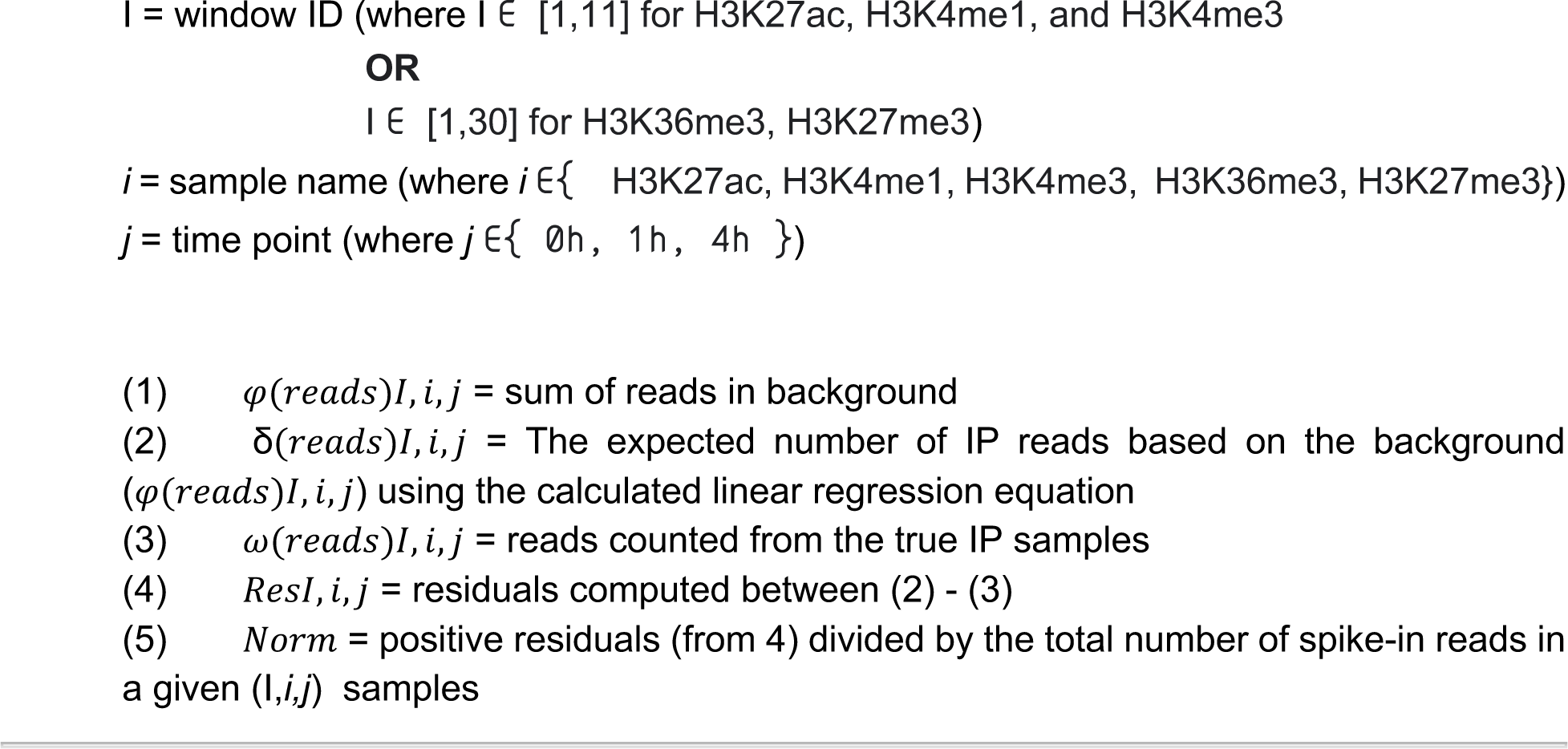

We denote the raw, observed signal in each window as:

Let *φ*(*reads*)*I*, *i*, *j* represent the raw signal per background window (I) in each IP sample or its corresponding input control (*i*) and time point (*j*).

SAP assumes that reads in each IP sample should be proportional to the input sample in background regions. To estimate the expected signal between input and IP, we fit a linear regression between the 10 deciles in the background regions of input and IP, resulting in estimates of two regression coefficients, α and β. Next, we fixed α and β and used those parameters to estimate the expected IP signal within given sets of MACS2-called peaks (δ(*reads*)*I*, *i*, *j*) as a function of the input:

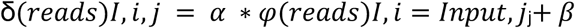

To estimate the contribution of histone mark to the signal in each IP sample in each window of interest (e.g., a MACS2 peak), we then computed the residual (*ResI*, *i*, *j*) between the number of reads expected based on the linear regression equation ( δ(*I*)*i*, *j*) and the number of IP reads observed experimentally within the same regions (ω(*reads*)*I*, *i*, *j*).

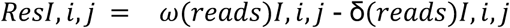

As residuals can be negative, and it does not make sense to have a negative signal for a mark, we set all negative residuals to zero, as in SAP. We note that the vast majority of loci we expected to have signals (e.g., because they were in ENCODE peaks) had residuals that were larger than 0.

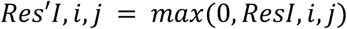

Last, to account for global changes, we divide each normalized region by the total number of spike-in reads (*Si*, *j*) in a given sample (*i*) of time point (*j*).

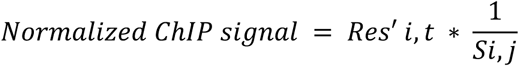

#### CUT&RUN normalization strategy

A total of 2.5pg/mL final concentration of S.cerevisiae MNase-fragmented DNA was added as spike-in control to each CUT&RUN experiment. After aligning all reads to a merged human and yeast genome, we determined the total number of yeast reads in each sample. For normalization purposes, we divided the human tags by the total number of yeast tags in each particular sample.

#### ATAC-seq normalization strategy

To account for changes during handling and sequencing of ATAC-seq libraries, we consider a constant background level between conditions. The background was estimated as the total number of tags mapping to gene-desserts, PRO-seq untranscribed, and Tn5-inaccessible coordinates in the human genome. To normalize, we divided the tags in a given sample by its respective background tags.

#### PRO-seq normalization strategy

Chromatin from *D.melanogaster* S2 was spike-in as internal control in a 1:10,000 [ng : ng] human:fly ratio. As normalization, we divided the human tags in each sample by the total number of tags aligning to the fly genome from that particular sample.

Maximum transcription start sites, as defined in Tome and Tippens, 2018 ^15^, were used to draw meta profiles of ChIP-seq, PRO-seq, CUT&RUN, and ATAC-seq signals.

### Training dHIT SVRs to predict histone marks using PRO-seq, GRO-seq or ChRO-seq data

#### Overview

The primary goal of dHIT is to map the signal intensity and “shape” in a run-on and sequencing dataset (PRO-seq, GRO-seq or ChROseq; henceforth referred to simply as PROseq) to the specific quantity of a histone modification at each position in the reference genome. The dHIT algorithm passes standardized read count data to a support vector regression (SVR) classifier. During a training phase, the SVR model optimized an objective function which mapped PRO-seq signal to the quantity of ChIP-seq signal at each position of the genome. Once a dHIT model is trained using existing ChIP-seq data, it can impute steady state histone modifications in any cell type, provided that the relationship between histone modification and transcription is preserved. The dHIT software package is provided at https://github.com/Danko-Lab/histone-mark-imputation.

### Training dataset

All data used for training were evaluated for quality content using PEPPRO^122^. We trained each model using five different run-on and sequencing datasets that were generated by different laboratories, thereby reducing the potential for overfitting to batch-specific features of a single dataset (see **Supplementary Table 2**) ^56^. Training data was distributed between PRO-seq and GRO-seq data. Sequencing depth of the training data ranged from 18 to 374 million uniquely mapped reads, and all five training datasets were highly correlated when comparing RPKM normalized read counts in gene bodies ^56^.

We trained SVR models for ten different histone modifications in K562 cells, primarily using data from the ENCODE project^16^, all of which passed the ENCODE 2 data quality standards^123^. Data for H3K122ac ChIP-seq in K562 cells was obtained from a recent paper^52^. Lastly, we trained models to recognize high-resolution ChIP-seq data using an MNase ChIP-seq protocol for H3K4me1, H3K4me2, H3K4me3, H3K27ac, and H3K36me3. For validation in holdout cell types, we obtained ChIP-seq data from six additional cell types from a variety of sources. All training and validation analyses used sequencing depth normalized read counts, where possible using bigWig or bedGraph files provided by the original authors as input. All ChIP-seq data used in training or for validation is listed in **Supplementary Tables 1 and 2**.

### SVR feature vector

We passed dHIT PRO-seq data from non-overlapping windows of multiple sizes that were centered on the position for which ChIP-seq signal intensity was being imputed. We have previously optimized the number of window sizes and the window sizes for optimal classification of TIRs using dREG ^56, 91^. Since the imputation of histone modifications uses signals in the PRO-seq data that are similar to dREG, we used the values that were optimal for dREG without modification. Like for dREG, we passed data from windows at multiple size scales, including 10, 25, 50, 500, and 5,000 bp windows (*n* = 10, 10, 30, 20, and 20 windows, respectively), representing read data as far as 100 KB from the genomic region in question. PRO-seq data was standardized across each length scale in a similar fashion as we use for dREG^91^, using a logistic function, *F(t)*, to transform raw read counts using two free parameters, α and β:

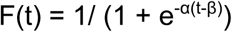

Where *t* denotes the read counts in each window. Tuning parameters α and β were defined in terms of two parameters, x and *y*. Intuitively, *y* gives the value of the logistic function at a read count of 0, and *x* represents the fraction of the maximal read count at which the logistic function approaches 1. Values of *x* and *y* are related to the parameters *α* and *β* by the following equations:

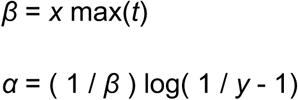

We have previously found that *x* = 0.05 and *y* = 0.01 optimized the discovery of transcription initiation regions (TIRs)^91^, and these values were used throughout this study.

#### Selecting training positions

We trained models using 3 million training examples divided evenly among five K562 training datasets (n = 600 thousand positions in each dataset). In all cases, human chromosome 22 was excluded from training to use as a holdout.

We found it convenient to use heuristics that identify regions with a high PRO-seq signal intensity when choosing training samples. We defined regions of potential PRO-seq signal, which we call “informative positions” using the same heuristics we described previously for dREG ^91^. Each window was defined as an “informative position” when the window had more than 3 reads within 100 bp on the single strand or at least one read within 1000 bp on both the positive and negative strands. These heuristics were selected as a way to optimize the tradeoff between the number of positions analyzed and the fraction of real TIRs that were scored based on the overlap with GRO-cap peaks. Within the five training datasets, informative positions accounted for 27.3% (855.9M), 6.7% (209.4M), 14.7% (460.0M), 13.8% (433.9M), and 9.4% (294.0M) of 10 bp windows, respectively.

Training examples were selected at random, according to the following criteria: In order to increase the frequency of windows with a strong signal intensity in the training dataset, we selected 5% of the training data from positions in the informative positions pool (defined above) that also intersected a transcription start site (TSS), defined using GRO-cap ^11^, and a DNase-I hypersensitive site^16^, 93% from the non-TSS informative sites, and the remaining 2% from the non-informative position pool. This was done to enrich the frequency of GRO-cap TSSs (these were 0.78% of hg19), and to increase the frequency of regions with substantial PRO-seq signal intensity, in the training dataset.

Training computations were conducted using Rgtsvm, a fast, GPU-based SVR implementation^124^. We trained 3M samples with 360 features for each sample from 5 data sets with an average training time of 27.9 hours (18.0∼37.8 hours) on an NVIDIA Tesla TITAN XP GPU. Training achieved an average Pearson correlation of 0.48 (0.109∼0.725) on holdout positions that matched the training dataset at 10bp resolution.

#### SVR imputation

We imputed histone modifications every 10 bp using the run-on and sequencing datasets outlined in **Supplementary Table 2**. We tested the accuracy of imputation on human chr22 (which was withheld during training) in four holdout cell lines HCT116, HeLa, and CD4+ T-cells^125–127^. Imputation was conducted using ChRO-seq data from 20 primary glioblastoma cases^54^. We also imputed data from two additional mammals: mouse embryonic stem cells (mESCs)^68^ and horse liver (new data). Computing imputed values on human chr22 (5.1M loci) took 3-5 hours on a Tesla TITAN XP GPU.

### Training models that impute histone marks using other histone marks

We selected 1M samples from chromosome 1 to train SVR models in which histone marks were used to predict other histone marks. In order to make a fair comparison with models trained to predict histone marks using PRO-seq data, we also trained new models from PRO-seq (using the dataset G1) using 1M samples. To select training positions when training models using histone marks, we calculated the maximum read count in every 50 bp windows on chr1 (4.99M regions), and selected 1/3 of the samples from regions that contain more read counts than median value in either the training or the experimental data (for instance, if using H3K4me1 to predict H3K4me3, we selected 33% of training positions that had higher read counts than the median H3K4me1 or H3K4me3 signal). We selected another 1/3 from regions which contained read counts that were less than 20% of the median value in either the training or the experimental data. We selected the last 1/3 of the training regions from remaining regions at random. To obtain training datasets when multiple histone marks were used to jointly predict a histone mark, we merged multiple experimental histone mark data together and sampled windows as described above. The feature vector and standardization for histone marks were identical to those used for PRO-seq data (see above). When generating the feature vectors for multiple histone marks, we concatenated the feature vectors extracted from multiple experimental histone marks together.

We compared the difference between imputation and original experimental data using the L1 norm, by median centering and scaling each dataset, as follows:

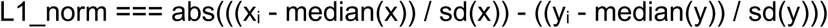

Where xi is the imputed signal, and yi is the experimental signal for a particular comparison, and i represents the set of all genomic positions on chr22. We use sd() to denote the standard deviation of the mark.

### Computing performance metrics using dHIT SVRs

Imputed profiles for 10 histone modifications in seven cell lines were compared to a variety of publicly available and newly generated ChIP-seq data available from ENCODE, Epigenome Roadmap, and a variety of other sources, as outlined in **Supplementary Table 1**. When measuring correlations, we subtracted the background (median) value from all positions, and applied a series of filters that were designed to remove artifacts of mappability or repeat content. Filters used to compute correlations include: 1) We masked all positions in which 30bp, the size of many of the older ENCODE ChIP-seq datasets, can not map uniquely to the reference genome; 2) We removed ENCODE blacklist regions annotated on hg19^128^; 3) We identified and masked “spikes” in the data, caused by putative experimental or mapping artifacts, that were not filtered by the above two criteria. Our filter identified blocks with a high signal intensity (top 2%) for which the sum of the absolute value of the two maximal derivatives was higher than the number of read counts in the region (i.e., [abs(d1) + abs(d2)] > h, where d1 and d2 are the maximal and second highest change in ChIP-seq signal intensity, and h is the total read density between the positions at which d1 and d2 occur). When comparing performance metrics between two experimental datasets, this filter was applied to both ChIP-seq datasets.

After masking the types of regions indicated above, we divided the whole genome or the entire chromosome into four granularities, 10 bp windows, 100 bp windows, 1,000 bp windows, and 10,000 bp windows. After collecting the sum of the read counts from experimental data and imputed data in each window, we compared the relationship between two datasets using four statistics: Pearson correlation, Spearman correlation, MAD, and JSD. Windows with 0 counts were removed from estimates of Pearson and Spearman correlation when using 10kb windows, as large regions without any ChIP-seq signal were likely driven by mappability issues.

To evaluate the accuracy of dHIT, we computed alternative performance metrics including MSE quantification at different subsets of genomic sites, as well as ROC and PRC curves for the recovery of peak calls. We added precision recall curves (PRC) following the setup introduced by Nair et. al. (see ref^129^), in which we divided the holdout chromosome into 500 bp non-overlapping windows from which we exacted ground truth labels using cell type specific peak calls generated by ENCODE. We generated PRCs or ROC curves by thresholding the imputed histone modification signal intensity to divide the same windows into those predicted to be enriched/ not enriched for each histone mark. To provide additional context for the PRC or ROC curves, we also computed PRC/ ROC curves in the same manner from experimental data. All analyses focus on the holdout chromosome (chr21) in the holdout cell type (GM12878).

We computed mean-squared errors (MSE) following performance metrics similar to those presented by Durham et. al. and Schreiber et. al. (see ref^44, 57^). We computed MSE in different genomic regions, including the top 1% of imputed windows (MSEimp); and the top 1% of experimental windows (MSEobs), two independent definitions of promoter and enhancer, using either proximity to gene annotations (GENCODE) or the stability of the transcription unit produced by each annotation following the nomenclature detailed in ref^11^.

### ChromHMM analysis

Chromatin state annotations were generated using ChromHMM^69^. We used the 18 state core model (model_18_core_K27ac) trained using ENCODE data^55^, because we had already imputed all of the histone modifications used in this model. To convert imputed histone modifications into data that met the requirements of ChromHMM, we fit the sum of imputed signal in 200 bp windows to a Poisson distribution, and identified windows with values higher than the 0.999th quantile. Chromatin segmentation was performed using the *MakeSegmentation* command, following the instructions from the authors^70^. We also made chromatin segmentations using an alternative source of experimental data for six histone marks, including H3K27ac, H3K27me3, H3K36me3, H3K4m1, H3K4me3, and H3K9me3 from ENCODE and other sources, as outlined in **Supplementary Table 1**. Chromatin segmentations were compared between experimental datasets, and between imputed and experimental data, using the Jaccard distance between each pair of states^130^. All computations were performed with bedtools^115^. When comparing enrichments of each state to those expected at random, we randomized the position of each state using bedtools random.

### Predicting bivalent TSSs

Bivalent genes in mESCs were identified using data from ref ^65^ and converted into mm9 coordinates using liftOver. Bivalent transcription start sites were predicted using a random forest. We used features representing H3K4me3 within 1,000 bp in 250 bp bins and H3K27me3 within 60,000 bp in 15,000 bp bins surrounding each promoter. All imputed histone modification data was based on models trained in K562 cells. We trained on a matched set of 100 bivalent and 100 non-bivalent promoters. The model was tested on a random set of 100 bivalent and 100 non-bivalent promoters that excluded promoters held out during training.

### Classification of H3K27me3 distribution

We obtained data from 86 H3K27me3 datasets from the Roadmap Epigenome Project (Data sources listed in **Supplementary Table 4**). Data from each sample was classified using a systematic approach designed to represent the degree to which each sample appeared to fit either the broad or punctate distribution of H3K27me3. Briefly, data from chromosome 21 was split into 10kb non-overlapping bins. The amount of H3K27me3 signal was counted in each bin. Bins from each sample were placed in descending order based on the read counts in that bin. The top and bottom 0.5% of bins were removed from each dataset and data was normalized to the total number of reads. Finally, we conducted a principal component analysis. We confirmed by manual inspection that principle component 1, accounting for 95.56% of the variance in the data, corresponded to the degree to which each sample showed a “punctate” or “broad” pattern. The value of principal component 1 in each sample was used in downstream analyses as a surrouge for the punctate and broad pattern. To compare the differences in patterns through differentiation, we manually categorized each of the 86 datasets as either pluripotent, multipotent, fetal, or adult/ somatic primary cells. We compared values of principal component 1 across these groups using a two-sided Wilcoxon rank sum test in R.

## Acknowledgements

We thank XSEDE allocation number TG-MCB160061 as well as an NVIDIA GPU Grant for providing computational resources required in this study. We thank James Lewis, Haiyuan Yu, Anniina Vihervaara, Mike DeBerardine, and all members of the Danko and Lis labs for valuable discussions and suggestions. Work in this publication was supported by R01-HG009309 (NHGRI) to CGD and a grant from the Zweig Memorial Fund for Equine Research to DFA and CGD, and by NIH grant RO1GM25232 to JTL. DFA is an Investigator of the Dorothy Russell Havemeyer Foundations, Inc. The content is solely the responsibility of the authors and does not necessarily represent the official views of the US National Institutes of Health. Some of the figures in this manuscript were created using BioRender. The dHIT software package is provided at https://github.com/Danko-Lab/histone-mark-imputation.

