## Supplementary Notes for "Interdependence between histone marks and steps in Pol II transcription"

**Supplementary Note 1: Types of errors common to dHIT.**

We examined systematic errors observed between experimental and dHIT imputed ChIP-seq data. Several types of recurring error we noticed are described in this Supplementary Note.

- **Poor prediction in background regions:** Several quality control metrics show evidence of a weak correlation between dHIT and ChIP-seq signal in background windows. Errors in background reflect small differences in the predicted and experimental read counts at individual genomic positions in background regions, which add up when summed over large window sizes. Errors in background signal can be observed clearly in the lower-left quadrant in scatter plots comparing experimental and imputed data (see **Supplementary Fig. 1**). Also, whereas predictions made using other ChIP-seq experimental data results in mean squared error that is significantly lower genome-wide than in peak regions (Durham et al., 2018; Schreiber et al., 2020) (due to the relative ease of predicting background signals using other ChIP-seq datasets), dHIT mean squared error is about the same genome wide or in peak regions (see **Supplementary Fig. 5**).

We think the most likely explanation is that these errors reflect differences in the way ChIP-seq and PRO-seq assays capture background. In ChIP-seq, there is substantial background pulldown of DNA due to non-specific binding of DNA to beads, tubes, tips or other sources of contamination. This background signal varies across the genome due to a variety of technical and biological factors (e.g., mappability; copy number alterations in some cell lines; etc.). In PRO-seq, the background is generally much lower and distributed in a very different way than ChIP-seq. Differences in the background distribution reflect fundamental differences in the assays: PRO-seq signal is derived from RNA (ChIP-seq is DNA), and PRO-seq has less background signal because the assay has three affinity purification steps (ChIP-seq has one). Differences in the background distribution between assays make it more difficult for dHIT to predict the number of ChIP-seq reads in regions that do not have much signal, especially in larger window sizes.

- **Mismapping in blacklist regions:** ENCODE blacklist regions had lower quality control metrics. Previous work has shown that ChIP-seq signal is not reliable in blacklist regions (Amemiya et al., 2019). Blacklist regions were excluded from all analyses and therefore did not affect quality control metrics.
- **Regions of focal amplification:** dHIT frequently predicted lower signal than observed in experimental data in regions with high copy number. This error was particularly noticeable in genome browser tracks (but also affected other quality control metrics) in cell types with abnormal karyotypes (e.g., K562). Our intuition is that this error occurred because increased DNA content is more difficult to detect using PRO-seq than ChIP-seq signal, due to fundamental differences in the way both assays capture background and signal.
- **Differences in the distribution of ChIP-seq signal within peaks:** Although dHIT captured most of the variation within experimental ChIP-seq peaks, the imputed signal was often spread over slightly larger regions than experimental signal (see especially **Fig. 1C**, **Supplementary Fig. 4, Supplementary Fig. 7**). This may indicate either systematic biological variation in the distance between Pol II and marked nucleosomes or it may reflect uncertainty in the model due to noise in either PRO-seq or ChIP-seq assays.
- **Clear disagreement between imputed and experimental data:** We identified a handful of cases where there were clearly defined peaks in the imputed histone modification data, but not the experimental data (or vice versa). Many of these examples are located in intergenic regions, and cannot be explained by signal in gene bodies or other adjacent regulatory elements. An outstanding example of this type of error at the *CERK* promoter is shown in **Supplementary Fig. 12A**. We found that these differences between dHIT imputed and ENCODE data were not reproducible in experimental data collected by our own lab in stocks of K562 cells that closely match those used for PRO-seq (**Supplementary Fig. 12B**). This implies that these systematic differences most likely reflect biological differences in cell stocks, handling, or other environmental conditions between cell lines.
