## Supplementary Figures and Tables for "Interdependence between histone marks and steps in Pol II transcription"

| ID | Data Type | Cell type/ Samp | Organism | GEO/ dbGaP ID | Lab | Institution | Sequencing Dep | Training/ test | Reference |
|---|---|---|---|---|---|---|---|---|---|
| | GRO-seq | mESC | Mouse | GSE48895 | Lis | Cornell University | | Test | Elife 2014 Apr 29;3:e02407. PMID: 24843027 |
| | leChRO-seq | GBM-05-16 | Human | phs001646.v1.p1 | Danko | Cornell University | 12026819 | Test | Nat Genet. 2018 Nov;50(11):1553-1564. PMID: 30349114 |
| | leChRO-seq | GBM-07-05 | Human | phs001646.v1.p1 | Danko | Cornell University | 33477124 | Test | Nat Genet. 2018 Nov;50(11):1553-1564. PMID: 30349114 |
| | leChRO-seq | GBM-05-17 | Human | phs001646.v1.p1 | Danko | Cornell University | 37121068 | Test | Nat Genet. 2018 Nov;50(11):1553-1564. PMID: 30349114 |
| | leChRO-seq | GBM-05-23 | Human | phs001646.v1.p1 | Danko | Cornell University | 38750059 | Test | Nat Genet. 2018 Nov;50(11):1553-1564. PMID: 30349114 |
| | leChRO-seq | GBM-05-05 | Human | phs001646.v1.p1 | Danko | Cornell University | 31814017 | Test | Nat Genet. 2018 Nov;50(11):1553-1564. PMID: 30349114 |
| | leChRO-seq | GBM-07-07 | Human | phs001646.v1.p1 | Danko | Cornell University | 53694384 | Test | Nat Genet. 2018 Nov;50(11):1553-1564. PMID: 30349114 |
| | leChRO-seq | GBM-05-35 | Human | phs001646.v1.p1 | Danko | Cornell University | 18865872 | Test | Nat Genet. 2018 Nov;50(11):1553-1564. PMID: 30349114 |
| | leChRO-seq | GBM-97-04 | Human | phs001646.v1.p1 | Danko | Cornell University | 10593102 | Test | Nat Genet. 2018 Nov;50(11):1553-1564. PMID: 30349114 |
| | ChRO-seq | GBM-15-90 | Human | phs001646.v1.p1 | Danko | Cornell University | 150711728 | Test | Nat Genet. 2018 Nov;50(11):1553-1564. PMID: 30349114 |
| | leChRO-seq | GBM-05-15 | Human | phs001646.v1.p1 | Danko | Cornell University | 21731177 | Test | Nat Genet. 2018 Nov;50(11):1553-1564. PMID: 30349114 |
| | leChRO-seq | GBM-07-02 | Human | phs001646.v1.p1 | Danko | Cornell University | 10303802 | Test | Nat Genet. 2018 Nov;50(11):1553-1564. PMID: 30349114 |
| | leChRO-seq | GBM-05-30 | Human | phs001646.v1.p1 | Danko | Cornell University | 26041476 | Test | Nat Genet. 2018 Nov;50(11):1553-1564. PMID: 30349114 |
| | leChRO-seq | GBM-05-18 | Human | phs001646.v1.p1 | Danko | Cornell University | 34744863 | Test | Nat Genet. 2018 Nov;50(11):1553-1564. PMID: 30349114 |
| | leChRO-seq | GBM-05-21 | Human | phs001646.v1.p1 | Danko | Cornell University | 27920964 | Test | Nat Genet. 2018 Nov;50(11):1553-1564. PMID: 30349114 |
| | leChRO-seq | GBM-05-33 | Human | phs001646.v1.p1 | Danko | Cornell University | 25874121 | Test | Nat Genet. 2018 Nov;50(11):1553-1564. PMID: 30349114 |
| | leChRO-seq | GBM-05-45 | Human | phs001646.v1.p1 | Danko | Cornell University | 21405989 | Test | Nat Genet. 2018 Nov;50(11):1553-1564. PMID: 30349114 |
| | leChRO-seq | GBM-06-12 | Human | phs001646.v1.p1 | Danko | Cornell University | 19037300 | Test | Nat Genet. 2018 Nov;50(11):1553-1564. PMID: 30349114 |
| | ChRO-seq / leCh | GBM-88-04 | Human | phs001646.v1.p1 | Danko | Cornell University | 22477998 | Test | Nat Genet. 2018 Nov;50(11):1553-1564. PMID: 30349114 |
| | leChRO-seq | GBM-05-26 | Human | phs001646.v1.p1 | Danko | Cornell University | 11581790 | Test | Nat Genet. 2018 Nov;50(11):1553-1564. PMID: 30349114 |
| | leChRO-seq | GBM-06-05 | Human | phs001646.v1.p1 | Danko | Cornell University | 32896583 | Test | Nat Genet. 2018 Nov;50(11):1553-1564. PMID: 30349114 |
| G1 | PRO-seq | K562 | Human | GSM1480327 | Lis | Cornell University | 374946808 | Training | Nat Genet. 2014 Dec;46(12):1311-20. PMID: 25383968 |
| G2 | GRO-seq | K562 | Human | GSM1480325 | Lis | Cornell University | 18129333 | Training | Nat Genet. 2014 Dec;46(12):1311-20. PMID: 25383968 |
| G3 | PRO-seq | K562 | Human | GSM3452725 | Danko | Cornell University | 57520888 | Training | Genome Res. 2019 Feb;29(2):293-303. PMID: 30573452 |
| G5 | PRO-seq | K562 | Human | GSM2361442,GSM2361443 | Sistonen | Åbo Akademi University | 71452942 | Training | Nat Commun. 2017 Aug 15;8(1):255. PMID: 28811569 |
| G6 | PRO-seq | K562 | Human | GSM2545324 | Lis | Cornell University | 26972822 | Training | Genome Res. 2017 Nov;27(11):1816-1829. PMID: 29025894 |
| G7 | PRO-seq | K562 | Human | GSM2545325 | Lis | Cornell University | 27046373 | Test | Genome Res. 2017 Nov;27(11):1816-1829. PMID: 29025894 |
| G8 | GRO-seq | K562 | Human | GSM1622612,GSM1622613,GSM1622614,GSM1622615 | Sistonen | Åbo Akademi University | 49159579 | Test | Genome Biol. 2015 Jul 28;16:153. PMID: 26259101 |
| GM | GRO-seq | GM12878 | Human | GSM1480326 | Lis | Cornell University | 105936649 | Test | Nat Genet. 2014 Dec;46(12):1311-20. PMID: 25383968 |
| GH | GRO-seq | HCT116 | Human | GSM1304424,GSM1304425 | Espinosa | University of Colorado, Boulder | 118728260 | Test | Elife. 2014 May 27;3:e02200. PMID: 24867637 |
| CD4 | PRO-seq | CD4 | Human | GSM2265095 | Danko | Cornell University | 33087414 | Test | Nat Ecol Evol. 2018 Mar;2(3):537-548. PMID: 29379187 |
| H1 | GRO-seq | HELA | Human | GSE62046 | Lis | Cornell University | 5812515 | Test | Nat Commun. 2014 Nov 12;5:5336. PMID: 25387874 |
| | ChRO-seq | Liver | Horse | | Danko | Cornell University | | Test | Herein |

**Supplementary Table 1: PRO-seq/ ChRO-seq data used in the present study.**

| Histone Mark | Cell type | Organism | GEO/ SRA ID | Training/ test | Lab | Institution | Reference | Antibody | Fragmentation type |
|---|---|---|---|---|---|---|---|---|---|
| H3K4me2 | K562 | Human | GSM733651 | Training | Bernstein | ENCODE Broad | Nature 2012 Sep 6;489(7414):57-74. PMID: 22955616 | ab7766 | Sonication, 200-700 bp |
| H3K4me3 | K562 | Human | GSM733680 | Training | Bernstein | ENCODE Broad | Nature 2012 Sep 6;489(7414):57-74. PMID: 22955616 | ab8580; 07-473 | Sonication, 200-700 bp |
| H3K9ac | K562 | Human | GSM733778 | Training | Bernstein | ENCODE Broad | Nature 2012 Sep 6;489(7414):57-74. PMID: 22955616 | ab4441 | Sonication, 200-700 bp |
| H3K27ac | K562 | Human | GSM733656 | Training | Bernstein | ENCODE Broad | Nature 2012 Sep 6;489(7414):57-74. PMID: 22955616 | ab4729 | Sonication, 200-700 bp |
| H3K122ac | K562 | Human | GSM2054695 | Training | Bickmore | University of Essex, Sch | Nat Genet 2016 Jun;48(6):681-6. PMID: 27089178 | Custom (Tropberger | MNase, Mononucleosomes |
| H3K4me1 | K562 | Human | GSM733692 | Training | Bernstein | ENCODE Broad | Nature 2012 Sep 6;489(7414):57-74. PMID: 22955616 | ab8895 | Sonication, 200-700 bp |
| H3K36me3 | K562 | Human | GSM733714 | Training | Bernstein | ENCODE Broad | Nature 2012 Sep 6;489(7414):57-74. PMID: 22955616 | ab9050 | Sonication, 200-700 bp |
| H4K20me1 | K562 | Human | GSM733675 | Training | Bernstein | ENCODE Broad | Nature 2012 Sep 6;489(7414):57-74. PMID: 22955616 | ab9051 | Sonication, 200-700 bp |
| H3K27me3 | K562 | Human | GSM733658 | Training | Bernstein | ENCODE Broad | Nature 2012 Sep 6;489(7414):57-74. PMID: 22955616 | 07-449 | Sonication, 200-700 bp |
| H3K9me3 | K562 | Human | GSM733776 | Training | Bernstein | ENCODE Broad | Nature 2012 Sep 6;489(7414):57-74. PMID: 22955616 | ab8898 | Sonication, 200-700 bp |
| H3K4me2 | GM12878 | Human | GSM733769 | Testing | Bernstein | ENCODE Broad | Nature 2012 Sep 6;489(7414):57-74. PMID: 22955616 | ab7766 | Sonication, 200-700 bp |
| H3K4me3 | GM12878 | Human | GSM733708 | Testing | Bernstein | ENCODE Broad | Nature 2012 Sep 6;489(7414):57-74. PMID: 22955616 | ab8580; 07-473 | Sonication, 200-700 bp |
| H3K9ac | GM12878 | Human | GSM733677 | Testing | Bernstein | ENCODE Broad | Nature 2012 Sep 6;489(7414):57-74. PMID: 22955616 | ab4441 | Sonication, 200-700 bp |
| H3K27ac | GM12878 | Human | GSM733771 | Testing | Bernstein | ENCODE Broad | Nature 2012 Sep 6;489(7414):57-74. PMID: 22955616 | ab4729 | Sonication, 200-700 bp |
| H3K4me1 | GM12878 | Human | GSM733772 | Testing | Bernstein | ENCODE Broad | Nature 2012 Sep 6;489(7414):57-74. PMID: 22955616 | ab8895 | Sonication, 200-700 bp |
| H3K36me3 | GM12878 | Human | GSM733679 | Testing | Bernstein | ENCODE Broad | Nature 2012 Sep 6;489(7414):57-74. PMID: 22955616 | ab9050 | Sonication, 200-700 bp |
| H4K20me1 | GM12878 | Human | GSM733642 | Testing | Bernstein | ENCODE Broad | Nature 2012 Sep 6;489(7414):57-74. PMID: 22955616 | ab9051 | Sonication, 200-700 bp |
| H3K27me3 | GM12878 | Human | GSM733758 | Testing | Bernstein | ENCODE Broad | Nature 2012 Sep 6;489(7414):57-74. PMID: 22955616 | 07-449 | Sonication, 200-700 bp |
| H3K9me3 | GM12878 | Human | GSM733664 | Testing | Bernstein | ENCODE Broad | Nature 2012 Sep 6;489(7414):57-74. PMID: 22955616 | ab8898 | Sonication, 200-700 bp |
| H3K4me2 | HCT116 | Human | GSE96381 | Testing | Bernstein | ENCODE Broad | Nature 2012 Sep 6;489(7414):57-74. PMID: 22955616 | Milipore, 07-030 | Sonication, 200-700 bp |
| H3K4me3 | HCT116 | Human | GSE96123 | Testing | Bernstein | ENCODE Broad | Nature 2012 Sep 6;489(7414):57-74. PMID: 22955616 | Miliport, 07-473; Abc | Sonication, 200-700 bp |
| H3K9ac | HCT116 | Human | GSE95916 | Testing | Bernstein | ENCODE Broad | Nature 2012 Sep 6;489(7414):57-74. PMID: 22955616 | ab4441 | Sonication, 200-700 bp |
| H3K27ac | HCT116 | Human | GSE96299 | Testing | Bernstein | ENCODE Broad | Nature 2012 Sep 6;489(7414):57-74. PMID: 22955616 | Active motif, 39133 | Sonication, 200-700 bp |
| H3K4me1 | HCT116 | Human | GSE95958 | Testing | Bernstein | ENCODE Broad | Nature 2012 Sep 6;489(7414):57-74. PMID: 22955616 | ab8895 | Sonication, 200-700 bp |
| H3K36me3 | HCT116 | Human | GSE95914 | Testing | Bernstein | ENCODE Broad | Nature 2012 Sep 6;489(7414):57-74. PMID: 22955616 | ab9050 | Sonication, 200-700 bp |
| H4K20me1 | HCT116 | Human | GSE96185 | Testing | Bernstein | ENCODE Broad | Nature 2012 Sep 6;489(7414):57-74. PMID: 22955616 | ab9051 | Sonication, 200-700 bp |
| H3K27me3 | HCT116 | Human | GSE86755 | Testing | Bernstein | ENCODE Broad | Nature 2012 Sep 6;489(7414):57-74. PMID: 22955616 | Milipore, 07-449 | Sonication, 200-700 bp |
| H3K9me3 | HCT116 | Human | GSE95968 | Testing | Bernstein | ENCODE Broad | Nature 2012 Sep 6;489(7414):57-74. PMID: 22955616 | ab8898 | Sonication, 200-700 bp |
| H3K4me2 | HeLa | Human | GSM733734 | Testing | Bernstein | ENCODE Broad | Nature 2012 Sep 6;489(7414):57-74. PMID: 22955616 | ab7766 | Sonication, 200-700 bp |
| H3K4me3 | HeLa | Human | GSM733682 | Testing | Bernstein | ENCODE Broad | Nature 2012 Sep 6;489(7414):57-74. PMID: 22955616 | ab8580; 07-473 | Sonication, 200-700 bp |
| H3K9ac | HeLa | Human | GSM733756 | Testing | Bernstein | ENCODE Broad | Nature 2012 Sep 6;489(7414):57-74. PMID: 22955616 | ab4441 | Sonication, 200-700 bp |
| H3K27ac | HeLa | Human | GSM733684 | Testing | Bernstein | ENCODE Broad | Nature 2012 Sep 6;489(7414):57-74. PMID: 22955616 | ab4729 | Sonication, 200-700 bp |
| H3K4me1 | HeLa | Human | GSM798322 | Testing | Bernstein | ENCODE Broad | Nature 2012 Sep 6;489(7414):57-74. PMID: 22955616 | ab8895 | Sonication, 200-700 bp |
| H3K36me3 | HeLa | Human | GSM733711 | Testing | Bernstein | ENCODE Broad | Nature 2012 Sep 6;489(7414):57-74. PMID: 22955616 | ab9050 | Sonication, 200-700 bp |
| H4K20me1 | HeLa | Human | GSM733689 | Testing | Bernstein | ENCODE Broad | Nature 2012 Sep 6;489(7414):57-74. PMID: 22955616 | ab9051 | Sonication, 200-700 bp |
| H3K27me3 | HeLa | Human | GSM733696 | Testing | Bernstein | ENCODE Broad | Nature 2012 Sep 6;489(7414):57-74. PMID: 22955616 | 07-449 | Sonication, 200-700 bp |
| H3K9me3 | HeLa | Human | GSM1003480 | Testing | Bernstein | ENCODE Broad | Nature 2012 Sep 6;489(7414):57-74. PMID: 22955616 | ab8898 | Sonication, 200-700 bp |
| H3K4me3 | CD4+ T-cells | Human | SRR001414,SRR001 | Testing | Zhao K | NIH | Cell 2007 129 (4): 823–37. PMID: 17512414 | ab8580 | Sonication, 200-300 bp |
| H3K9ac | CD4+ T-cells | Human | SRR037853 | Testing | Zhao K | NIH | Cell. 2009 Sep 4;138(5):1019-31. PMID: 19698979 | Not reported | Sonication, 200-300 bp |
| H3K4me1 | CD4+ T-cells | Human | SRR001439,SRR001 | Testing | Zhao K | NIH | Cell 2007 129 (4): 823–37. PMID: 17512414 | ab8895 | Sonication, 200-300 bp |
| H3K36me3 | CD4+ T-cells | Human | SRR001392,SRR001 | Testing | Zhao K | NIH | Cell 2007 129 (4): 823–37. PMID: 17512414 | ab9050 | Sonication, 200-300 bp |
| H3K27me3 | CD4+ T-cells | Human | SRR001426,SRR001 | Testing | Zhao K | NIH | Cell 2007 129 (4): 823–37. PMID: 17512414 | Upstate 07-449 | Sonication, 200-300 bp |
| H3K9me3 | CD4+ T-cells | Human | SRR001422,SRR001 | Testing | Zhao K | NIH | Cell 2007 129 (4): 823–37. PMID: 17512414 | abcam-8898 | Sonication, 200-300 bp |
| H3k27ac | K562 | Human | GSM2309710 | Testing | Xu | UT Southwestern Medic | Nat Cell Biol 2017 Jun;19(6):626-638. PMID: 28504707 | Abcam ab4729 | Sonication, 500 bp |
| H3k27ac | K562 | Human | GSM2877104 | Testing | Yuan | UT Southwestern Medic | Nat Commun 2018 Mar 5;9(1):943. PMID: 29507293 | Abcam, ab4729, GR | Not specified |
| H3k27ac | K562 | Human | GSM2877103 | Testing | Yuan | UT Southwestern Medic | Nat Commun 2018 Mar 5;9(1):943. PMID: 29507293 | Abcam, ab4729, GR | Not specified |
| H3k27ac | K562 | Human | GSM2054696 | Testing | Bickmore | University of Essex, Sch | Nat Genet 2016 Jun;48(6):681-6. PMID: 27089178 | Abcam ab4729, GR | MNase, Mononucleosomes |
| H3k4me1 | K562 | Human | GSM2054697 | Testing | Bickmore | University of Essex, Sch | Nat Genet 2016 Jun;48(6):681-6. PMID: 27089178 | Abcam ab8895, GR | MNase, Mononucleosomes |
| H3k27ac | K562 | Human | GSM646434 | Testing | Bernstein | Broad Institute | Nature 2011 May 5;473(7345):43-9. PMID: 21441907 | ab4729 | Sonication, 250 bp |
| H3k27ac | K562 | Human | GSM646435 | Testing | Bernstein | Broad Institute | Nature 2011 May 5;473(7345):43-9. PMID: 21441907 | ab4729 | Sonication, 250 bp |
| H3k27me3 | K562 | Human | GSM646436 | Testing | Bernstein | Broad Institute | Nature 2011 May 5;473(7345):43-9. PMID: 21441907 | 07-449 | Sonication, 250 bp |
| H3k27me3 | K562 | Human | GSM646437 | Testing | Bernstein | Broad Institute | Nature 2011 May 5;473(7345):43-9. PMID: 21441907 | 07-449 | Sonication, 250 bp |
| H3k36me3 | K562 | Human | GSM646438 | Testing | Bernstein | Broad Institute | Nature 2011 May 5;473(7345):43-9. PMID: 21441907 | ab9050 | Sonication, 250 bp |
| H3k36me3 | K562 | Human | GSM646439 | Testing | Bernstein | Broad Institute | Nature 2011 May 5;473(7345):43-9. PMID: 21441907 | ab9050 | Sonication, 250 bp |
| H3k4me1 | K562 | Human | GSM646440 | Testing | Bernstein | Broad Institute | Nature 2011 May 5;473(7345):43-9. PMID: 21441907 | ab8895 | Sonication, 250 bp |
| H3k4me1 | K562 | Human | GSM646441 | Testing | Bernstein | Broad Institute | Nature 2011 May 5;473(7345):43-9. PMID: 21441907 | ab8895 | Sonication, 250 bp |
| H3k4me2 | K562 | Human | GSM646442 | Testing | Bernstein | Broad Institute | Nature 2011 May 5;473(7345):43-9. PMID: 21441907 | ab7766 | Sonication, 250 bp |
| H3k4me2 | K562 | Human | GSM646443 | Testing | Bernstein | Broad Institute | Nature 2011 May 5;473(7345):43-9. PMID: 21441907 | ab7766 | Sonication, 250 bp |
| H3k4me3 | K562 | Human | GSM646444 | Testing | Bernstein | Broad Institute | Nature 2011 May 5;473(7345):43-9. PMID: 21441907 | 07-473 | Sonication, 250 bp |
| H3k4me3 | K562 | Human | GSM646445 | Testing | Bernstein | Broad Institute | Nature 2011 May 5;473(7345):43-9. PMID: 21441907 | 07-473 | Sonication, 250 bp |
| H3k9ac | K562 | Human | GSM646446 | Testing | Bernstein | Broad Institute | Nature 2011 May 5;473(7345):43-9. PMID: 21441907 | ab4441 | Sonication, 250 bp |
| H3k9ac | K562 | Human | GSM646447 | Testing | Bernstein | Broad Institute | Nature 2011 May 5;473(7345):43-9. PMID: 21441907 | ab4441 | Sonication, 250 bp |
| H3k9me3 | K562 | Human | GSM646448 | Testing | Bernstein | Broad Institute | Nature 2011 May 5;473(7345):43-9. PMID: 21441907 | ab8898 | Sonication, 250 bp |
| H3k9me3 | K562 | Human | GSM646449 | Testing | Bernstein | Broad Institute | Nature 2011 May 5;473(7345):43-9. PMID: 21441907 | ab8898 | Sonication, 250 bp |
| H4k20me1 | K562 | Human | GSM646450 | Testing | Bernstein | Broad Institute | Nature 2011 May 5;473(7345):43-9. PMID: 21441907 | ab9051 | Sonication, 250 bp |
| H4k20me1 | K562 | Human | GSM646451 | Testing | Bernstein | Broad Institute | Nature 2011 May 5;473(7345):43-9. PMID: 21441907 | ab9051 | Sonication, 250 bp |
| H3k27ac | GM12878 | Human | GSM646316 | Testing | Bernstein | Broad Institute | Nature 2011 May 5;473(7345):43-9. PMID: 21441907 | ab4729 | Sonication, 250 bp |
| H3k27ac | GM12878 | Human | GSM646317 | Testing | Bernstein | Broad Institute | Nature 2011 May 5;473(7345):43-9. PMID: 21441907 | ab4729 | Sonication, 250 bp |
| H3k27me3 | GM12878 | Human | GSM646318 | Testing | Bernstein | Broad Institute | Nature 2011 May 5;473(7345):43-9. PMID: 21441907 | 07-449 | Sonication, 250 bp |
| H3k27me3 | GM12878 | Human | GSM646319 | Testing | Bernstein | Broad Institute | Nature 2011 May 5;473(7345):43-9. PMID: 21441907 | 07-449 | Sonication, 250 bp |
| H3k36me3 | GM12878 | Human | GSM646320 | Testing | Bernstein | Broad Institute | Nature 2011 May 5;473(7345):43-9. PMID: 21441907 | ab9050 | Sonication, 250 bp |
| H3k36me3 | GM12878 | Human | GSM646321 | Testing | Bernstein | Broad Institute | Nature 2011 May 5;473(7345):43-9. PMID: 21441907 | ab9050 | Sonication, 250 bp |
| H3k4me1 | GM12878 | Human | GSM646322 | Testing | Bernstein | Broad Institute | Nature 2011 May 5;473(7345):43-9. PMID: 21441907 | ab8895 | Sonication, 250 bp |
| H3k4me1 | GM12878 | Human | GSM646323 | Testing | Bernstein | Broad Institute | Nature 2011 May 5;473(7345):43-9. PMID: 21441907 | ab8895 | Sonication, 250 bp |
| H3k4me2 | GM12878 | Human | GSM646324 | Testing | Bernstein | Broad Institute | Nature 2011 May 5;473(7345):43-9. PMID: 21441907 | ab7766 | Sonication, 250 bp |
| H3k4me2 | GM12878 | Human | GSM646325 | Testing | Bernstein | Broad Institute | Nature 2011 May 5;473(7345):43-9. PMID: 21441907 | ab7766 | Sonication, 250 bp |
| H3k4me3 | GM12878 | Human | GSM646326 | Testing | Bernstein | Broad Institute | Nature 2011 May 5;473(7345):43-9. PMID: 21441907 | 07-473 | Sonication, 250 bp |
| H3k4me3 | GM12878 | Human | GSM646327 | Testing | Bernstein | Broad Institute | Nature 2011 May 5;473(7345):43-9. PMID: 21441907 | 07-473 | Sonication, 250 bp |
| H3k9ac | GM12878 | Human | GSM646328 | Testing | Bernstein | Broad Institute | Nature 2011 May 5;473(7345):43-9. PMID: 21441907 | ab4441 | Sonication, 250 bp |
| H3k9ac | GM12878 | Human | GSM646329 | Testing | Bernstein | Broad Institute | Nature 2011 May 5;473(7345):43-9. PMID: 21441907 | ab4441 | Sonication, 250 bp |
| H4k20me1 | GM12878 | Human | GSM646330 | Testing | Bernstein | Broad Institute | Nature 2011 May 5;473(7345):43-9. PMID: 21441907 | ab9051 | Sonication, 250 bp |
| H4k20me1 | GM12878 | Human | GSM646331 | Testing | Bernstein | Broad Institute | Nature 2011 May 5;473(7345):43-9. PMID: 21441907 | ab9051 | Sonication, 250 bp |
| H3k27ac | K562 | Human | GSM1782704 | Testing | Bock | CeMM Research Cente | Nat Methods 2015 Oct;12(10):963-965. PMID: 26280331 | Diagenode pAb-196 | Sonication, 200-700 bp |
| H3k36me3 | K562 | Human | GSM1782705 | Testing | Bock | CeMM Research Cente | Nat Methods 2015 Oct;12(10):963-965. PMID: 26280331 | Diagenode pAb-192 | Sonication, 200-700 bp |
| H3k4me1 | K562 | Human | GSM1782706 | Testing | Bock | CeMM Research Cente | Nat Methods 2015 Oct;12(10):963-965. PMID: 26280331 | Diagenode pAb-194 | Sonication, 200-700 bp |
| H3k4me1 | K562 | Human | GSM1782707 | Testing | Bock | CeMM Research Cente | Nat Methods 2015 Oct;12(10):963-965. PMID: 26280331 | Diagenode pAb-194 | Sonication, 200-700 bp |
| H3k27ac | K562 | Human | GSM1782738 | Testing | Bock | CeMM Research Cente | Nat Methods 2015 Oct;12(10):963-965. PMID: 26280331 | Diagenode pAb-196 | Sonication, 200-700 bp |
| H3k27me3 | K562 | Human | GSM1782739 | Testing | Bock | CeMM Research Cente | Nat Methods 2015 Oct;12(10):963-965. PMID: 26280331 | Millipore 07-449 | Sonication, 200-700 bp |
| H3K27me3 | K562 | Human | GSM1782740 | Testing | Bock | CeMM Research Cente | Nat Methods 2015 Oct;12(10):963-965. PMID: 26280331 | Millipore 07-449 | Sonication, 200-700 bp |
| H3K4me3 | K562 | Human | GSM1782741 | Testing | Bock | CeMM Research Cente | Nat Methods 2015 Oct;12(10):963-965. PMID: 26280331 | Diagenode C15410( | Sonication, 200-700 bp |
| H3K4me3 | K562 | Human | GSM1782742 | Testing | Bock | CeMM Research Cente | Nat Methods 2015 Oct;12(10):963-965. PMID: 26280331 | Diagenode C15410( | Sonication, 200-700 bp |
| H3K27ac | HCT116 | Human | GSM2916001 | Testing | Wu | Wuhan University | Nucleic Acids Res 2019 Mar 18;47(5):2349-2364. PMID: 30 | abcam, ab4729 | MNase, Mono/ di nucleosomes |
| H3K27ac | HCT116 | Human | GSM2916002 | Testing | Wu | Wuhan University | Nucleic Acids Res 2019 Mar 18;47(5):2349-2364. PMID: 30 | abcam, ab4729 | MNase, Mono/ di nucleosomes |
| H3K4me3 | HCT116 | Human | GSM2916031 | Testing | Wu | Wuhan University | Nucleic Acids Res 2019 Mar 18;47(5):2349-2364. PMID: 30 | Millipore, 2739249 | MNase, Mono/ di nucleosomes |
| H3K4me3 | HCT116 | Human | GSM2916032 | Testing | Wu | Wuhan University | Nucleic Acids Res 2019 Mar 18;47(5):2349-2364. PMID: 30 | Millipore, 2739249 | MNase, Mono/ di nucleosomes |
| H3K27me3 | HCT116 | Human | GSM2916037 | Testing | Wu | Wuhan University | Nucleic Acids Res 2019 Mar 18;47(5):2349-2364. PMID: 30 | 9733S | MNase, Mono/ di nucleosomes |
| H3K27me3 | HCT116 | Human | GSM2916038 | Testing | Wu | Wuhan University | Nucleic Acids Res 2019 Mar 18;47(5):2349-2364. PMID: 30 | 9733S | MNase, Mono/ di nucleosomes |
| H3K4me3 | mESC | Mouse | GSM307618 | Testing | Bernstein | Broad Institute | Nature 2007 Aug 2;448(7153):553-60. PMID: 17603471 | Abcam 8580 | Sonication, 200-700 bp |
| H3K27me3 | mESC | Mouse | GSM307619 | Testing | Bernstein | Broad Institute | Nature 2007 Aug 2;448(7153):553-60. PMID: 17603471 | Upstate 07-449 | Sonication, 200-700 bp |
| H3K4me3 | Liver (horse) | Horse | | Testing | Bellone | Diagenode | Genes 2019 PMID: 31861495 | C15410003 | Sonication |
| H3K27ac | Liver (horse) | Horse | | Testing | Bellone | Diagenode | Genes 2019 PMID: 31861495 | C15410196 | Sonication |
| H3K4me1 | Liver (horse) | Horse | | Testing | Bellone | Diagenode | Genes 2019 PMID: 31861495 | C15410194 | Sonication |
| H3K27me3 | Liver (horse) | Horse | | Testing | Bellone | Diagenode | Genes 2019 PMID: 31861495 | C15410195 | Sonication |
| H3K4me1 | K562 | Human | | Training | Danko | Cornell University | Herein | ab8895 | MNase, Mono/ di nucleosomes |
| H3K4me2 | K562 | Human | | Training | Danko | Cornell University | Herein | ab7766 | MNase, Mono/ di nucleosomes |
| H3K4me3 | K562 | Human | | Training | Danko | Cornell University | Herein | ab8580 | MNase, Mono/ di nucleosomes |
| H3K27ac | K562 | Human | | Training | Danko | Cornell University | Herein | ab4729 | MNase, Mono/ di nucleosomes |
| H3K36me3 | K562 | Human | | Training | Danko | Cornell University | Herein | ab9050 | MNase, Mono/ di nucleosomes |
| H3K79me3 | K562 | Human | | Training | Danko | Cornell University | Herein | ab2621 | MNase, Mono/ di nucleosomes |

**Supplementary Table 2: ChIP-seq data used in the present study.**

| ID | Data Type | Cell Type | Organism | GEO | File_mb | Read_type |
|---|---|---|---|---|---|---|
| G1-11 | PRO-seq | K562 | Human | | 84367.94 | single |
| G1-12 | PRO-seq | K562 | Human | | 82879.5 | single |
| G2 | PRO-seq | K562 | Human | GSM148032[illegible] | 7188.66 | single |
| G3 | PRO-seq | K562 | Human | GSM345272[illegible] | 25426.8 | single |
| G5-1 | PRO-seq | K562 | Human | GSM236144[illegible] | 29601.79 | single |
| G5-2 | PRO-seq | K562 | Human | GSM236144[illegible] | 9792 | single |
| G6 | PRO-seq | K562 | Human | GSM254532[illegible] | 9393.66 | single |
| G-7 | PRO-seq | K562 | Human | GSM254532[illegible] | 8699.25 | single |
| G-8 | GRO-seq | K562 | Human | GSM162261[illegible] | 18508.96 | single |
| GM | GRO-seq | GM12878 | Human | GSM148032[illegible] | 69603.81 | single |
| H1 | GRO-seq | HELA | Human | GSE62046 | 18869.88 | single |
| HCT | GRO-seq | HCT | Human | GSM130442[illegible] | 127433.73 | single |
| CD4 | PRO-seq | CD4+ T-cells | Human | GSM226509[illegible] | 12458.1 | single |
| horse liver | ChRO-seq | Liver | Horse | | 4841.07 | single |

| Reads_with_ | Uninformativ | Pct_uninform | Trimmed_re | Trim_loss_ra | Peak_adapte | Degradation_ |
|---|---|---|---|---|---|---|
| 213181221 | 6253894 | 2.4965 | 244247718 | 2.5 | 0 | 0.1953 |
| 209877962 | 6184089 | 2.513 | 239895976 | 2.51 | 0 | 0.1965 |
| 22296806 | 1041369 | 3.4296 | 29322367 | 3.43 | 0 | 0.6269 |
| 100035186 | 15715 | 0.0138 | 113876613 | 0.01 | 25 | 0.8826 |
| 123231282 | 795572 | 0.5946 | 133007914 | 0.59 | 36 | 0.5504 |
| 41611292 | 208165 | 0.4657 | 44495551 | 0.47 | 41 | 0.8861 |
| 37038939 | 256020 | 0.6055 | 42027780 | 0.61 | 27 | 0.8069 |
| 31294262 | 433383 | 1.1064 | 38738698 | 1.11 | 33 | 0.5704 |
| 27309115 | 0 | 0 | 80235991 | 0 | 4 | 0 |
| 199673166 | 4681680 | 2.2773 | 200900066 | 2.28 | 0 | 0.4796 |
| 63012915 | 650143 | 0.8229 | 78357782 | 0.82 | 19 | 0.8697 |
| 194562378 | 794155 | 0.1479 | 536252205 | 0.15 | 21 | 36.5419 |
| 41464339 | 400564 | 0.768 | 51753538 | 0.77 | 35 | 0.3248 |
| 107686290 | 539132 | 0.4775 | 112364111 | 0.48 | 27 | 0.2913 |

| Alignment_r | Total_efficie | Read_depth | Mitochondria | Maximum_re | Genome_siz | NRF |
|---|---|---|---|---|---|---|
| 80.44 | 78.43 | 16.21 | 4632216 | 100 | 3137161264 | 0.46 |
| 80.41 | 78.39 | 15.93 | 4534817 | 100 | 3137161264 | 0.47 |
| 63.09 | 60.93 | 2.65 | 826389 | 50 | 3137161264 | 0.69 |
| 14.81 | 14.81 | 2.74 | 236602 | 76 | 3137161264 | 0.85 |
| 51.01 | 50.71 | 5.36 | 700258 | 75 | 3137161264 | 0.75 |
| 53 | 52.76 | 3.07 | 301373 | 75 | 3137161264 | 0.83 |

| | | | | | | |
|---|---|---|---|---|---|---|
| 64.93 | 64.54 | 3.11 | 744220 | 76 | 3137161264 | 0.85 |
| 72.03 | 71.23 | 3.31 | 937956 | 76 | 3137161264 | 0.83 |
| 61.6 | 61.6 | 4.29 | 707980 | 51 | 3137161264 | 0.7 |
| 70.89 | 69.28 | 10.99 | 1617934 | 100 | 3137161264 | 0.44 |
| 64.64 | 64.1 | 4.78 | 1289701 | 51 | 3137161264 | 0.54 |
| 62.05 | 61.96 | 18.58 | 6336477 | 50 | 3137161264 | 0.11 |
| 74.41 | 73.84 | 5.51 | 25905 | 51 | 3137161264 | 0.28 |
| 2.28 | 2.27 | 9.87 | 4205 | 85 | 3137161264 | 0.02 |

| TSS_coding_ | TSS_non-cod | Pause_index | Plus_FRiP | Minus_FRiP | mRNA_conta | Time |
|---|---|---|---|---|---|---|
| 14.2 | 5.8 | 10.51 | 0.36 | 0.34 | 1.36 | 1 day, 7:38:4 |
| 14.2 | 5.8 | 10.59 | 0.36 | 0.34 | 1.36 | 1 day, 7:35:4 |
| 6.3 | 17.8 | 2.77 | 0.32 | 0.34 | 2.17 | 2:37:58 |
| 26.9 | 10.1 | 27.4 | 0.35 | 0.33 | 1.54 | 12:40:20 |
| 18.9 | 10.8 | 19.83 | 0.36 | 0.34 | 1.48 | 11:14:43 |
| 19.9 | 11.5 | 20.33 | 0.36 | 0.34 | 1.53 | 4:30:12 |
| 20.2 | 7.5 | 16.14 | 0.35 | 0.34 | 1.48 | 4:30:27 |
| 25 | 8.7 | 19.41 | 0.35 | 0.33 | 1.51 | 4:51:11 |
| 4.7 | 11.3 | 2.32 | 0.34 | 0.35 | 1.73 | 10:36:45 |
| 10.7 | 22 | 2.92 | 0.33 | 0.36 | 1.47 | 15:21:14 |
| 9.2 | 23.8 | 2.9 | 0.33 | 0.35 | 1.58 | 6:07:49 |
| 5.3 | 9 | 2.84 | 0.3 | 0.31 | 1.3 | 1 day, 14:46: |
| 66.2 | 2.4 | 36.37 | 0.36 | 0.34 | 1.24 | 4:37:31 |
| 5.1 | 7.5 | 39.69 | 0.25 | 0.26 | 3 | 4:51:39 |

| Genome | Raw_reads | Fastq_reads |
|---|---|---|
| hg19 | 250501612 | 250501612 |
| hg19 | 246080065 | 246080065 |
| hg19 | 30363736 | 30363736 |
| hg19 | 113892328 | 113892328 |
| hg19 | 133803486 | 133803486 |
| hg19 | 44703716 | 44703716 |
| hg19 | 42283800 | 42283800 |
| hg19 | 39172081 | 39172081 |
| hg19 | 80235991 | 80235991 |
| hg19 | 205581746 | 205581746 |
| hg19 | 79007925 | 79007925 |
| hg19 | 537046360 | 537046360 |
| hg19 | 52154102 | 52154102 |
| hg19 | 112903243 | 112903243 |

| Mapped_rea | QC_filtered_ | Aligned_reads |
|---|---|---|
| 241211008 | 44746176 | 196464832 |
| 236969113 | 44067726 | 192901387 |
| 27133808 | 8632894 | 18500914 |
| 83080315 | 66211849 | 16868466 |
| 114856678 | 47008980 | 67847698 |
| 38384194 | 14799606 | 23584588 |
| 40423549 | 13134469 | 27289080 |
| 37430713 | 9526784 | 27903929 |
| 74503646 | 25076738 | 49426908 |
| 192195295 | 49770108 | 142425187 |
| 73996259 | 23348753 | 50647506 |
| 524549967 | 191810902 | 332739065 |
| 50011527 | 11500301 | 38511226 |
| 15804196 | 13242824 | 2561372 |

| PBC1 | PBC2 | Unmapped_reads |
|---|---|---|
| 0.7 | 4.93 | 3036710 |
| 0.7 | 5 | 2926863 |
| 0.86 | 8.7 | 2188559 |
| 0.93 | 18.25 | 30796298 |
| 0.9 | 17.03 | 18151236 |
| 0.94 | 25.09 | 6111357 |

| | | |
|---|---|---|
| 0.94 | 21.96 | 1604231 |
| 0.93 | 19.27 | 1307985 |
| 0.87 | 9.24 | 5732345 |
| 0.69 | 5.1 | 8704771 |
| 0.77 | 6.01 | 4361523 |
| 0.44 | 2.78 | 11702238 |
| 0.58 | 3.15 | 1742011 |
| 0.21 | 2.24 | 96559915 |

| Success |
|---|
| 09-05-00:45:21 |
| 09-05-00:42:47 |
| 09-03-19:45:34 |
| 09-04-05:48:17 |
| 09-04-07:59:38 |
| 09-04-16:10:34 |
| 09-05-19:45:54 |
| 09-05-20:05:06 |
| 09-12-11:02:04 |
| 09-06-06:34:31 |
| 09-12-06:06:56 |
| 09-15-00:19:32 |
| 09-12-05:00:10 |
| 09-13-19:52:29 |

**Table 3: PEPPRO quality control results of the PROseq, GROseq, and ChRO-seq data used in the present study.**

GSM1102787_UW.CD3_Primary_Cells.H3K27me3.RO_01679.Histone.DS22927.wig.bw
GSM1112827_BI.hESC_Derived_CD56+_Ectoderm_Cultured_Cells.H3K27me3.DNA_Lib_3193.wig.bw
GSM1127134_UCSF-UBC.Breast_Fibroblast_Primary_Cells.H3K27me3.RM071.wig.bw
GSM1160192_UW.Fetal_Adrenal_Gland.H3K27me3.H-24800.Histone.DS23067.wig.bw
GSM428295_UCSF-UBC.H3K27me3.wig.bw
GSM433167_BI.H1.H3K27me3.Solexa-8039.wig.bw
GSM433167_BI.H3K27me3.wig.bw
GSM537613_BI.CD3_Primary_Cells.H3K27me3.CD3_39661.wig.bw
GSM537627_BI.ES-I3.H3K27me3.Solexa-12678.wig.bw
GSM537631_BI.CD3_Primary_Cells.H3K27me3.CD3_39804.wig.bw
GSM537648_BI.ES-I3.H3K27me3.Solexa-10213-15380.wig.bw
GSM537657_BI.ES-WA7.H3K27me3.Solexa-12609.wig.bw
GSM537674_BI.iPS-18c.H3K27me3.Solexa-19791.wig.bw
GSM537698_BI.Adult_Liver.H3K27me3.3.wig.bw
GSM537700_BI.iPS-20b.H3K27me3.Solexa-15159.wig.bw
GSM537707_BI.Adult_Liver.H3K27me3.5.wig.bw
GSM605308_UCSD.H1.H3K27me3.YL95.wig.bw
GSM613815_UCSF-UBC.CD8_Naive_Primary_Cells.H3K27me3.TC001.wig.bw
GSM613872_UCSF-UBC.Breast_Myoepithelial_Cells.H3K27me3.RM066.wig.bw
GSM613887_UCSF-UBC.Breast_Myoepithelial_Cells.H3K27me3.RM080.wig.bw
GSM621393_BI.Fetal_Brain.H3K27me3.UW_H-22510.wig.bw
GSM621416_BI.iPS-20b.H3K27me3.Lib_MC_20100119_06--ChIP_MC_20100113_06_hiPS-20b_H3K27Me3.wig.bw
GSM621420_BI.Adipose_Derived_Mesenchymal_Stem_Cell_Cultured_Cells.H3K27me3.1.wig.bw
GSM621433_BI.iPS-15b.H3K27me3.Lib_MC_20100128_05--ChIP_MC_20100125_05_hiPS-15b_H3K27Me3.wig.bw
GSM621454_BI.Bone_Marrow_Derived_Mesenchymal_Stem_Cell_Cultured_Cells.H3K27me3.57.wig.bw
GSM667622_UCSD.H9.H3K27me3.SK99.wig.bw
GSM669594_UCSF-UBC.Breast_vHMEC.H3K27me3.RM035.wig.bw
GSM669625_UCSF-UBC.Fetal_Brain.H3K27me3.HuFNSC-T.wig.bw
GSM669913_BI.Brain_Hippocampus_Middle.H3K27me3.112.wig.bw
GSM669915_BI.Brain_Anterior_Caudate.H3K27me3.112.wig.bw
GSM669916_BI.Chondrocytes_from_Bone_Marrow_Derived_Mesenchymal_Stem_Cell_Cultured_Cells.H3K27me3.5
9.wig.bw GSM669930_BI.Adipose_Nuclei.H3K27me3.92.wig.bw
GSM669953_BI.Brain_Substantia_Nigra.H3K27me3.112.wig.bw
GSM669957_BI.Bone_Marrow_Derived_Mesenchymal_Stem_Cell_Cultured_Cells.H3K27me3.60.wig.bw
GSM669959_BI.Chondrocytes_from_Bone_Marrow_Derived_Mesenchymal_Stem_Cell_Cultured_Cells.H3K27me3.5
7.wig.bw GSM670007_BI.Bone_Marrow_Derived_Mesenchymal_Stem_Cell_Cultured_Cells.H3K27me3.59.wig.bw
GSM670009_BI.Bone_Marrow_Derived_Mesenchymal_Stem_Cell_Cultured_Cells.H3K27me3.58.wig.bw
GSM670017_BI.Adipose_Nuclei.H3K27me3.95.wig.bw
GSM670035_BI.Adipose_Nuclei.H3K27me3.94.wig.bw
GSM693278_UCSF-UBC.Breast_Luminal_Epithelial_Cells.H3K27me3.RM080.wig.bw
GSM706066_UCSD.H9.H3K27me3.AK142.wig.bw
GSM706067_UCSD.iPS_DF_19.11.H3K27me3.SK230.wig.bw
GSM706068_UCSD.iPS_DF_6.9.H3K27me3.SK275.wig.bw
GSM707001_UCSF-UBC.Brain_Germinal_Matrix.H3K27me3.HuFGM01.wig.bw
GSM752968_UCSD.H1_BMP4_Derived_Mesendoderm_Cultured_Cells.H3K27me3.AK132.wig.bw
GSM752969_UCSD.H1_BMP4_Derived_Mesendoderm_Cultured_Cells.H3K27me3.AK148.wig.bw
GSM752970_UCSD.iPS_DF_19.11.H3K27me3.SK435.wig.bw
GSM752971_UCSD.iPS_DF_6.9.H3K27me3.SK433.wig.bw
GSM772744_BI.Mesenchymal_Stem_Cell_Derived_Adipocyte_Cultured_Cells.H3K27me3.93.wig.bw
GSM772771_BI.Adipose_Nuclei.H3K27me3.93.wig.bw
GSM772772_BI.Brain_Inferior_Temporal_Lobe.H3K27me3.112.wig.bw

GSM772773_BI.Chondrocytes_from_Bone_Marrow_Derived_Mesenchymal_Stem_Cell_Cultured_Cells.H3K27me3.60.wig.bw
GSM772788_BI.Chondrocytes_from_Bone_Marrow_Derived_Mesenchymal_Stem_Cell_Cultured_Cells.H3K27me3.58.wig.bw GSM772801_BI.H9_Derived_Neuronal_Progenitor_Cultured_Cells.H3K27me3.113.wig.bw
GSM772819_BI.Mesenchymal_Stem_Cell_Derived_Adipocyte_Cultured_Cells.H3K27me3.92.wig.bw
GSM772821_BI.Adipose_Derived_Mesenchymal_Stem_Cell_Cultured_Cells.H3K27me3.92.wig.bw
GSM772827_BI.Brain_Anterior_Caudate.H3K27me3.149.wig.bw
GSM772833_BI.Brain_Mid_Frontal_Lobe.H3K27me3.149.wig.bw
GSM772855_BI.CD4_Memory_Primary_Cells.H3K27me3.Donor_101_8_pooled_leukopaks_Jan_20_2011.wig.bw
GSM772871_BI.CD8_Naive_Primary_Cells.H3K27me3.Donor_100_7_pooled_leukopaks_Jan_7_2011.wig.bw
GSM772878_BI.CD8_Memory_Primary_Cells.H3K27me3.Donor_100_7_pooled_leukopaks_Jan_7_2011.wig.bw
GSM772900_BI.CD8_Naive_Primary_Cells.H3K27me3.Donor_101_8_pooled_leukopaks_Jan_20_2011.wig.bw
GSM772906_BI.CD4+_CD25-_CD45RO+_Memory_Primary_Cells.H3K27me3.Donor_62.wig.bw
GSM772911_BI.CD4+_CD25-_CD45RA+_Naive_Primary_Cells.H3K27me3.Donor_62.wig.bw
GSM772919_BI.CD4_Naive_Primary_Cells.H3K27me3.Donor_101_8_pooled_leukopaks_Jan_20_2011.wig.bw
GSM772935_BI.iPS-18a.H3K27me3.DNA_Lib_348.wig.bw
GSM772947_BI.CD4_Naive_Primary_Cells.H3K27me3.Donor_100_7_pooled_leukopaks_Jan_7_2011.wig.bw
GSM772956_BI.CD8_Memory_Primary_Cells.H3K27me3.Donor_101_8_pooled_leukopaks_Jan_20_2011.wig.bw
GSM772974_BI.Colon_Smooth_Muscle.H3K27me3.Donor_83_REMC_18.wig.bw
GSM772983_BI.Brain_Angular_Gyrus.H3K27me3.149.wig.bw
GSM772989_BI.Brain_Cingulate_Gyrus.H3K27me3.149.wig.bw
GSM772993_BI.Brain_Inferior_Temporal_Lobe.H3K27me3.149.wig.bw
GSM772998_BI.CD4_Memory_Primary_Cells.H3K27me3.Donor_100_7_pooled_leukopaks_Jan_7_2011.wig.bw
GSM806937_UCSF-UBC.Fetal_Brain.H3K27me3.HuFNSC02.wig.bw
GSM806945_UCSF-UBC.Fetal_Brain.H3K27me3.HuFNSC01.wig.bw
GSM817226_UCSF-UBC.Brain_Germinal_Matrix.H3K27me3.HuFGM02.wig.bw
GSM818033_UCSD.H1_Derived_Neuronal_Progenitor_Cultured_Cells.H3K27me3.AK221.wig.bw
GSM896165_UCSD.H1_Derived_Neuronal_Progenitor_Cultured_Cells.H3K27me3.AK320.wig.bw
GSM916038_BI.Brain_Hippocampus_Middle.H3K27me3.150.wig.bw
GSM916055_BI.Adipose_Nuclei.H3K27me3.7.wig.bw
GSM916061_BI.Fetal_Brain.H3K27me3.UW_H22676.wig.bw
GSM916062_BI.hESC_Derived_CD56+_Mesoderm_Cultured_Cells.H3K27me3.DNA_Lib_1274.wig.bw
GSM997227_BI.CD4+_CD25-_CD45RA+_Naive_Primary_Cells.H3K27me3.Donor_332.wig.bw
GSM997238_BI.CD4+_CD25-_CD45RO+_Memory_Primary_Cells.H3K27me3.Donor_332.wig.bw
GSM997247_BI.hESC_Derived_CD56+_Ectoderm_Cultured_Cells.H3K27me3.DNA_Lib_1807.wig.bw
GSM997259_BI.Colon_Smooth_Muscle.H3K27me3.156.wig.bw

**Table 4: Roadmap Epigenome H3K27me3 datasets used to draw Supplementary Figure 14.**

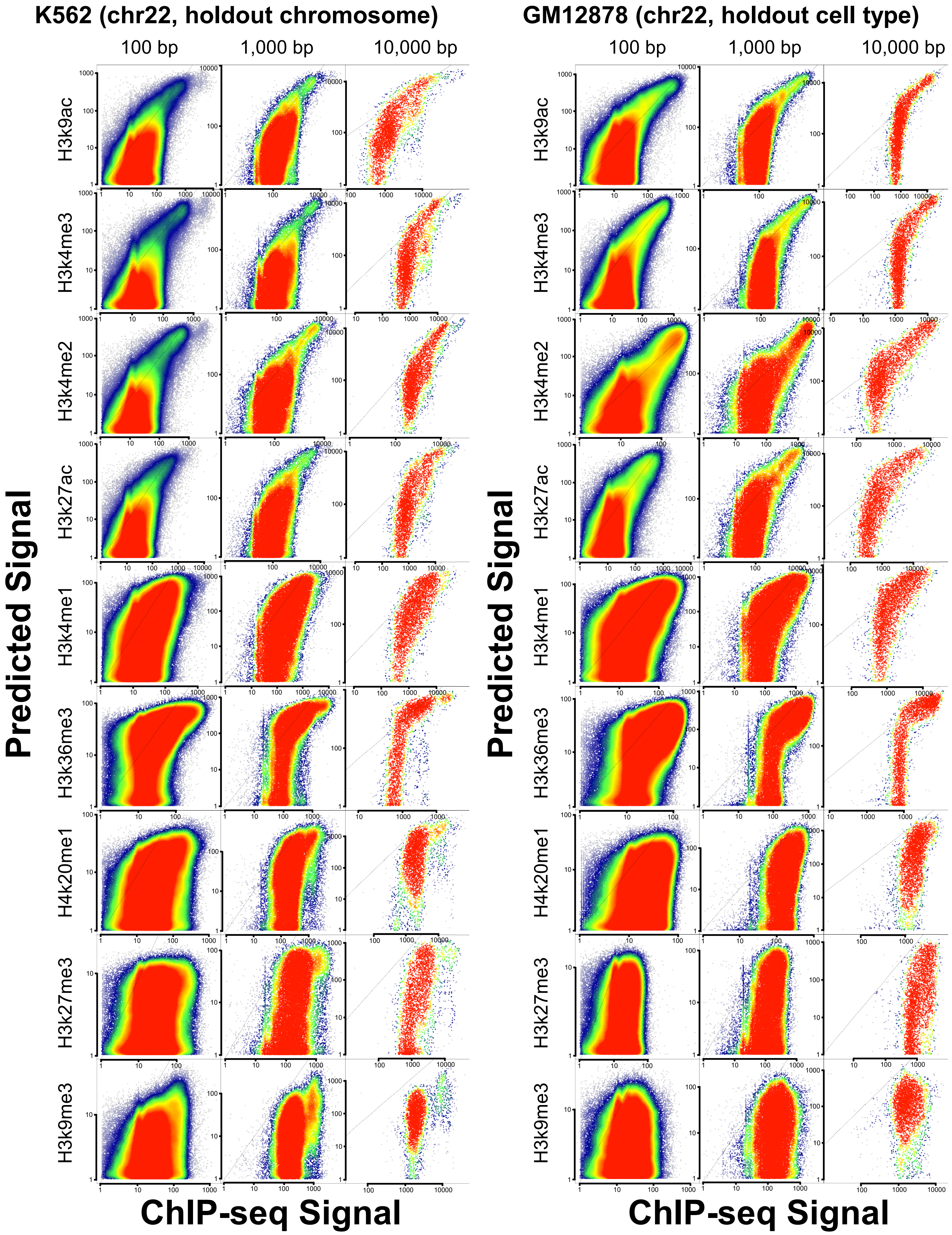


**Supplementary Figure 1. Imputation of histone marks using nascent transcription.**
Scatterplots show predicted (Y-axis) as a function of experimental ChIP-seq signal (X-axis) for ten different histone modifications in K562 and GM12878. Plots show correlations in a holdout chromosome (chr22) at three distinct length scales.

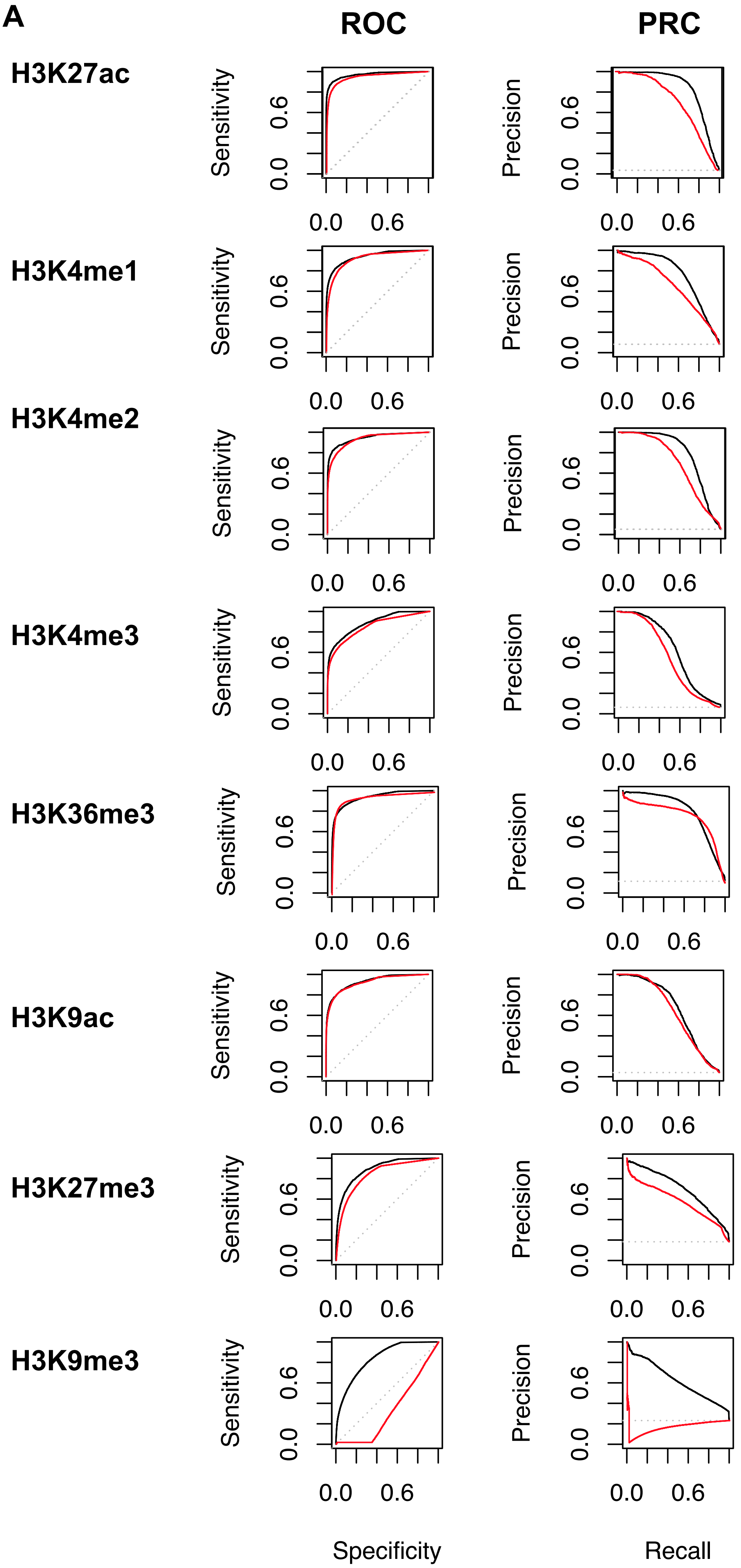


**B**

■ Experiment

■ Imputed

| | ROC | PRC |
|---|---|---|
| H3K27ac | 0.9693<br>0.9483 | 0.8299<br>0.7108 |
| H3K4me1 | 0.9371<br>0.9193 | 0.7635<br>0.6555 |
| H3K4me2 | 0.9456<br>0.9328 | 0.7898<br>0.6889 |
| H3K4me3 | 0.8864<br>0.8497 | 0.6223<br>0.5591 |
| H3K36me3 | 0.9371<br>0.9429 | 0.7992<br>0.7706 |
| H3K9ac | 0.9281<br>0.9196 | 0.6672<br>0.6395 |
| H3K27me3 | 0.8934<br>0.8448 | 0.7018<br>0.5783 |
| H3K9me3 | 0.8378<br>0.3335 | 0.6258<br>0.1714 |

**Supplementary Figure 2: Performance metrics to evaluate dHIT predictions.**
A. ROC and PRC plots describe the relationship between imputed and ENCODE ChIP-seq data within ENCODE peaks on chr21, holdout during dHIT training. B. Quantification of area und precision curves for both ROC and PRC plots in A.

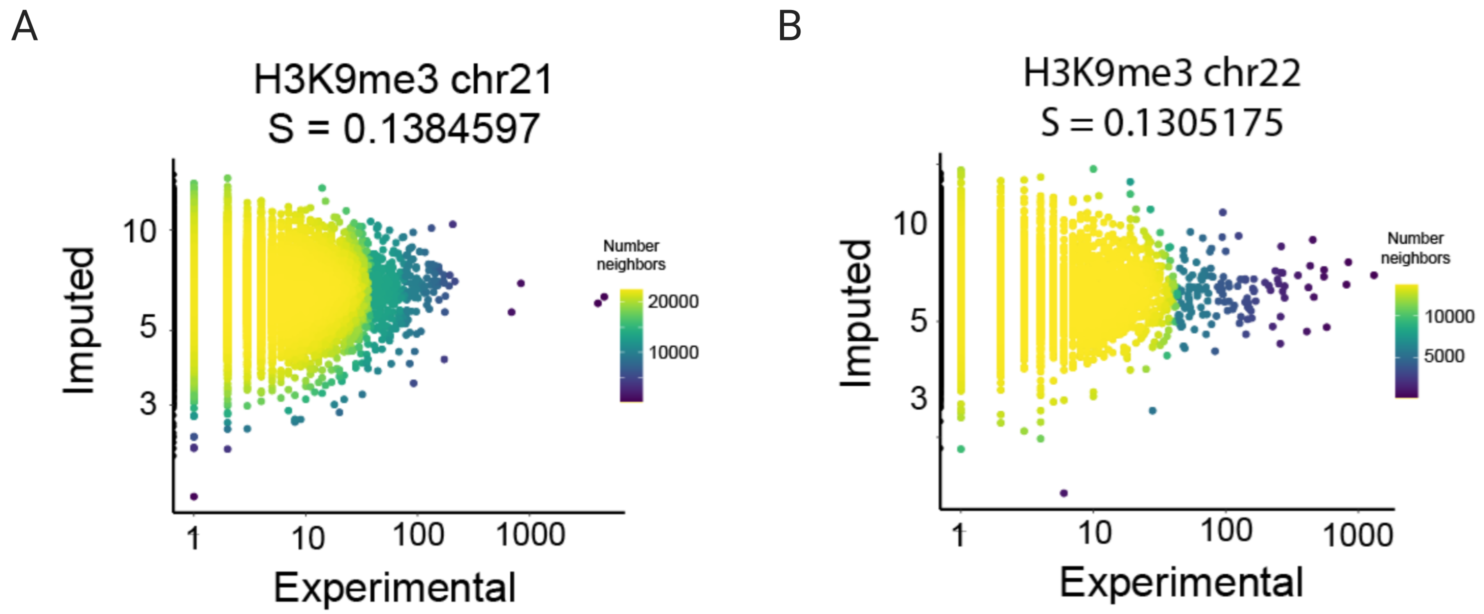


**Supplementary Figure 3. Correlation between CUT&TAG experimental and imputed H3K9me3.** Scatter plots depict imputed H3K9me3 (Y-axis) as a function of CUT&TAG experimental (X-axis) for H3K9me3 in K562. Spearman correlations were computed on the holdout chromosome chr21 (A) and chr22 (B).

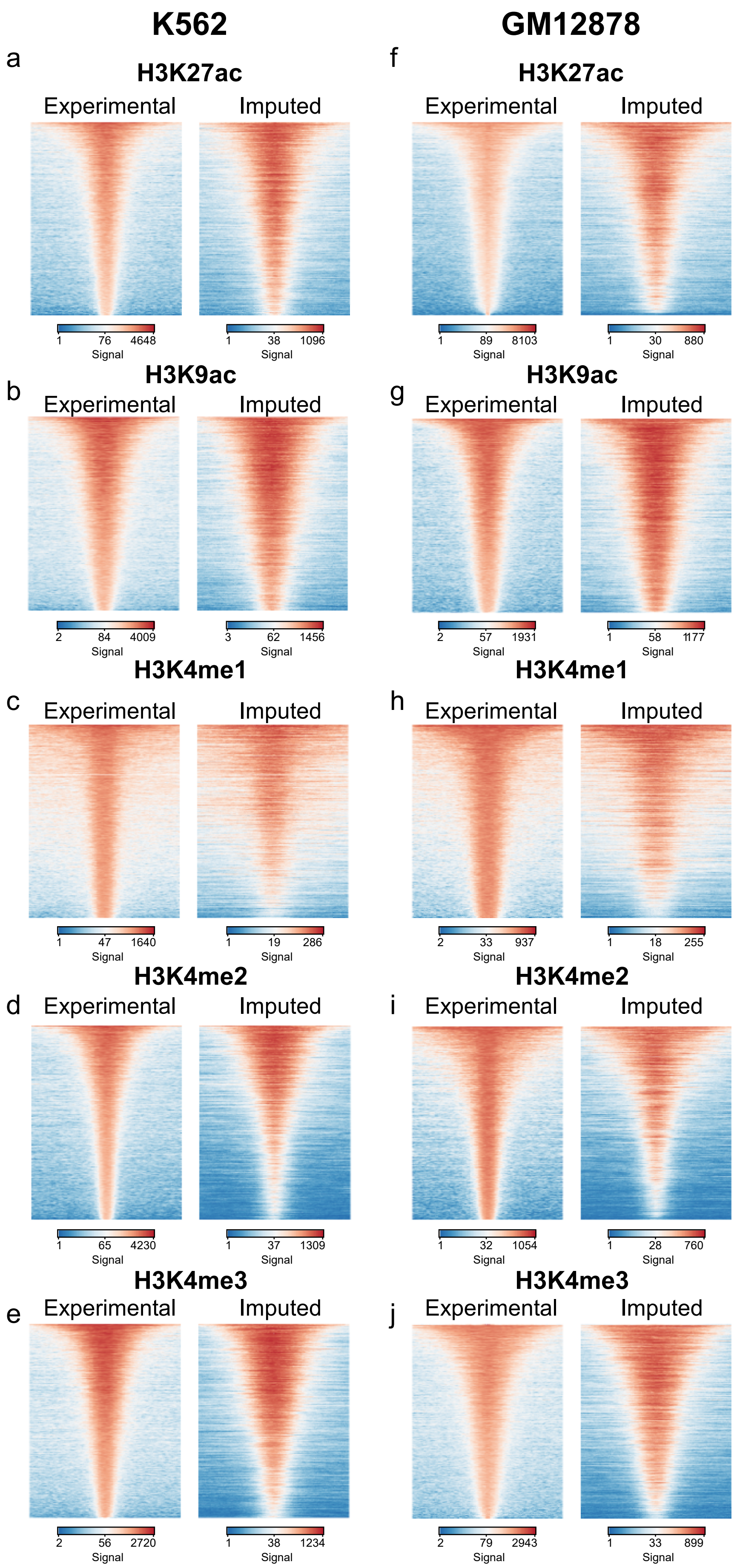


**Supplementary Figure 4. Comparison between experimental and imputed histone mark abundance.** Heatmaps show the experimental and imputed abundance of active, punctate histone marks in K562 (A-E) or GM12878 (F-J). Heatmaps show all peaks calls based on experimental ChIP-seq data ordered by the highest total signal intensity.

| | H3K27ac | H3K4me1 | H3K4me2 | H3K4me3 | H3K36me3 | H3K9ac | H3K27me3 | H3K9me3 | H3K20me1 |
|---|---|---|---|---|---|---|---|---|---|
| **MSE global** | 2.236 | 0.720 | 1.844 | 2.509 | 0.526 | 0.970 | 0.578 | 0.866 | 0.793 |
| **MSEobs** | 2.179 | 0.554 | 1.791 | 2.411 | 0.384 | 0.89 | 0.288 | 0.703 | 0.494 |
| **MSEimp** | 1.52 | 0.617 | 1.759 | 2.25 | 0.415 | 0.891 | 0.351 | 0.046 | 0.612 |
| **MSE (s/s, s/u, u/s)* transcripts** | 0.354 | 1.337 | 0.385 | 0.112 | 1.933 | 0.237 | 1.279 | 0.722 | 0.487 |
| **MSE (u/u)** transcripts** | 0.529 | 0.455 | 0.251 | 0.119 | 0.753 | 0.429 | 0.661 | 0.471 | 1.032 |
| **MSE near gene promoters** | 0.735 | 0.403 | 0.261 | 0.212 | 0.750 | 0.482 | 1.159 | 0.642 | 0.778 |
| **MSE at transcribed enhancers** | 1.054 | 0.531 | 0.387 | 0.474 | 0.820 | 0.747 | 2.734 | 0.781 | 1.032 |

Transcripts nomenclature as described in Core and Martins, *et al*, 2014.
*s/s = stable/stable; *s/u = stable/unstable; *u/s = unstable/stable; **u/u = unstable/unstable

**Supplementary Figure 5: Mean-squared error (MSE) quantification at different subsets of genomic sites in GM12878.**

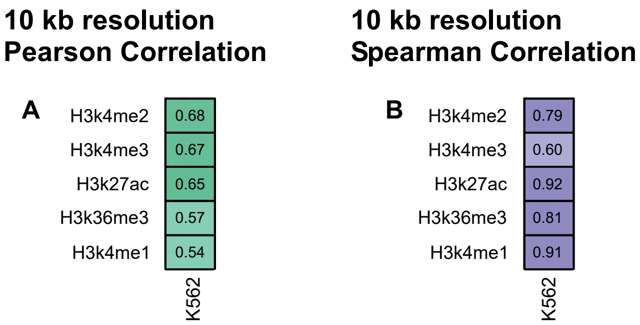


**Supplementary Figure 6. Correlation between imputed and experimental MNase ChIP-seq.** Heatmaps show the Pearson (A) and Spearman (B) correlations between predicted and experimental MNase ChIP-seq in 10kb windows on a holdout chromosome (chr22).

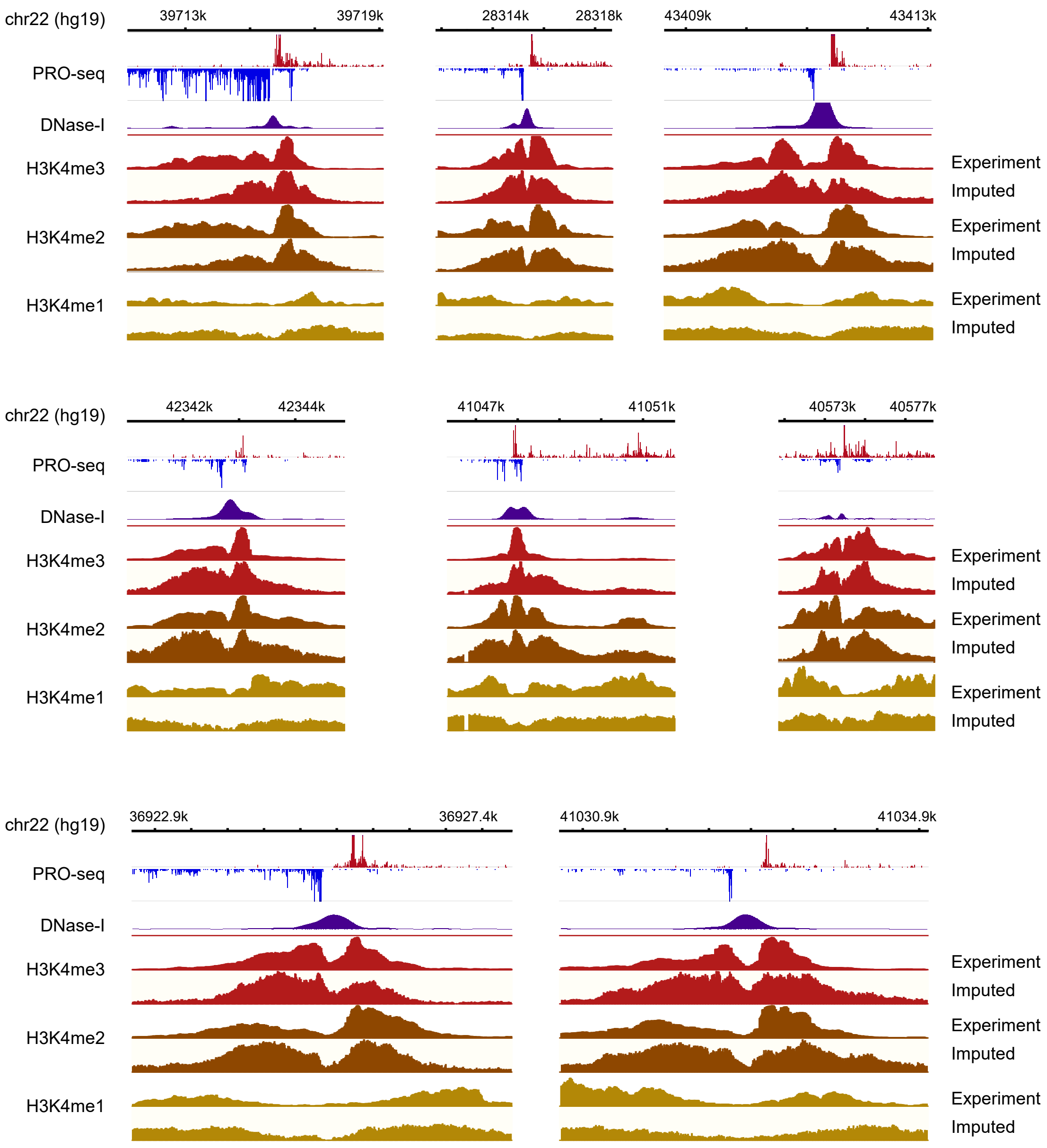


**Supplementary Figure 7. Comparison between experimental and imputed MNase ChIP-seq.** Genome-browser plots show the distribution of PRO-seq, DNase-I hypersensitivity signal, and the signal for H3K4me3, H3K4me2, and H3K4me1 derived from MNase ChIP-seq and imputation near 9 transcribed regions in K562 cells.

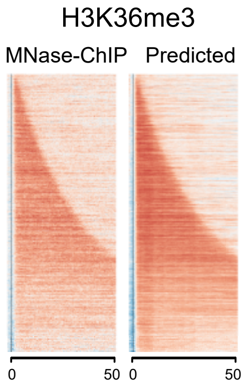


**Supplementary Figure 8. Heatmaps show the signal for H3K36me3.**
Heatmaps show MNase ChIP-seq and imputed signal intensity for H3K36me3, a gene body mark, deposited in the body of annotated genes. Genes are sorted by gene length.

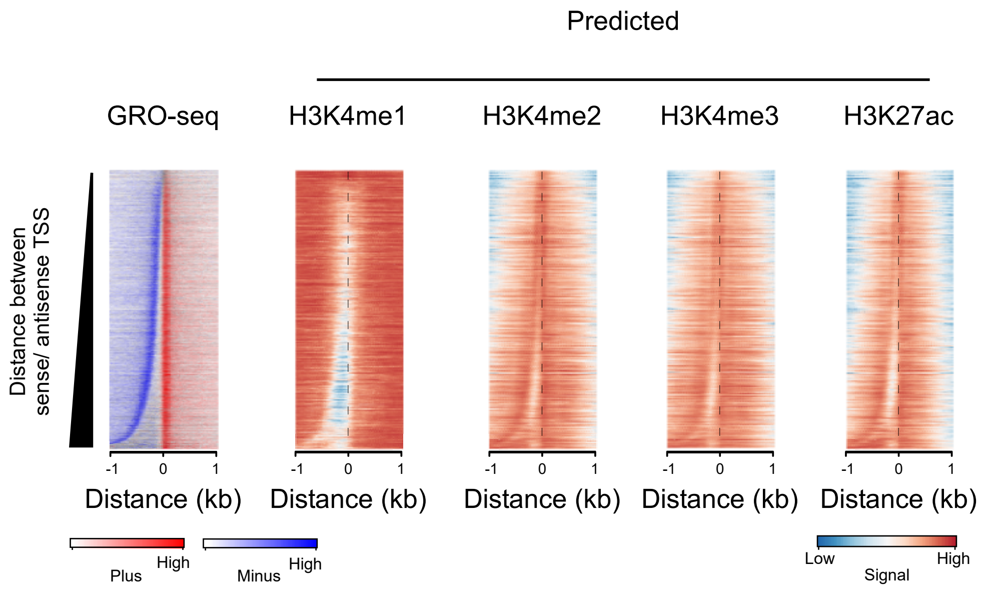


**Supplementary Figure 9. Heatmaps show histone marks near transcription initiation domains in holdout cell type, GM12878.**

Heatmaps show the distribution of transcription (left) and histone modifications (right) predicted using transcription. Rows represent transcription initiation domains in GM12878 cells defined using GRO-cap data by Core, Martins, et. al. (2014) Nat. Gen. Heatmaps were ordered by the distance between the most frequently used TSS in each transcription initiation domain on the plus and minus strand.

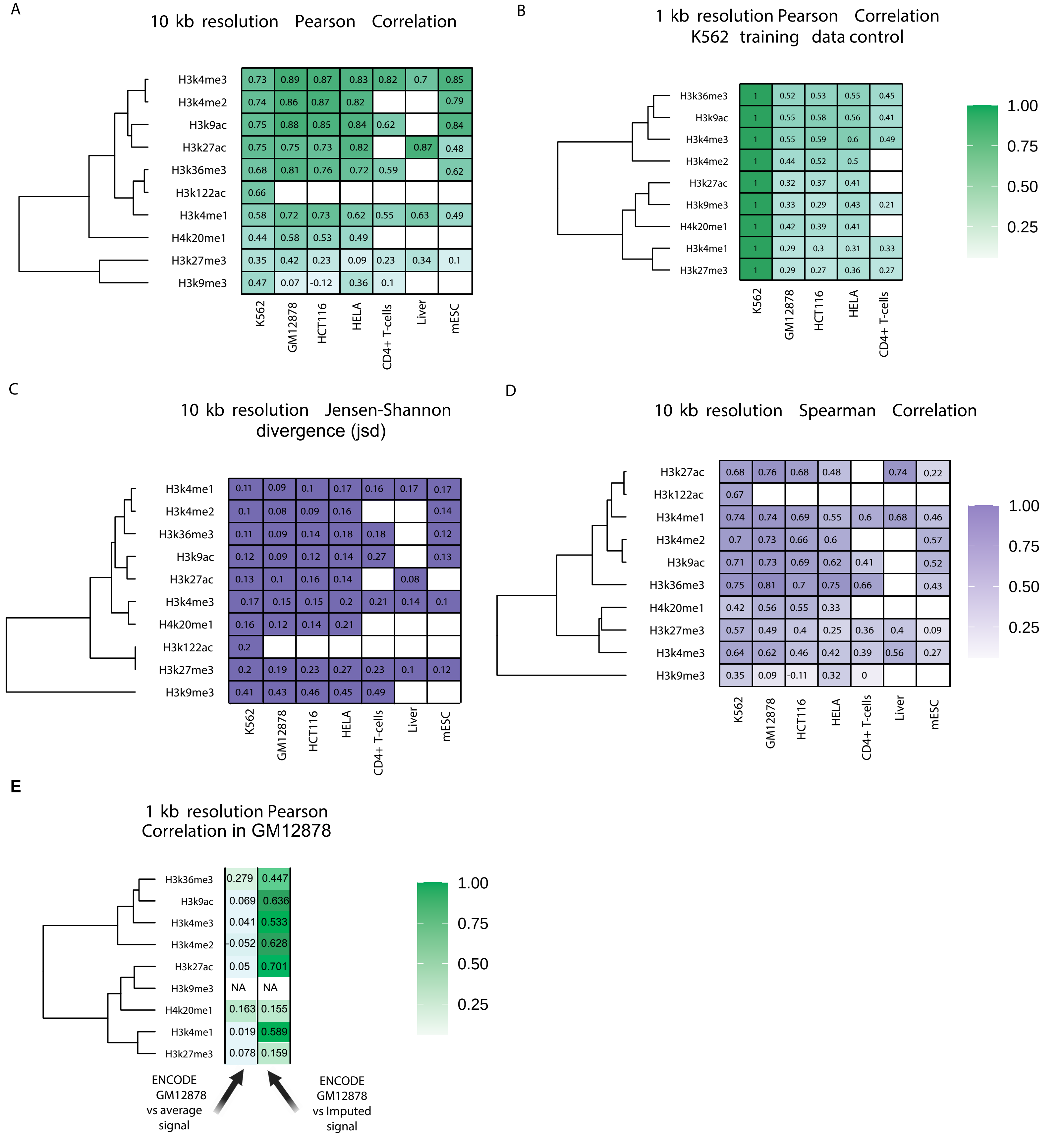


**Supplementary Figure 10. Evaluation of cross-cell line imputation by different metrics.**
(A-C) Heatmaps show Pearson's correlation (A), Spearman's rank correlation (B), Jensen-Shannon and divergence (C) between predicted and ChIP-seq measurements of nine histone modifications. Values are computed in 10kb windows on the holdout chromosome (chr22) in humans, chr1 in horse, and chr1 in mice. Empty cells indicate that no experimental data is available for comparison in the cell type shown.
(D) Heatmap shows Pearson's correlation between the training dataset in K562 cells and experimental data collected in the indicated human cell line. Values are computed in 1kb windows on the holdout chromosome (chr22) in humans.
(E) Heatmap shows Pearson's correlation between the ENCODE experimental data and either Imputed data or the average signal of the other human cell lines investigated. Values are computed in 1kb windows on the holdout chromosome (chr22) in GM12878.

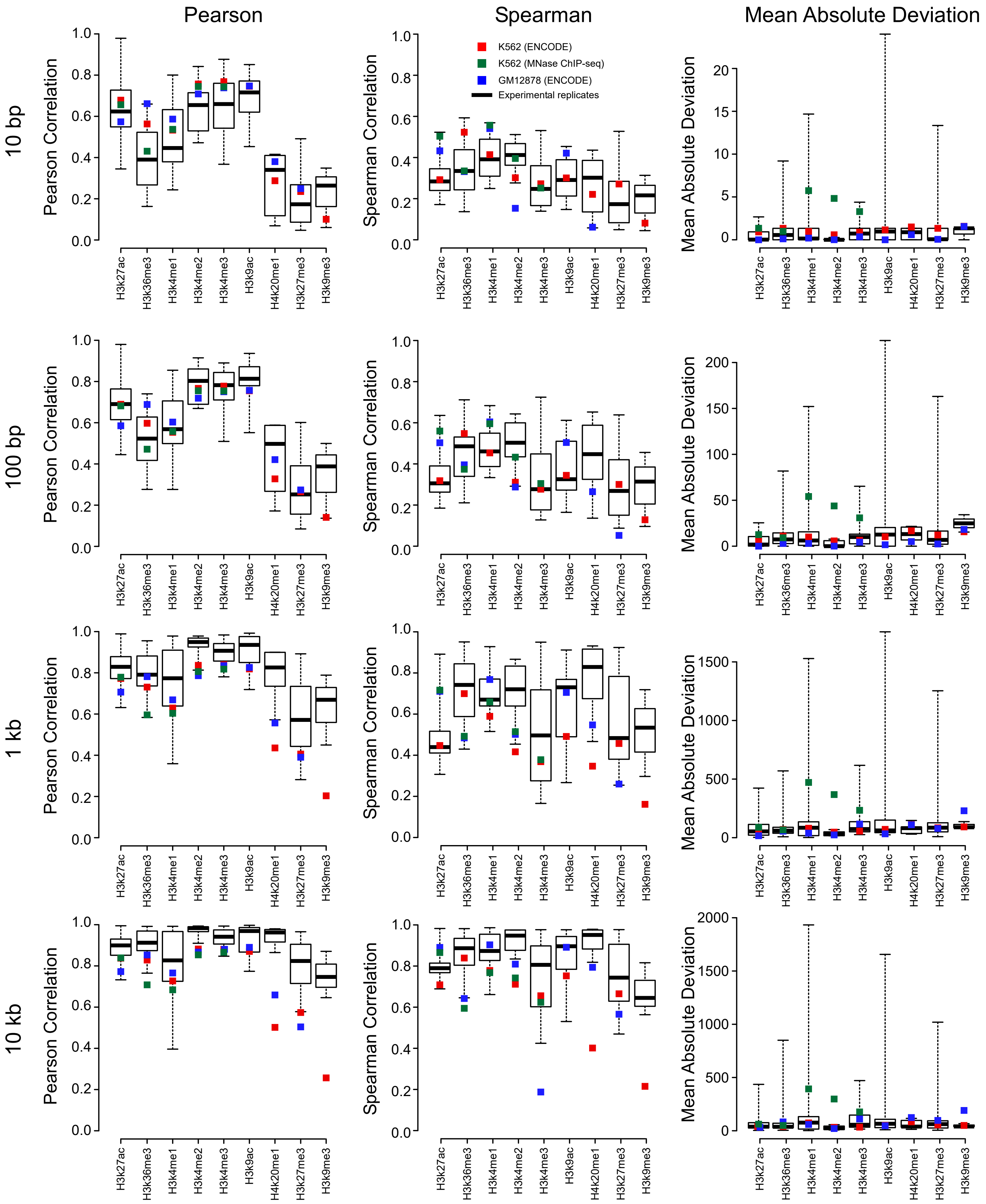


**Supplementary Figure 11. Comparison between imputation and multiple ChIP-seq experiments.**
Box and whiskers plot shows the Pearson correlation between different experimental datasets for six histone marks in K562 and GM12878.The correlation between data imputed in K562 and GM12878 and the ENCODE experimental data in the same cell line is shown respectively by red and blue squares. All values are computed on a holdout chromosome (chr22) not used during training.

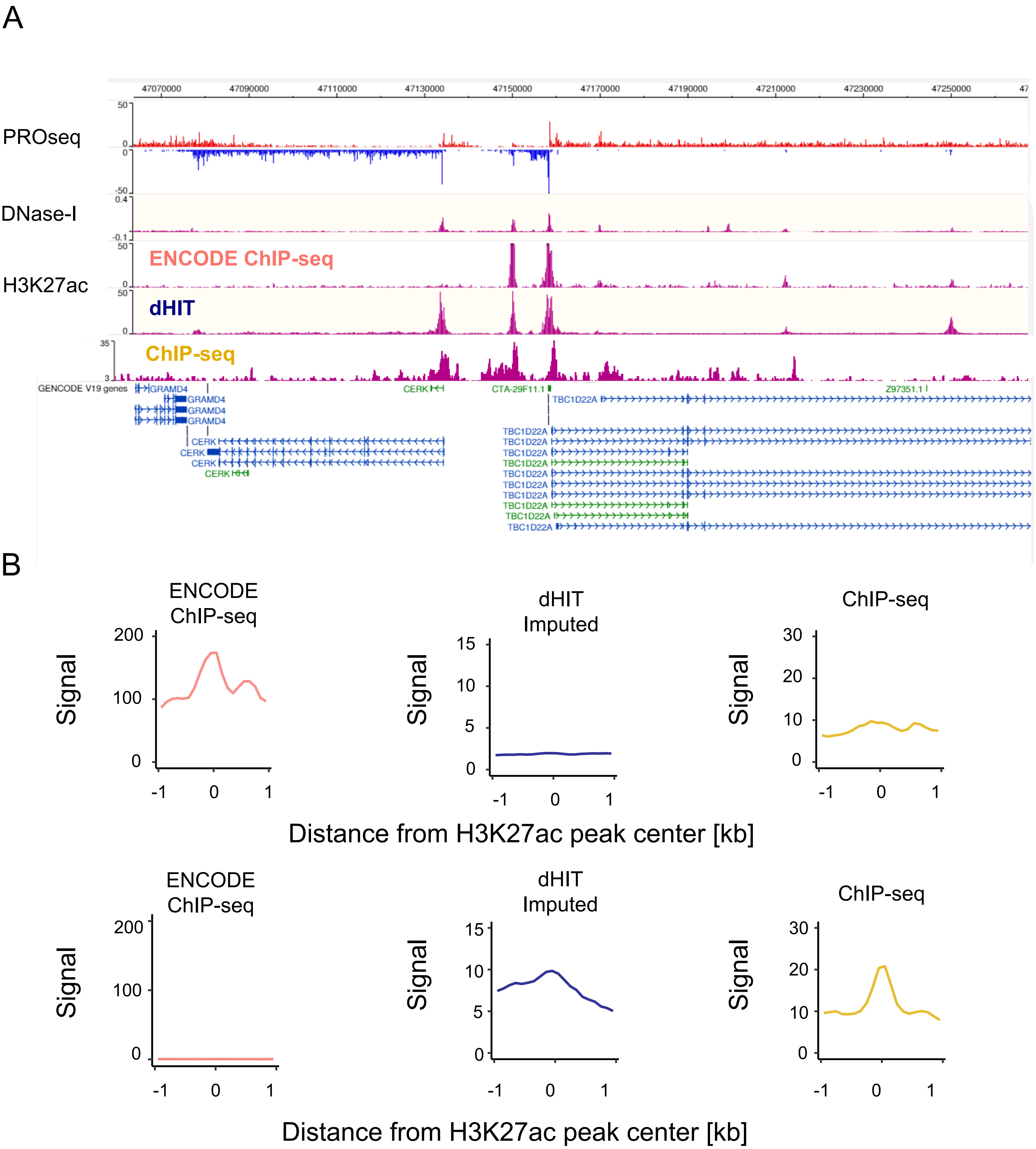


**Supplementary Figure 12. Comparison between ENCODE, imputed, and experimental ChIP-seq.**
(A) Browser shot shows the ENCODE, imputed, and experimental ChIP-seq signals at the CERK locus.
(B) Meta plots compare the H3K27ac content of two different sets of H3K27ac annotated peaks: peaks high in ENCODE signal and depleted in imputed ChIP (top) or vice-versa (bottom).

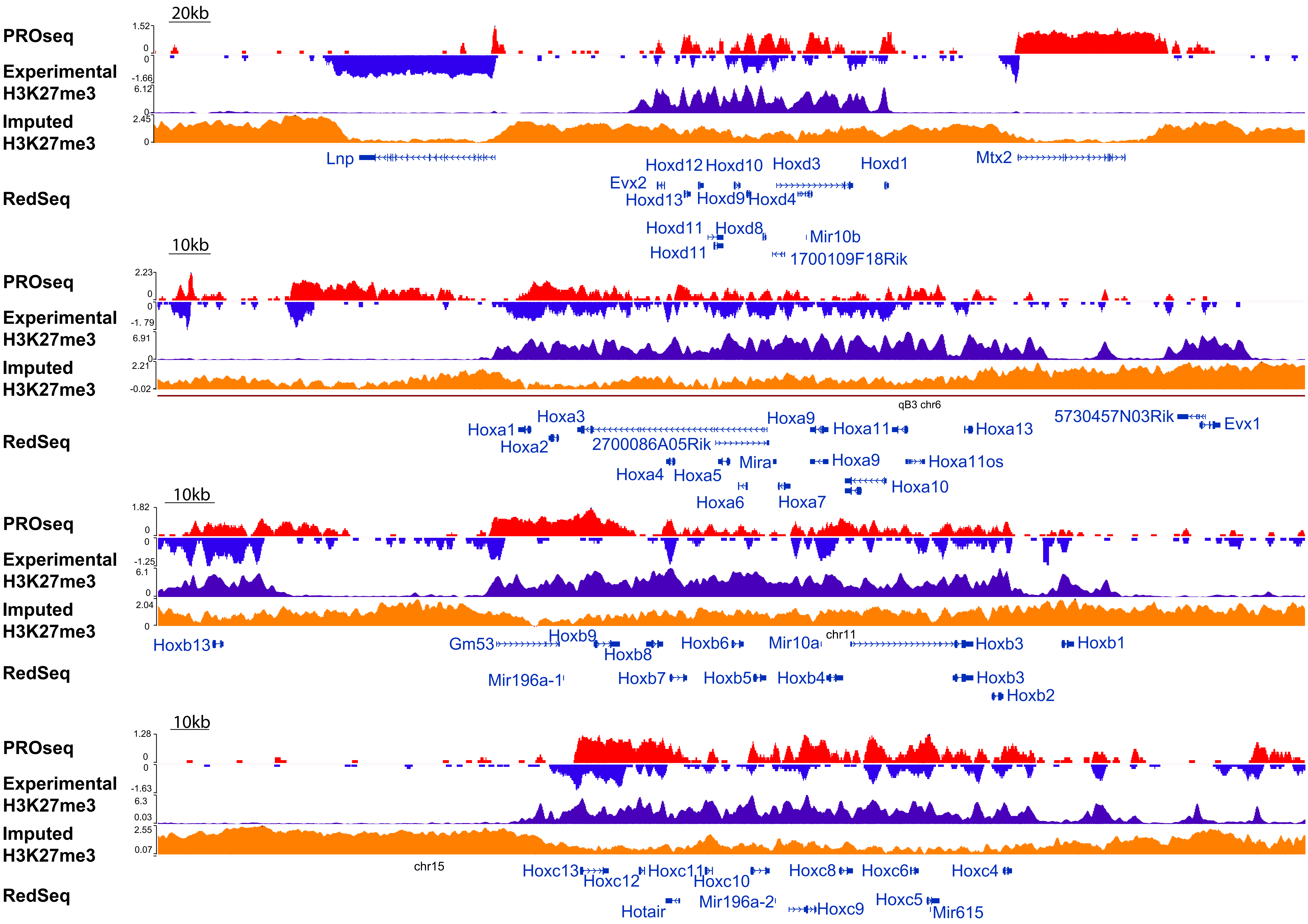


**Supplementary Figure 13. Comparison between experimental and imputed H3K27me3 ChIP-seq at Hox gene clusters.** Genome-browser compares experimental and predicted H3K27me3 signals at all four Hox gene clusters in relation to PROseq signal.

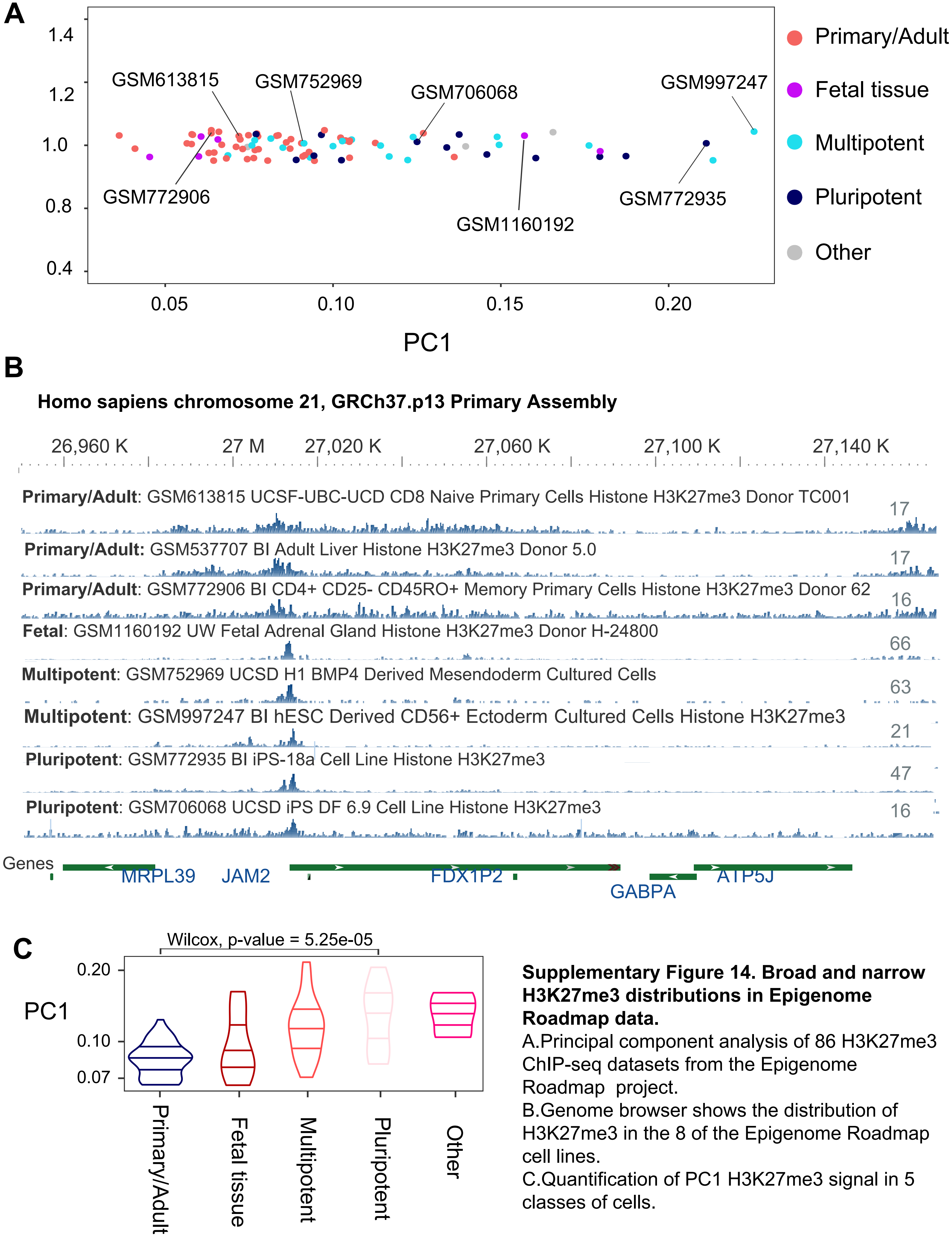


**Supplementary Figure 14. Broad and narrow H3K27me3 distributions in Epigenome Roadmap data.**
A.Principal component analysis of 86 H3K27me3 ChIP-seq datasets from the Epigenome Roadmap project.
B.Genome browser shows the distribution of H3K27me3 in the 8 of the Epigenome Roadmap cell lines.
C.Quantification of PC1 H3K27me3 signal in 5 classes of cells.

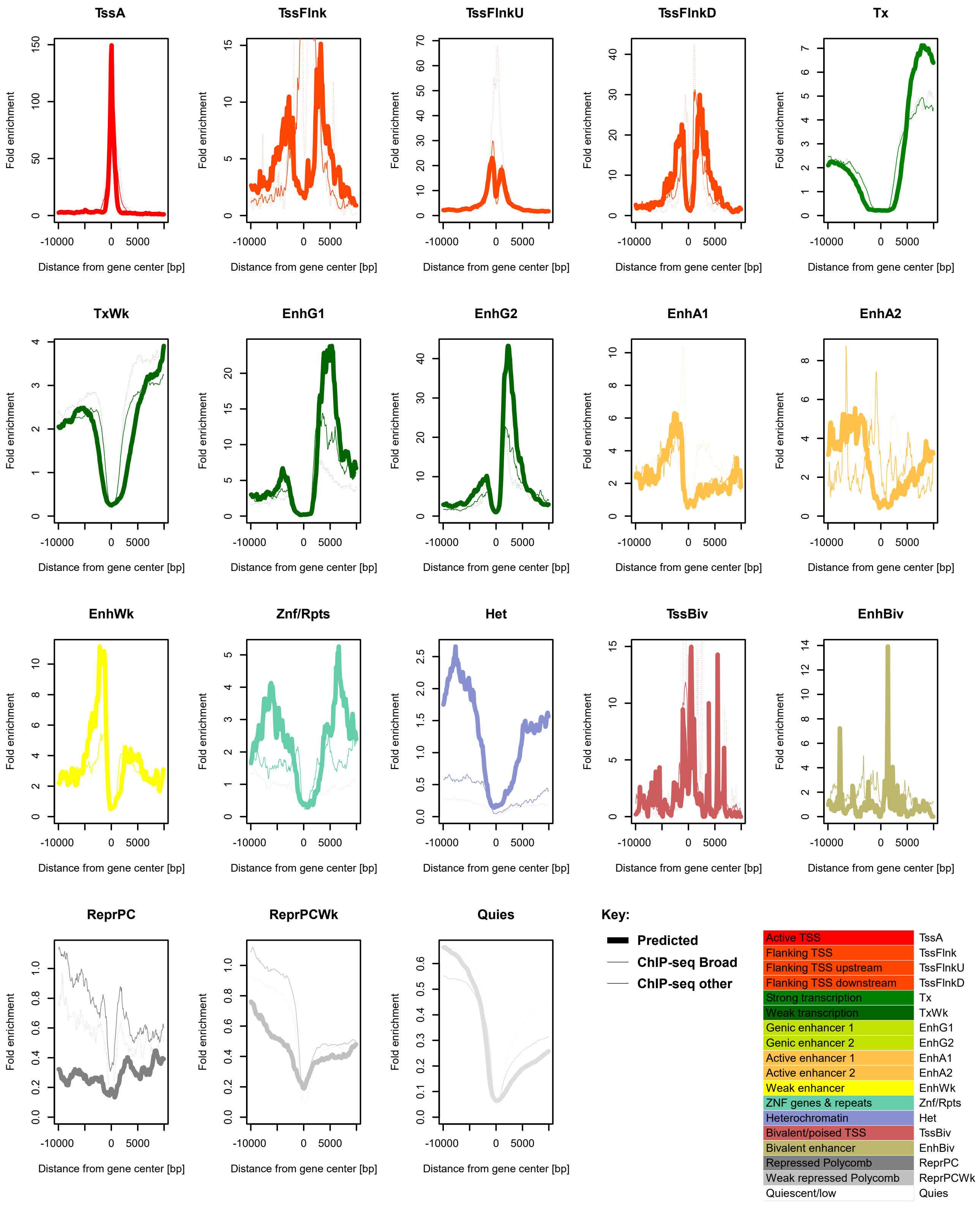


**Supplementary Figure 15. ChromHMM states near annotated transcription start sites.**
Enrichment of 18 chromatin states near RefSeq annotated transcription start sites for histone abundance predicted by dHIT (thick solid line), ChIP-seq from Broad (thin solid line), or using an alternative source of ChIP-seq data (thin dashed line).

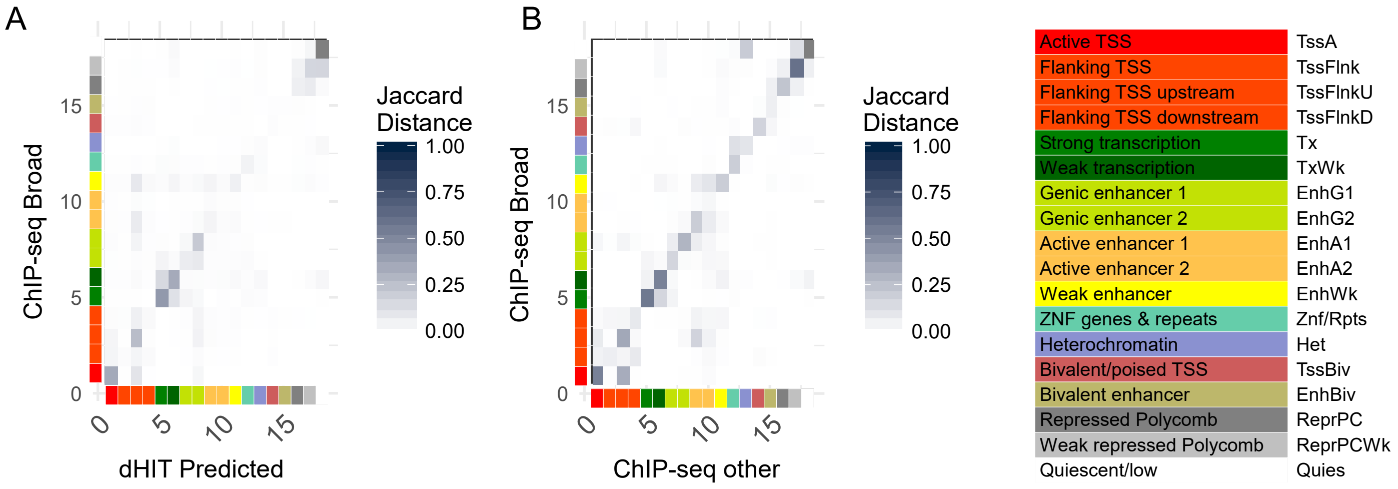


**Supplementary Figure 16. Confusion matrix comparing ChromHMM based on to dHIT.**
Confusion matrix shows the Jaccard distance between dHIT and ChIP-seq data in 18 chromatin states (A) or between two separate sources of ChIP-seq data (B). Color scales are shown beside the plot, and are identical between panels (A) and (B).

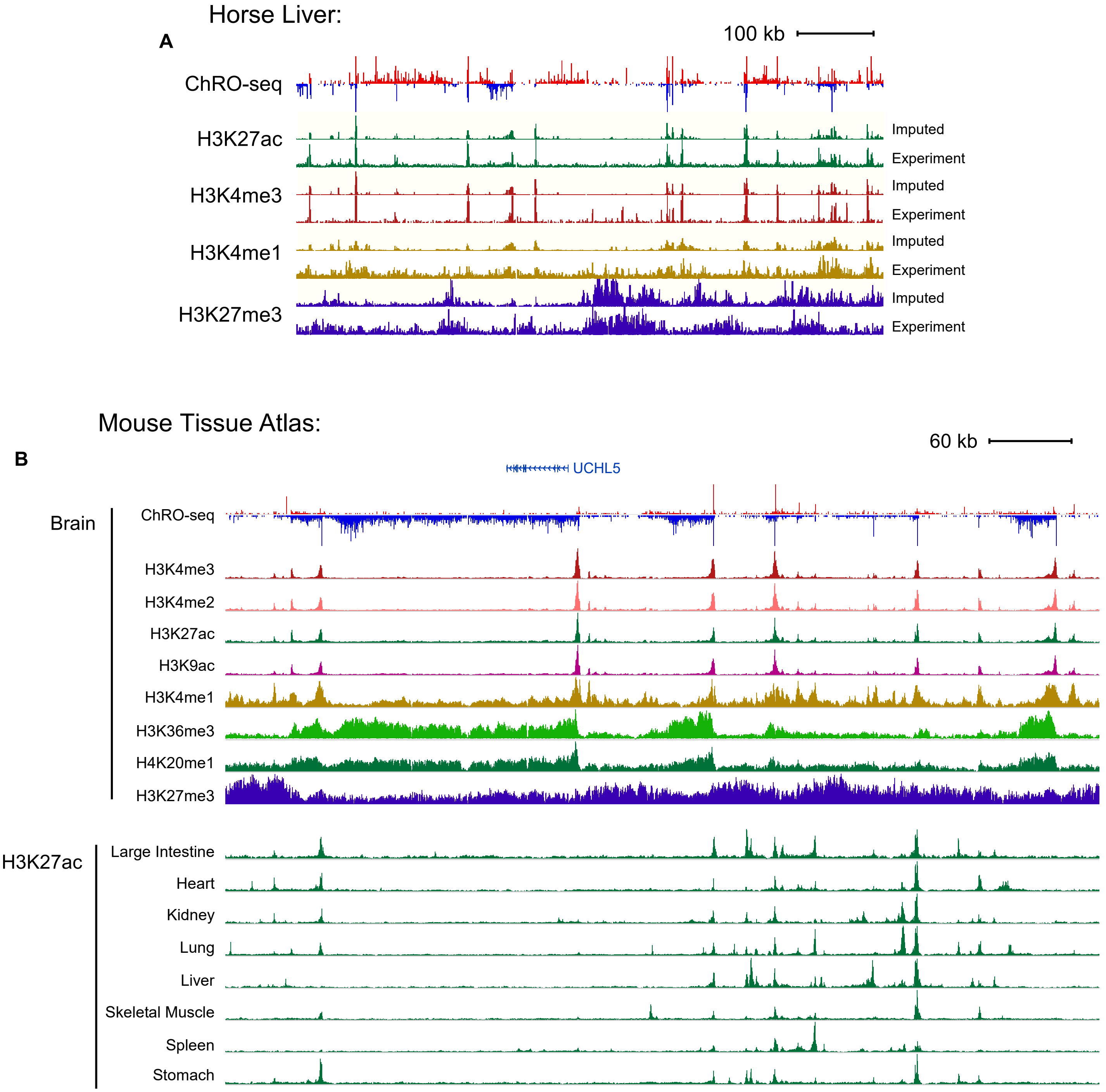


**Supplementary Figure 17. Chromatin marks in equine liver and nine murine tissues.**
(A) Genome browser shows the distribution of transcription, H3K27ac, H3K4me3, H3K4me1 and H3K27me3 in equine liver. (B) Genome browser shows the distribution of eight histone marks in mouse brain (top) and H3K27ac across nine murine tissues (buttom).

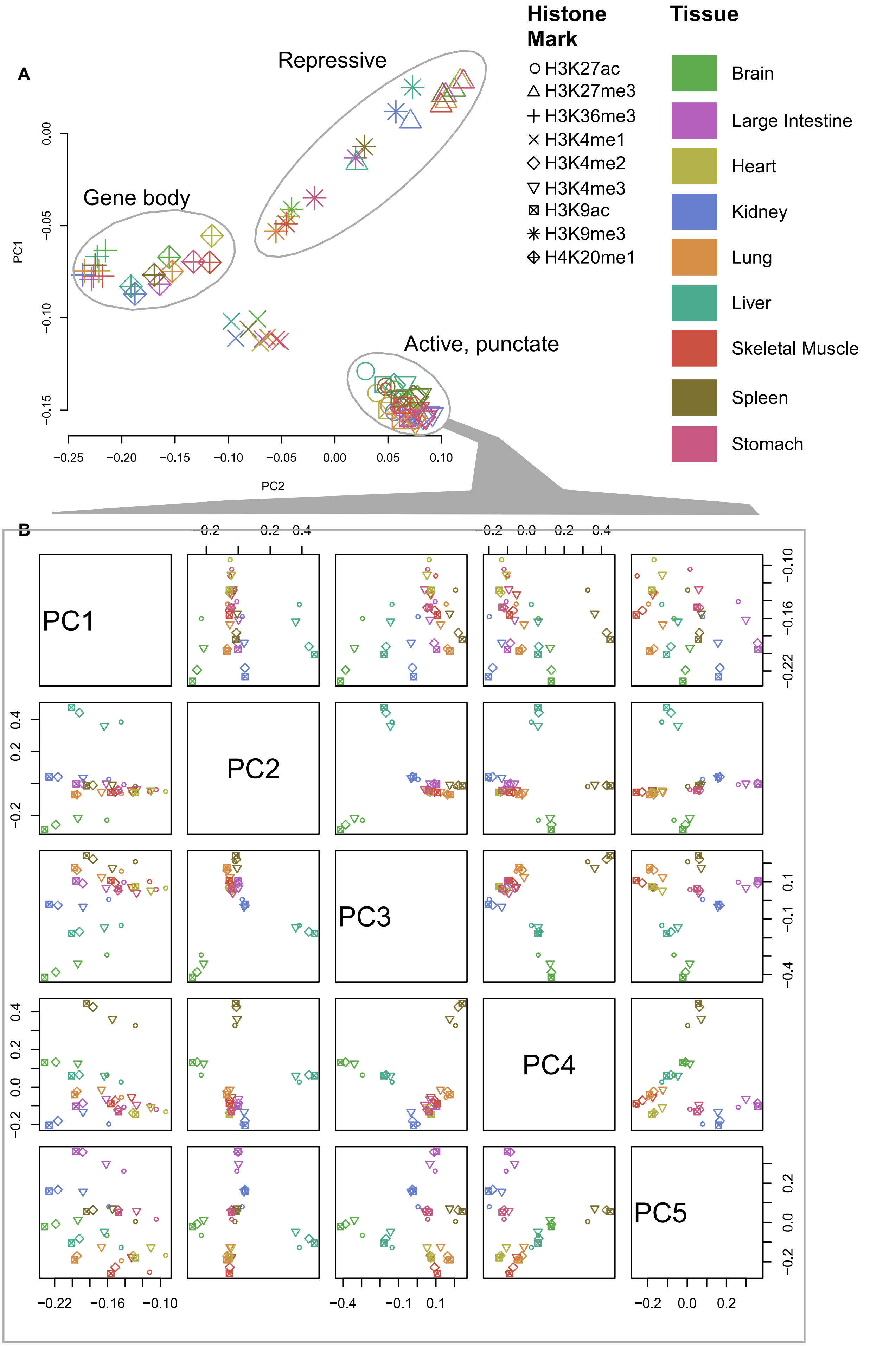


**Supplementary Figure 18. Summary of nine histone modifications in nine murine tissues.**
(A) PCA shows the first two principal components of nine histone modifications in nine murine tissues (81 total datasets) in 100 bp bins on mm10 chr1. (B) PCA of active, punctate marks (H3K4me3, H3K4me2, H3K9ac, and H3K27ac) shows that active punctate marks cluster by tissue.

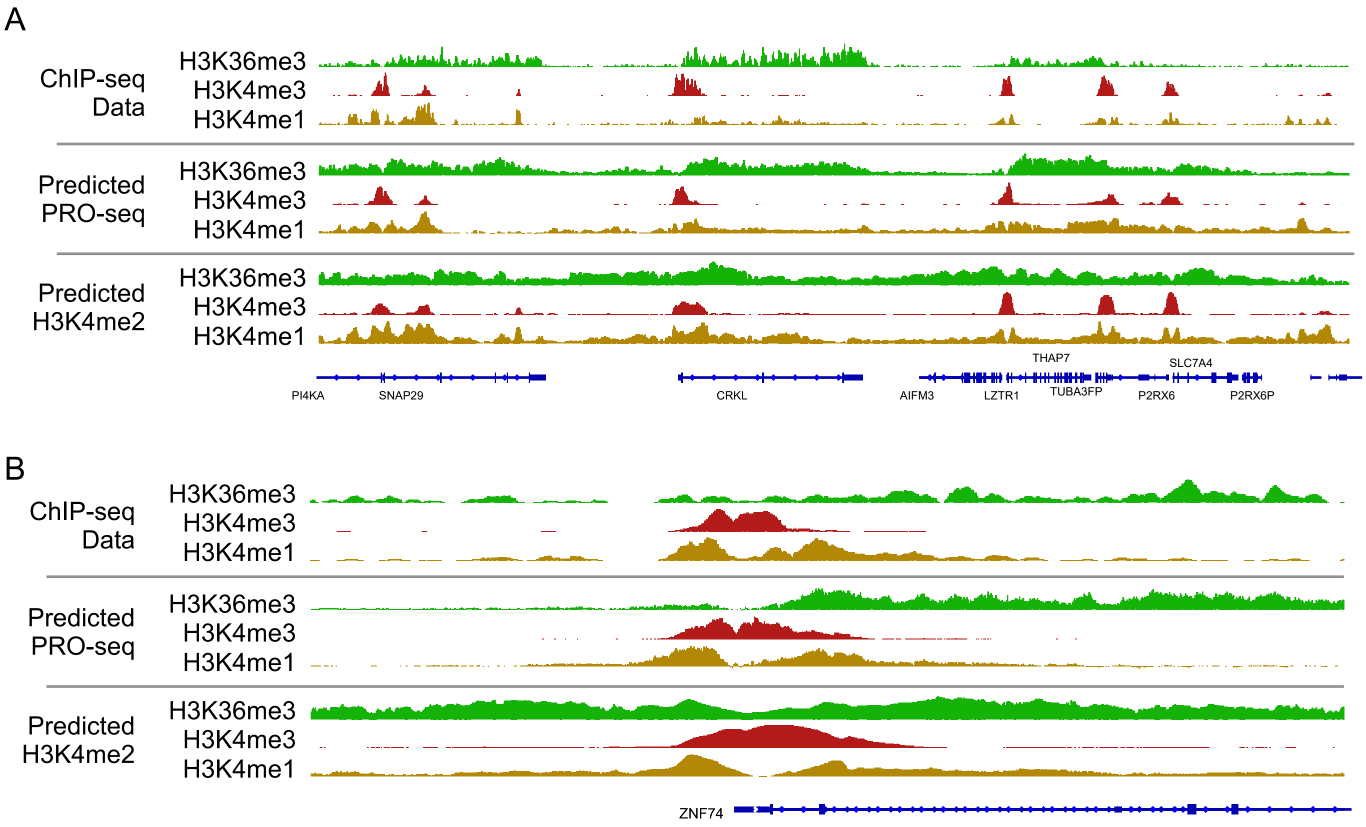


**Supplementary Figure 19. Chromatin mark abundance imputed using different marks as input.**
Genome browser shows the distribution of H3K36me3, H3K4me3, and H3K4me1 observed using ChIP-seq experiments or predicted using either PRO-seq or H3K4me2. Data is shown in two loci covering several transcribed genes (A) and near the transcription start site of ZNF74 (B).

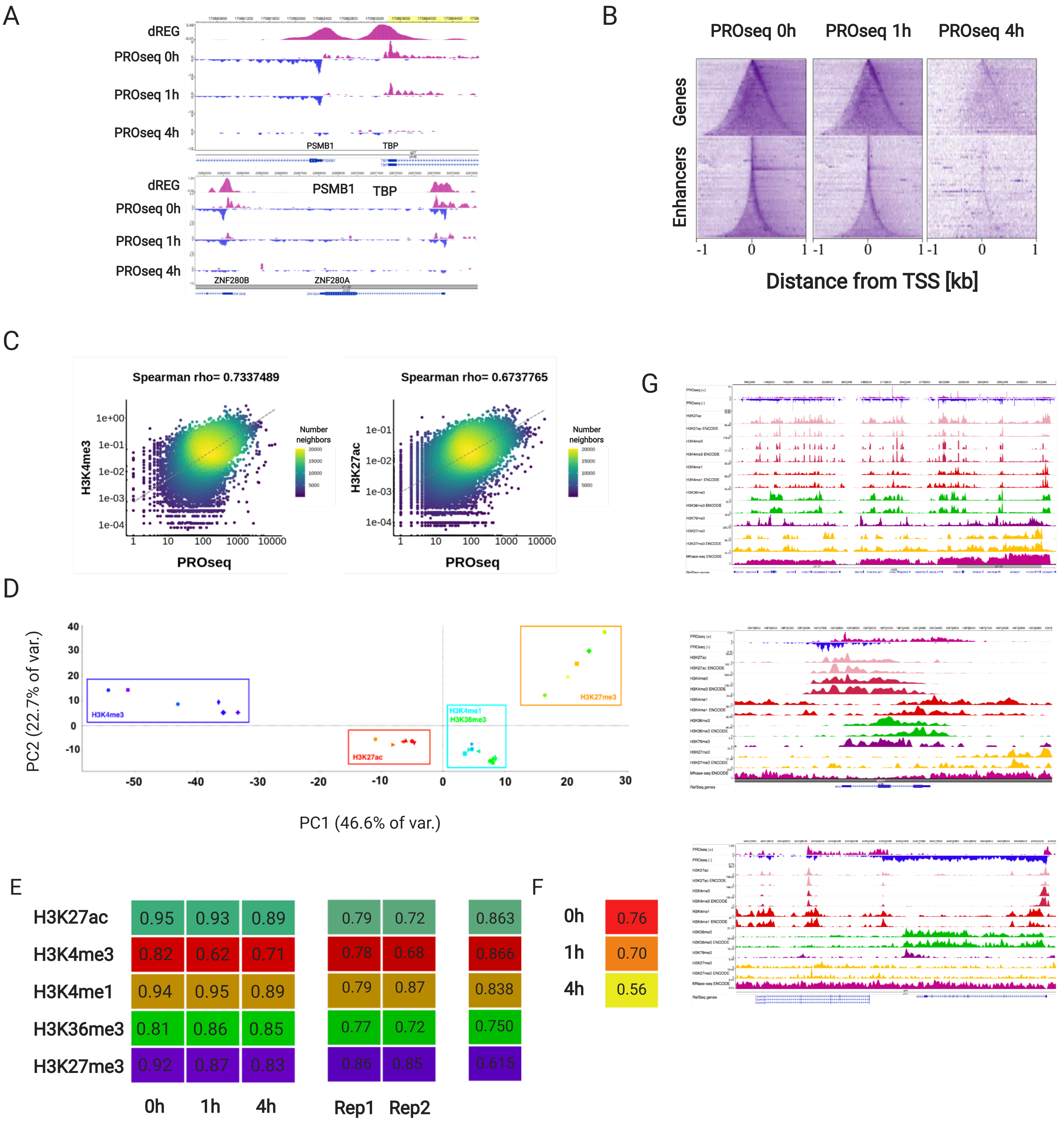


**Supplementary Figure 20. MNase ChIP-seq validation and analysis.**

(A) Genome-browser shows loss in transcription measured by PRO-seq after Trp treatment. Loss in PRO-seq signal at both enhancers and gene promoters.

(B) Heatmaps centered on transcription initiation domains show loss in transcription measured by PRO-seq after Trp treatment. Loss in PRO-seq signal at both enhancers and gene promoters.

(C) Correlations between PROseq 0h and H3K4me3 and H3K27ac at TSSs.

(D) Genome-wide, 10kb resolution PCA of all ChIP-seq samples.

(E) Spearman correlations between ChIP-seq replicates (left), each ChIP-seq replicate and ENCODE data (middle) genome-wide at 10kb resolution, and at ENCODE peaks between merged Reps and ENCODE.

(F) H3 Cut&Run 10kb resolution Spearman correlation between replicates.

(G) Browser shots comparing ChIPSeq with ENCODE data.

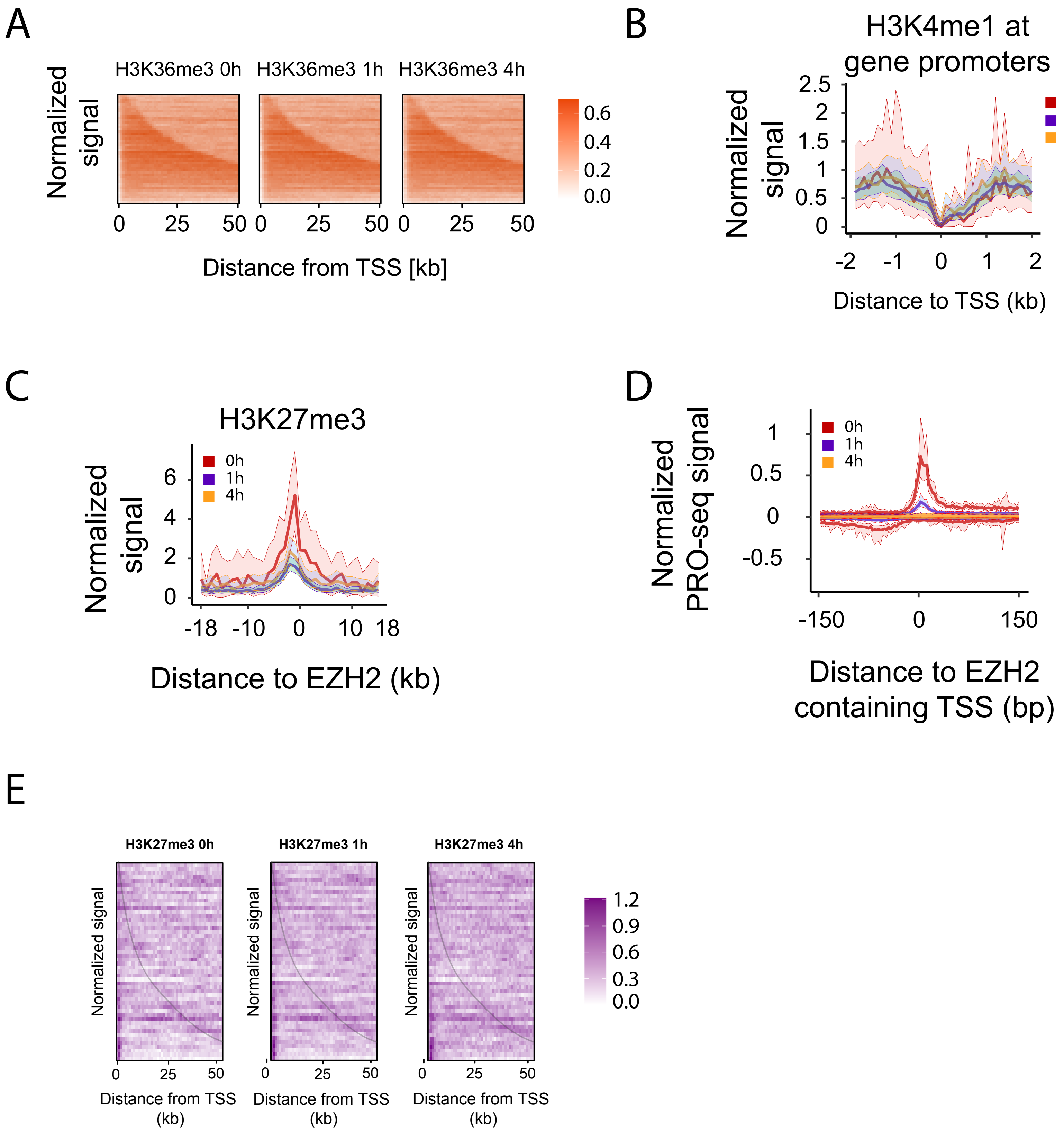


**Supplementary Figure 21. Changes in histone marks during Triptolide time course.**
(A) Heatmaps compare the level of H3K36me3 ChIP-seq after Triptolide inhibiti
(B) Meta plots show the H3K4me1 levels in a 4kb window centered on transcription start sites in K562 cel
(C) Meta plots show the level in H3K27me3 in a 40kb window centered in EZH2 binding sit
(D) Meta plots show transcription content of EZH2 binding sites during the Triptolide time course (E). Heatmaps disp
H3K27me3 signal within gene bodies during the Triptolide treatment. Genes are sorted by gene length.

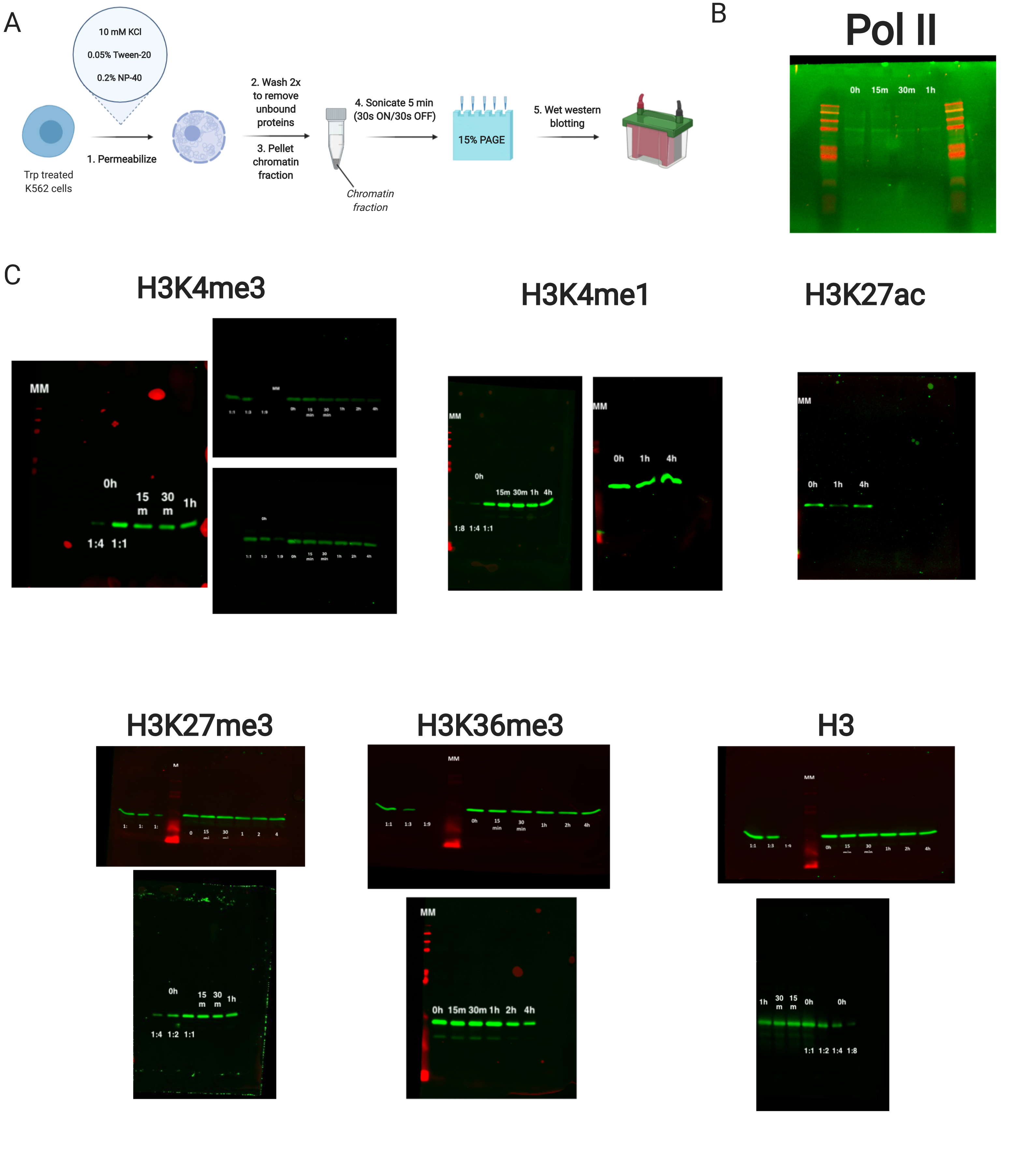


**Supplementary Figure 22. Western blots of histone marks and Pol II after Triptolide treatment.**
(A) Schematics of western blot experimental design.
(B-C) Each western blot depicts the abundance of chromatin bound histone mark or Pol II during the indicated Triptolide incubation time point. Each blot represents a different experiment. A dilution series of the untreated samples was used as standard curve to quantify changes in signal.

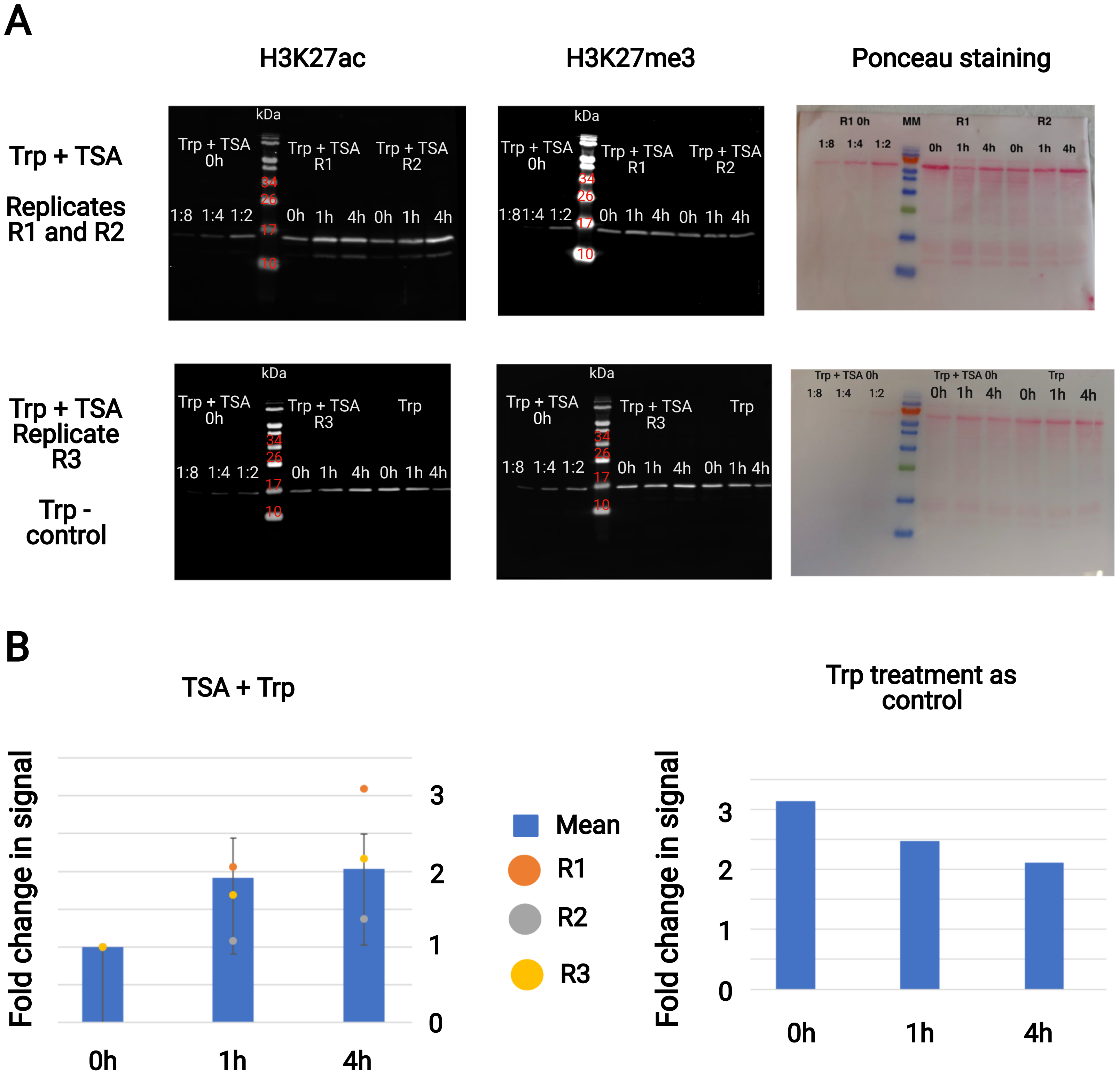


**Supplementary Figure 23. Western blots of H3K27ac after Triptolide (Trp) and Trichostatin A (TSA) treatment.**
**(A)** Each western blot depicts the abundance of chromatin bound H3K27ac or H3K27me3 during the indicated incubation time point of Triptolide, or Triptolide and Trichostatin dual treatment. Each blot represents a different experiment. A dilution series of the untreated samples was used as standard curve to quantify changes in signal. Ponceau staining of membranes imaged are also depicted as total protein loading control.
**(B)** Quantification of H3K27ac/H3K27me3 signals of the western blot in (A). H3K27me3 was used as loading control.

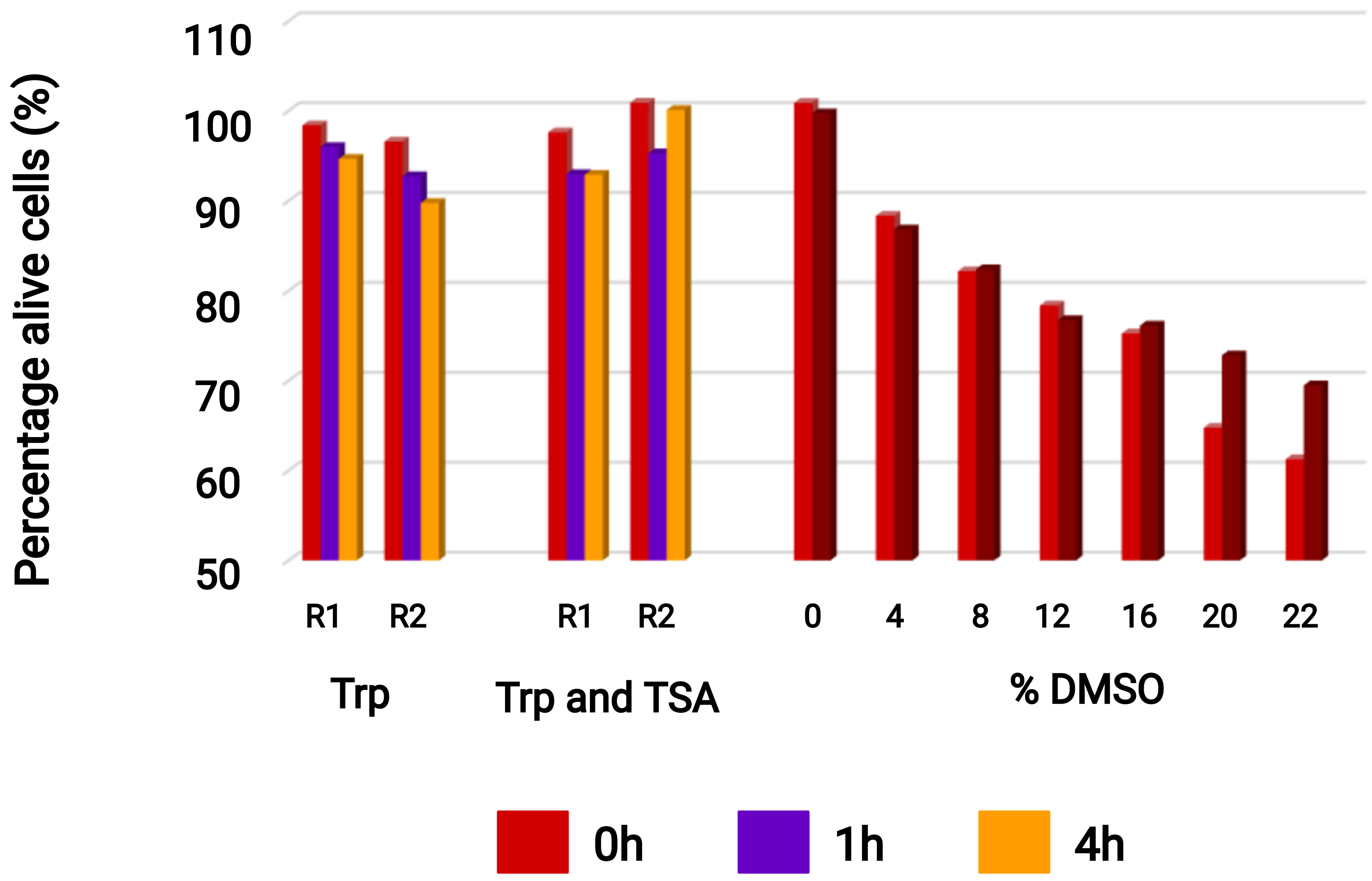


**Supplementary Figure 24. Cytotoxicity measurements for Triptolide, and Triptolide - Trichostatin A dual treatment in K562 cells**
Bar plots display absorbance quantified at **590**nm for AlmarBlue dye incubated with K**562** cells during Triptolide**,** or Triptolide and Trichostatin A treatments**.** Two technical replicates were averaged for each time point**.** R**1** and R**2** define separate biological replication of the experiment**.**

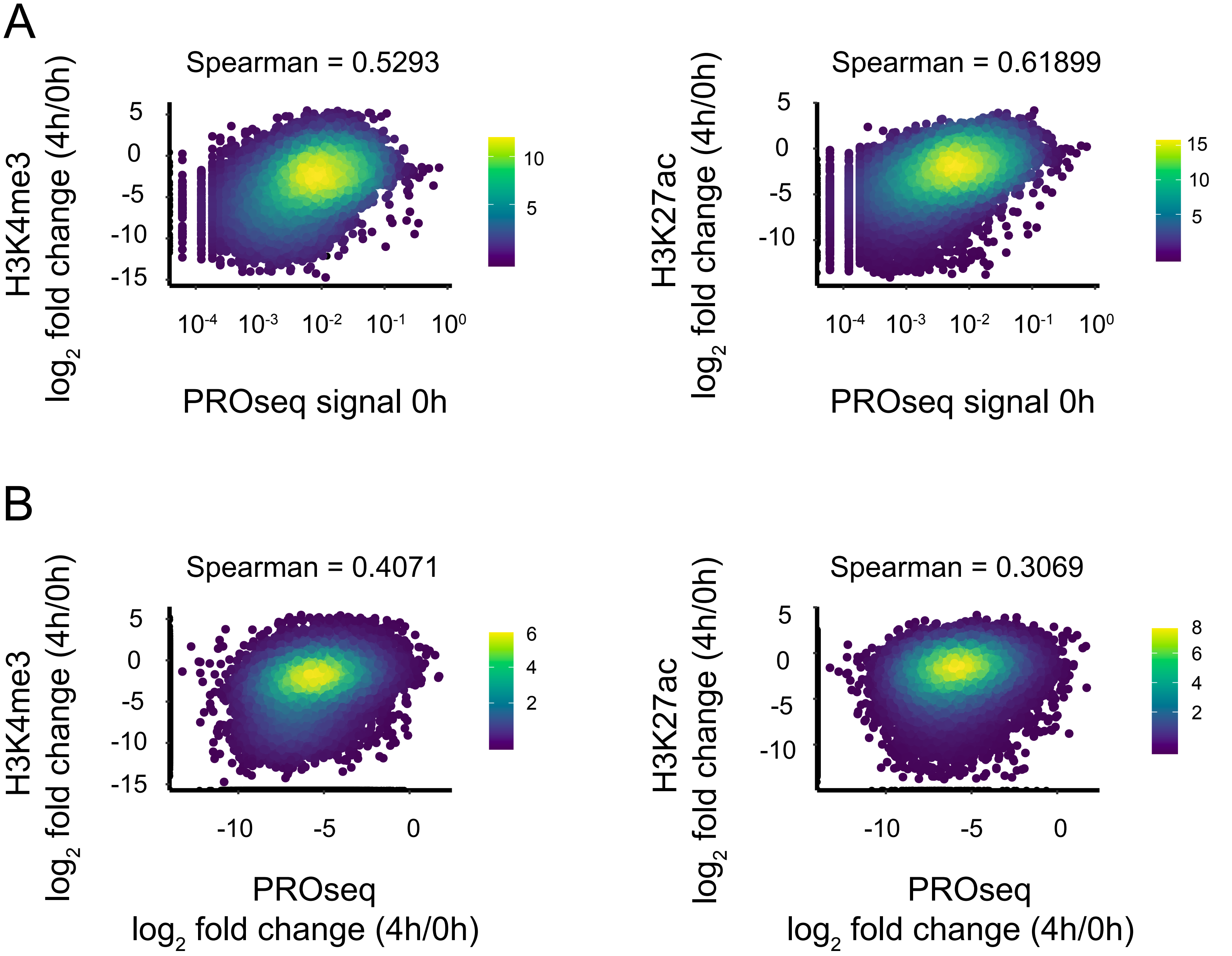


**Supplementary Figure 25. Comparison between the change in H3K4me3 and H3K27ac as a function of transcription.**(A-B) Scatter plots display the loss in H3K4me3 (left) and H3K27ac (right) as a function of Pol II transcription (A) or change in transcription (B). Changes in histone marks and transcription were calculated as log2 fold changes between 4h of Triptolide treatment and untreated cells. Plots show spearman rho correlations between conditions.

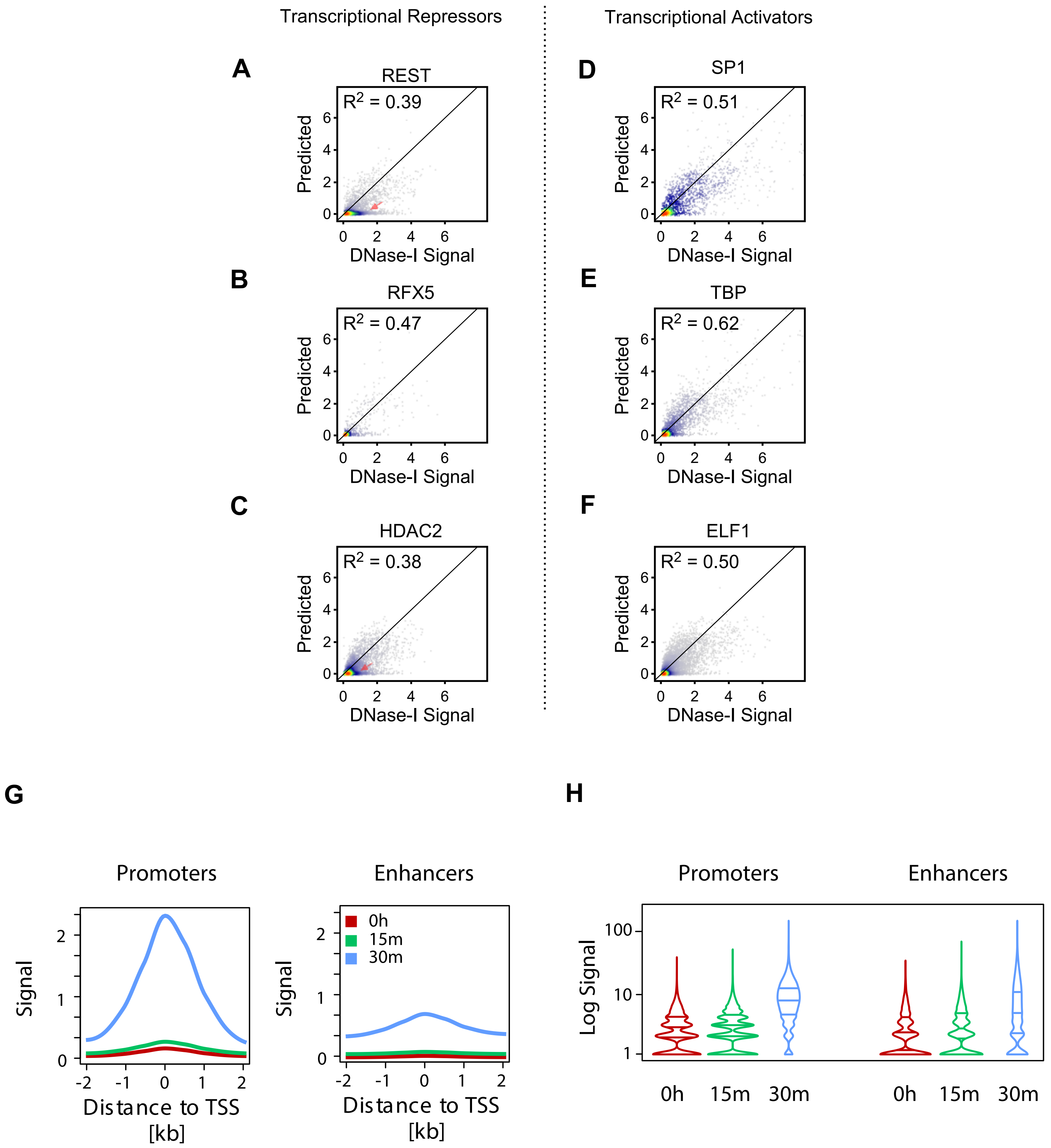


**Supplementary Figure 26. DNase-I hypersensitivity imputation near transcription repressors and activators.**
(A-F) Scatterplots show experimental DNase-I hypersensitivity (x-axis) as a function of predicted DNase-I hypersensitivity (y-axis) in 100 bp windows intersected with transcriptional repressors (A-C) or transcriptional activators (D-F).
(G-H) Meta (G) and Violin (H) plots display TBP CUT&RUN signal at gene promoters and enhancers in a short 30min Triptolide time course.
